# *N*-Alkyl Sulfamates as a New Class of nsP2 Cysteine Protease Inhibitors with Broad Spectrum Antialphaviral Activity

**DOI:** 10.1101/2025.06.30.662352

**Authors:** Anirban Ghoshal, John D. Sears, Mohammad Anwar Hossain, Edwin G. Tse, Stefanie Howell, Jane E. Burdick, Noah L. Morales, Sabian A. Martinez, Isabella Law, Zachary J. Streblow, Daniel N. Streblow, Rafael Miguez Couñago, Nathaniel J. Moorman, Mark T. Heise, Timothy M. Willson

## Abstract

The emergence of mosquito-borne alphaviruses that cause chronic arthritis or encephalitis underscores the urgent need for broad-spectrum antiviral therapeutics. The viral nsP2 cysteine protease, which is essential for alphavirus replication, is a promising antiviral target. Vinyl sulfone-based inhibitors, such as RA-2034, potently inhibit nsP2 protease but suffer from glutathione reactivity and species-dependent systemic clearance catalyzed by glutathione *S*-transferase. To address these liabilities, we explored alternative electrophilic warheads and identified reverse amide inhibitors bearing *N*-alkyl sulfamate warheads with improved biochemical and antiviral profiles. *N*-methyl sulfamate acetamide **5** emerged as a lead compound with potency against both New and Old World alphaviruses, low GSH reactivity, and high proteome-wide selectivity. Despite its promising antialphaviral activity, **5** exhibited rapid clearance due to hepatic glucuronidation. Structure–activity studies revealed modifications that improve metabolic stability while retaining antiviral activity. These findings introduce sulfamate acetamides as a new class of covalent nsP2 protease inhibitors and advance the discovery of direct acting pan-alphavirus drugs.

## Introduction

The increasing global incidence of mosquito-borne alphaviral infections,^1^ such as chikungunya virus (CHIKV)^2^ and Venezuelan equine encephalitis virus (VEEV),^3, 4^ highlights the urgent need for development of broad-spectrum, directly acting antiviral drugs (DAAs).^5^ Old World alphaviruses such as CHIKV cause chronic arthritis-like disease, while New World alphaviruses like VEEV result in an often fatal encephalitis.^6^ Despite advances in vaccine development,^7^ effective antiviral therapies for treatment of alphavirus diseases remain lacking. The nonstructural protein 2 protease (nsP2pro), a papain-like cysteine protease essential for alphaviral replication,^8^ has recently emerged as a promising small molecule DAA target.^9^

Among the most potent known inhibitors of nsP2pro is RA-2034 (**1**, Figure 1), a β-aminomethyl vinyl sulfone that functions as an irreversible covalent inhibitor of the cysteine protease.^10^ Vinyl sulfone **1** inhibits CHIKV nsP2pro with an IC₅₀ of 60 nM (*k*inact/*K*i of 6,400 M^-1^s^-1^) and demonstrates antiviral activity against multiple clinical isolates of infectious alphaviruses.^10^ However, while vinyl sulfones such as **1** demonstrate potent antiviral activity, they exhibit a bias towards the arthritogenic Old World alphaviruses over the encephalitogenic New World alphaviruses.^11^ In addition, vinyl sulfones like **1** suffer from poor pharmacokinetic properties^12, 13^ due to reactivity with glutathione (GSH), extensive metabolism by glutathione *S*-transferases (GSTs), and rapid peripheral clearance in rodents and primates.^14^

**Figure 1.**
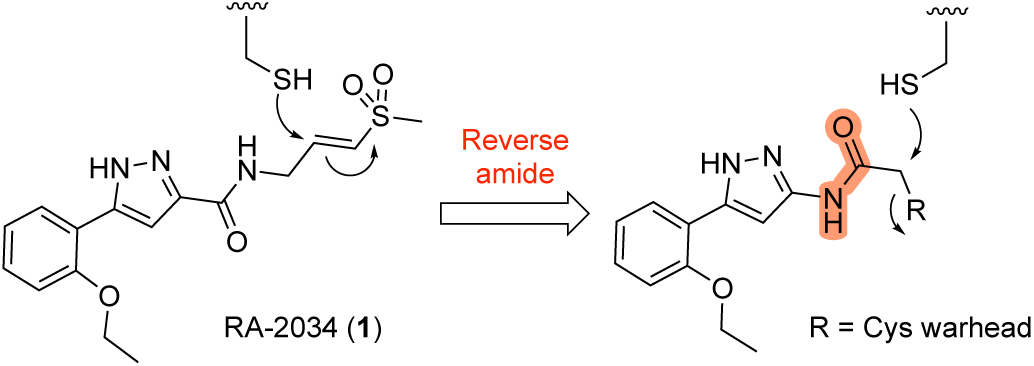
Design of nsP2 cysteine protease inhibitors

To address these limitations, we sought to identify a broad-spectrum nsP2pro inhibitor with improved GSH stability and reduced GST-mediated metabolism. Attempts to optimize the vinyl sulfone warhead of **1** failed to yield analogs with both potent protease inhibition and acceptable metabolic stability.^11, 14^ This result prompted a broader exploration of alternative electrophilic warheads with potential for targeting the nsP2 cysteine protease.

Here, we report the identification of a new class of nsP2pro inhibitors featuring reverse amide scaffolds bearing sulfamates as cysteine-targeting warheads (Figure 1). These sulfamate acetamides maintain potent anti-nsP2pro activity while displaying significantly lower reactivity toward GSH, addressing a major liability of vinyl sulfone-based inhibitors. Although the sulfamate acetamides are subject to rapid hepatic clearance due to UDP-glucuronosyltransferase (UGT)-mediated metabolism, they nonetheless represent a promising chemotype for the development of pan-alphavirus DAAs.

## Results and Discussion

### Alternative Warheads

We synthesized analogs of RA-2034 (**1**) in which the amide was reversed to allow the incorporation of several known cysteine warheads^15^ in analogs **2**–**5** (Figure 1 and Table 1). The analogs were tested for their ability to inhibit both encephalitogenic New World (VEEV) and arthritogenic Old World (CHIKV) alphaviruses using nLuc reporter assays that had been developed for the characterization of vinyl sulfone **1** and its analogs.^10, 11, 16^ Analogs were also tested for inhibition of the CHIKV nsP2pro enzyme using our established assay.^10, 11, 16^ Unfortunately, the corresponding VEEV nsP2pro assay was unavailable due to difficulty in expressing recombinant enzyme with robust biochemical activity, possibly due to the formation of an autoinhibited state as noted by others.^17, 18^ In antiviral assays, vinyl sulfone **1** showed a bias towards the arthritogenic Old World alphaviruses^19^ and was >10-fold more potent for inhibition of CHIKV compared to VEEV (Table 1). The reverse amide analogs were also tested for their stability to conjugation by GSH under conditions where vinyl sulfone **1** had a half-life of 2.3 h. The acrylamide **2** was inactive in the VEEV-nLuc and CHIKV-nLuc assays at 10 μM. Chloroacetamide **3** inhibited VEEV and CHIKV with EC50 = 100 nM on both viruses and was also a potent inhibitor the CHIKV nsP2pro enzyme. However, the chloroacetamide warhead in **3** was previously noted for its promiscuous reactivity with many proteins in cell lysates as shown by TAMRA labelling.^20, 21^ Methyl sulfonate **4** was less active than **1** in both the enzyme inhibition and antiviral assays. Sulfamate acetamides have been recently described as covalent warheads by London et al.^22^ When incorporated into the our reverse amide template, *N*-methyl sulfamate **5** demonstrated potent antiviral activity on VEEV and CHIKV with EC50 ∼ 100 nM on both viruses. *N*-methyl sulfamate **5** was also active in the CHIKV nsP2pro enzyme assay using our standard 30 min incubation. In the GSH stability assay the acrylamide **2**, chloroacetamide **3**, and methyl sulfonate **4** showed half-lives in the 2 h range, similar to vinyl sulfone **1**. Remarkably, *N*-methyl sulfamate **5** was much more stable to GSH with a half-life >12 h (Table 1).

**Table 1.** Antiviral Activity, nsP2 Protease Inhibition, and GSH Reactivity of 1–5.

| Compound | R | VEEV-nLuc<br>(pEC<sub>50</sub>)<sup>a</sup> | CHIKV-nLuc<br>(pEC<sub>50</sub>)<sup>a</sup> | CHIKV<br>nsP2pro<br>(pIC<sub>50</sub>)<sup>b</sup> | GSH<br>t<sub>1/2</sub> (h)<sup>c</sup> |
| --- | --- | --- | --- | --- | --- |
| RA-2034 (**1**) |  | 6.5 | 7.6 | 7.2 | 2.3 |
| **2** | CH=CH<sub>2</sub> | i.a. | i.a. | i.a. | 2.6 |
| **3** | Cl | 7.0 | 7.0 | 7.4 | 2.4 |
| **4** | OSO<sub>2</sub>Me | 5.7 | 6.3 | 5.6 | 1.7 |
| **5** | OSO<sub>2</sub>NHMe | 7.2 | 6.9 | 6.3 | 14 |

<sup>a</sup>pEC<sub>50</sub> calculated from an 8-point dose response curve. Values are the average of duplicate determinations. <sup>b</sup>pIC<sub>50</sub> calculated from at 10-point dose response curve. Values are the mean of triplicate determinations with variance <0.1. i.a. = inactive at 10 μM. <sup>c</sup>Half-life in 5 mM GSH at pH 7.4.

From these initial studies, *N*-methyl sulfamate **5** emerged as the most interesting covalent warhead. It demonstrated equivalent activity in the VEEV and CHIKV antiviral assays while having much lower GSH reactivity. *N*-methyl sulfamate **5** was isolated as a free flowing white solid with m.p. 173 °C that was stable when stored at room temperature for 6 months and was also stable in solution across a pH range from 3–12 with no evidence of degradation (Supplementary Figure S1). **5** has aqueous kinetic solubility of 70 µM and showed no evidence of aggregation up to 100 mM by dynamic light scattering (Supplementary Figure S2).

### Enzyme kinetics

*N*-methyl sulfamate **5** was a potent inhibitor of the CHIKV nsP2 protease with IC50 = 0.5 µM (Table 1). To demonstrate covalent enzyme inhibition, time dependent inactivation of CHIKV nsP2pro was measured at multiple concentrations of **5** (Figure 2). Analysis of the kinetic inactivation data produced a *k*inact/*K*i ratio of 2030 M^−1^s^−1^ (Figure 2), approximately one-third the value determined for vinyl sulfone **1**.^20, 21^ Thus, *N*-methyl sulfamate **5** was a covalent cysteine protease inhibitor, but was less efficient at inactivating the enzyme than the vinyl sulfone **1**.

**Figure 2.**
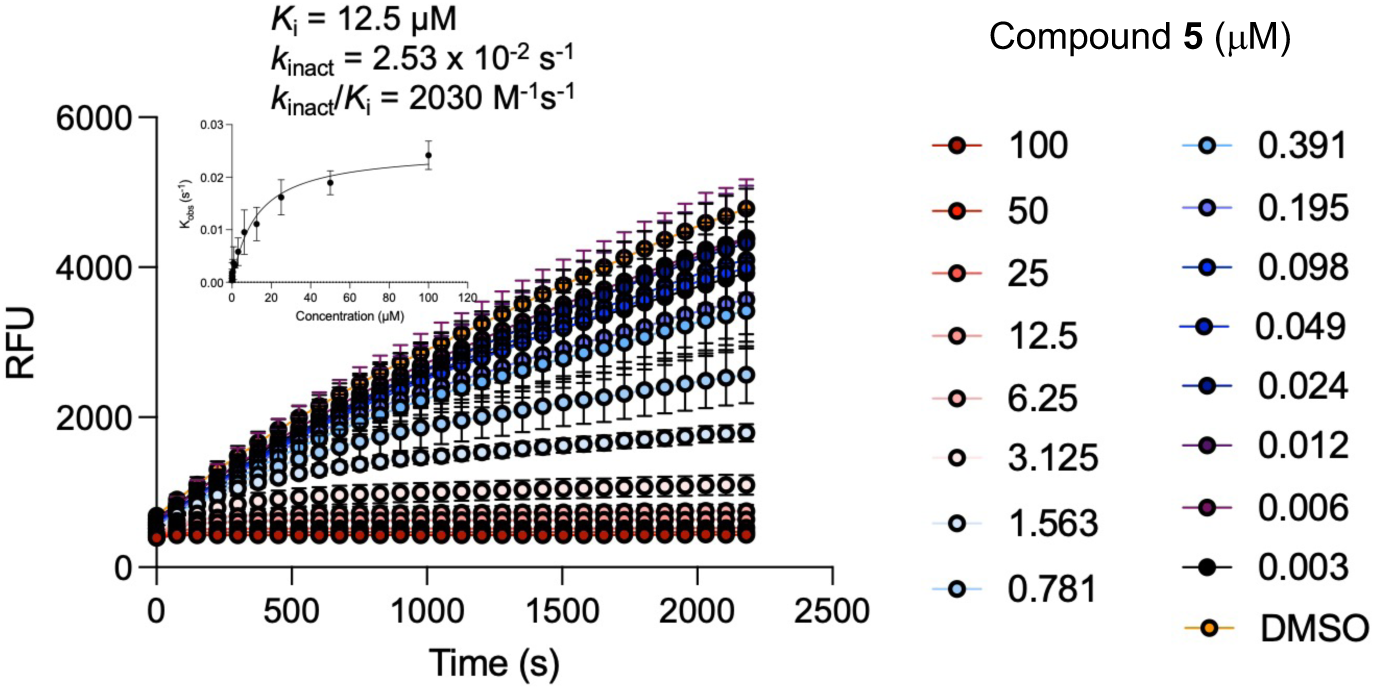
Time-dependent inhibition of CHIKV nsP2pro by *N*-methyl sulfamate **5**. 2-fold serial dilutions of **5** were used to determine the *kinact/Ki* value.

### Cysteine Protease Selectivity

*N*-methyl sulfamate **5** was screened against three human cysteine proteases and one viral cysteine protease at 10 µM in duplicate (Figure 3 and Supplementary Table S2). Compared to control inhibitors, **5** demonstrated no significant inhibition on cathepsin L, ubiquitin specific peptidase USP7, cysteine protease UCHL1, and papain-like protease SARS-CoV-2 Plpro. The data indicated that *N*-methyl sulfamate **5** was at least 30-fold selective for CHIKV nsP2pro over these other cysteine proteases.

**Figure 3.**
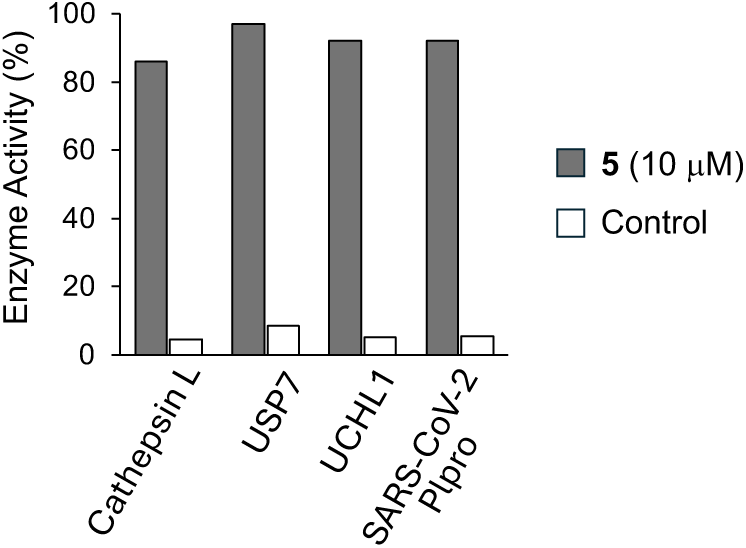
Percent activity of human and viral cysteine proteases in the presence of *N*-methyl sulfamate **5** (10 µM). The following were used as controls: E-64 (500 nM) for cathepsin L; ubiquitin-aldehyde (10 nM) for USP7 and UCHL1; GC376 (100 nM) for SARS-CoV-2 Plpro. Values are the average of duplicate determinations.

### Proteome-wide selectivity

The selectivity of the *N*-methyl sulfamate warhead was evaluated by fluorescence-based chemoproteomics in human HEK293 cell lysates (Figure 4). A clickable analog of *N*- methyl sulfamate **5** containing an alkyne handle on the phenyl substituent was synthesized: **NMS** (Figure 4A). Several control compounds were available from prior studies:^21^ clickable chloroacetamide **CA**, non-clickable control **VSC**, and fluorescent probe **TVS** containing a TAMRA dye (Figure 4A). Human HEK293 cell lysates were incubated with **NMS** or **CA** (10 µM, 1 h), followed by copper-catalyzed click reaction with TAMRA-N3 fluorophore, separation of proteins on a denaturing polyacrylamide gel, and fluorescence imaging. Fluorescent **TVS** that had been incubated with full length nsP2 was run on the gel as a control and showed the expected band at 42.9 kDa (Figure 4B, lane P). Chloroacetamide **CA** showed extensive labelling of proteins from the cell lysate, indicating its promiscuous activity (Figure 4B, lane C) as had been noted previously.^20, 21^ By contrast, no labeling of the HEK293 cell lysate was seen with **NMS** (Figure 4B, lane A). Control experiments using cell lysates spiked with purified nsP2 showed strong labelling by **CA** (Figure 4B, lane D) and weak labeling by **NMS** (Figure 4B, lane B). Surprisingly, preincubation with **VSC** lacking an alkyne was ineffective at blocking the labeling of nsP2 by **NMS** and **CA** (Figure 4B, lanes B1 and D1, respectively) suggesting that either higher concentrations or longer pre-incubation may be required for **VSC** to fully compete for labeling of nsP2 protein that was spiked into the lysate. These TAMRA labeling experiments demonstrated *N*-methyl sulfamate **5** was not a promiscuous covalent ligand, and that the combination of its sulfamate warhead with the pyrazole core generated a selective covalent nsP2pro inhibitor. Notably, **5** did not display cellular toxicity as measured by Cell Titer-Glo at concentrations up to 10 µM in A549ACE2 or HEK293 after 48 h exposure (Figure S4).

**Figure 4.**
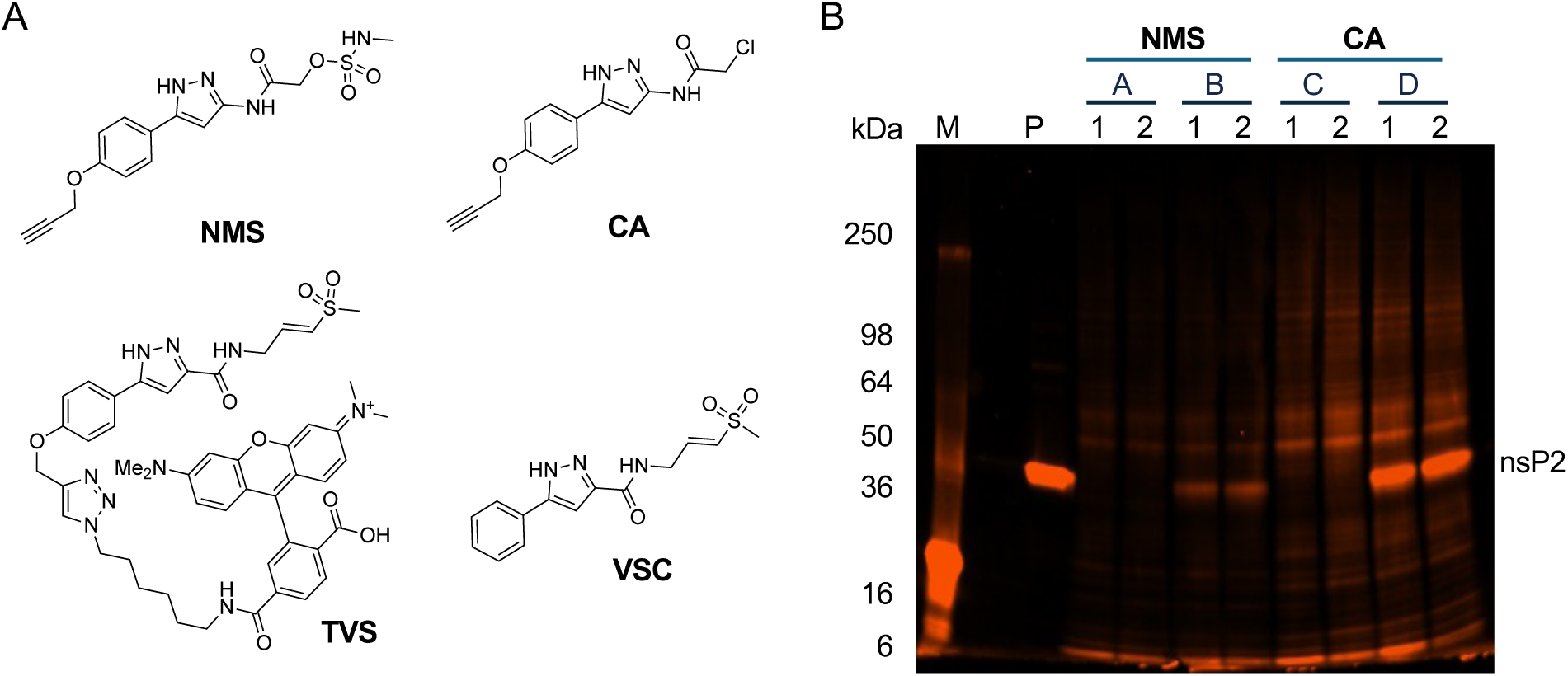
TAMRA fluorescence chemoproteomics. (A) Chemical structures of chemoproteomics probes: *N*-methyl sulfamate (**NMS**), vinyl sulfone control (**VSC**), chloroacetamide (**CA**), and TAMRA vinyl sulfone (**TVS**). (B) Fluorescent imaging of SDS-PAGE following covalent labeling of purified nsP2 (lane P) by **TVS** or human cell lysates (lanes A–D) by **NMS** or **CA**. In lanes 2, **NMS** or **CA** were incubated with cell lysates prior to click reaction to append the TAMRA fluorophore. In lanes 1, lysates were treated with **VSC** for 30 min prior to addition of **NMS** or **CA**. All ligands were used at 10 µM. Lanes A contain HEK293 cell lysates (140 µg total protein each), whereas lanes B and D equivalent amounts of cell lysates were supplemented with purified CHIKV nsP2 (16 µg). M.Wt markers are indicated. The nsP2 band with calculated M.Wt. 42.9 kDa is marked while accounting for the slight curvature of the gel. An uncropped copy of the gel with membrane edges visible is shown in Supplementary Figure SX.

### Alphavirus activity

*N*-methyl sulfamate **5** inhibited the replication of both VEEV-nLuc and CHIKV-nLuc reporter viruses with EC50 ∼100 nM (Table 1). To determine if the antiviral activity measured in the reporter assays translated to inhibition of infectious alphaviruses, **5** was tested against CHIKV (181/25),^23^ MAYV (BeAr),^24^ and VEEV (TC83).^25^ CHIKV and MAYV are representatives of the arthritogenic clade of Old World alphaviruses, while VEEV is an encephalitogenic New World alphavirus.^26^ Following infection of NDHF cells for 24 h, viral titer was determined by plaque assay in Vero cells (Table 2). Vinyl sulfone **1** was effective at reducing viral titer on the three alphaviruses. However, as was noted in the nLuc reporter assays, **1** was more potent on CHIKV compared to VEEV. Antiviral activity of **1** on MAYV mirrored the activity on CHIKV. Thus, vinyl sulfone **1** was more effective on two arthritogenic alphaviruses than a representative encephalitogenic alphavirus. In contrast, *N*-methyl sulfamate **5** was potent and highly effective at reducing viral titer on all three alphaviruses. The equipotent antiviral activity of **5** on VEEV and CHIKV matched its profile in the nLuc reporter assays. *N*-methyl sulfamate **5** was also >20-fold more potent and >7-fold more effective than vinyl sulfone **1** at reducing viral titer in the VEEV infected cells (Table 2). Together, the virus replication and the titer reduction assays demonstrated that *N*- methyl sulfamate **5** had an improved pan-antialphavirus profile compared to the vinyl sulfone **1**.

**Table 2.** Antialphaviral breadth of 1 and 5.

| Compound | CHIKV (181/25) |  | MAYV (BeAr) |  | VEEV (TC 83) |  |
| --- | --- | --- | --- | --- | --- | --- |
|  | VTR (x 10<sup>4</sup>)<sup>a</sup> | pEC<sub>90</sub> | VTR (x 10<sup>4</sup>)<sup>a</sup> | pEC<sub>90</sub> | VTR (x 10<sup>4</sup>)<sup>a</sup> | pEC<sub>90</sub> |
| **1** | 2.1 | 7.2 | 12 | 7.1 | 46 | 6.5 |
| **5** | 2.4 | 7.5 | 28 | 7.5 | 350 | 7.9 |

<sup>a</sup>VTR, viral titer reduction at 10 mM compared to DMSO control measured by plaque assay. IC<sub>90</sub> calculated from 11-point dose response curves. All data n = 1 in triplicate.

Importantly, even though *N*-methyl sulfamate **5** was less efficient than vinyl sulfone **1** at inactivating the nsP2pro enzyme (Figure 2), it was just as effective at reducing viral titer in CHIKV infected cells and more effective in VEEV infected cells (Table 2).

### Hepatocyte Metabolism

Vinyl sulfone **1** was rapidly metabolized in mice by GST-catalyzed GSH conjugation.^14^ *N*-methyl sulfamate **5** was more stable to uncatalyzed GSH conjugation than **1** (Table 1). To determine if **5** also had improved metabolic stability it was incubated in primary mouse and human hepatocytes (Table 3). Unfortunately, when **5** was incubated with primary mouse hepatocytes, it was rapidly metabolized with a half-life of only 2.6 min and an intrinsic clearance of 544 µL/min/10^6^ cells. Although, the metabolism of **5** was slower in primary human hepatocytes, it still showed a moderately high rate of clearance. The rapid clearance of **5** in hepatocytes indicated that it would subject to extensive first pass metabolism if dosed *in vivo*. Therefore, we decided to determine the basis of the hepatic clearance and address the metabolic instability before pursuing *in vivo* pharmacokinetic studies. To identify the pathway of metabolic clearance, co-dosing studies in primary mouse hepatocytes were performed with either 1- aminobenzotriazole (1-ABT), an irreversible inhibitor of P450 enzymes, or ethacrynic acid (EA), an irreversible inhibitor of GST enzymes.^27^ In the presence of 1-ABT the metabolism of **5** was decreased only ∼3-fold based on the half-life and intrinsic clearance (Table 3). EA had no effect on the metabolism of **5**. These results indicated that P450 oxidation was only a minor contributor to the metabolism of **5** and that GST-catalyzed GSH conjugation was not contributing to rapid clearance in mouse hepatocytes. Thus, the metabolic pathway responsible for rapid metabolism of **5** in hepatocytes remained unknown.

**Table 3.** ***In Vitro* Metabolism of 5 in Hepatocytes**

| | t<sub>1/2</sub><br>(min) | CL<sub>int</sub><br>($\mu\text{L}/\text{min}/10^6$ cells) |
| --- | --- | --- |
| Mouse hepatocytes | 2.6 | 540 |
| Human hepatocytes | 23 | 61 |
| Mouse hepatocytes + 1-ABT<sup>a</sup> | 8.9 | 160 |
| Mouse hepatocytes + EA<sup>b</sup> | 2.7 | 520 |

<sup>a</sup>P450 inhibitor, 1 mM. <sup>b</sup>GST inhibitor, 0.2 mM.

### Metabolite Identification

To identify the specific pathways of metabolism of **5**, it was incubated with cryopreserved male CD-1 mouse hepatocytes in the presence of 1-ABT (1 mM) to suppress any P450- mediated oxidation. Samples were collected after 0, 15, 30 min for analysis by LC-MS/MS, which resulting in the identification of 13 metabolites of **5** (Table 4 and Figure 5). The structures of metabolites M1–13 were assigned by analysis of MS fragmentations and quantitation of their relative abundance was determined from the UV peak area in the 30 min sample (Table 4 and Figure S5).

**Figure 5.**
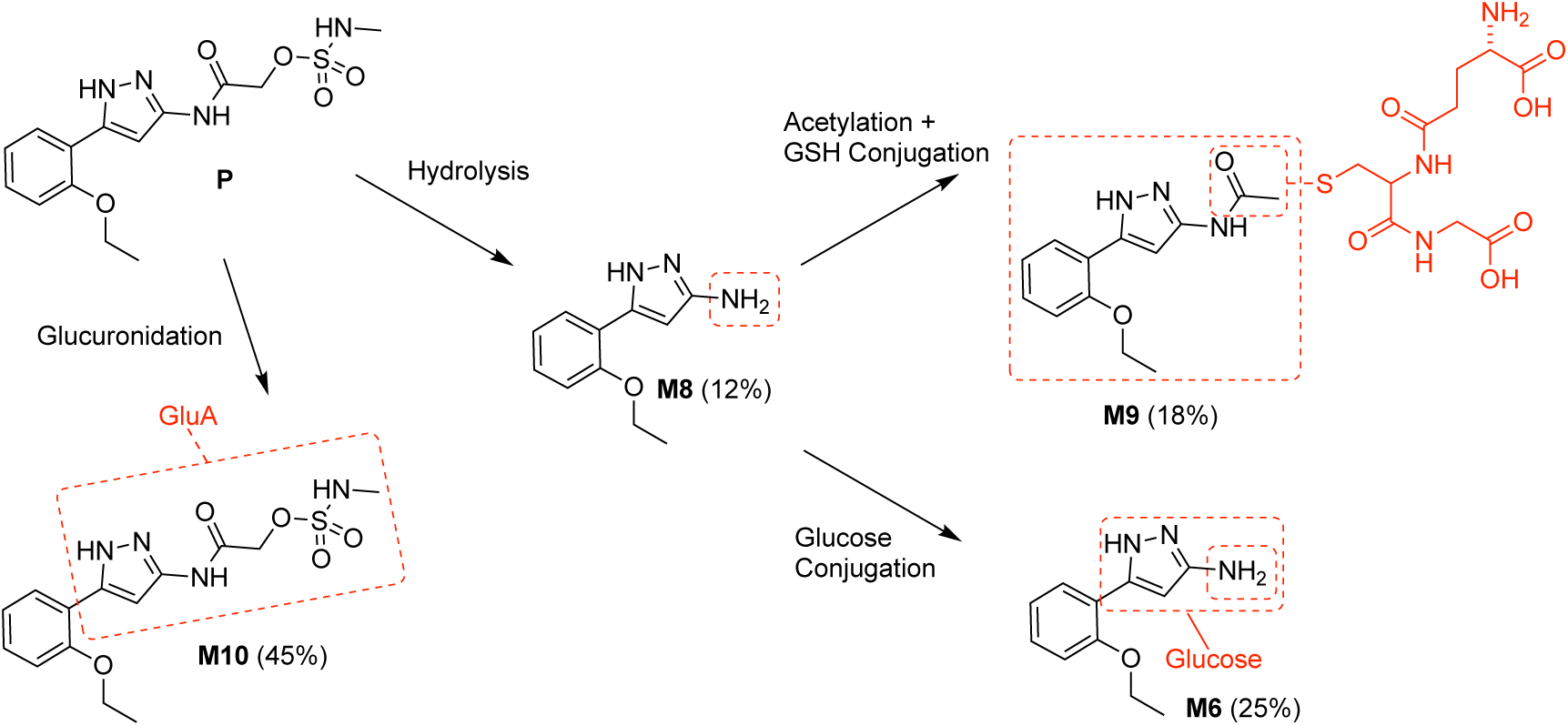
Major metabolites of **5** (P) identified in mouse hepatocytes.

**Table 4.**
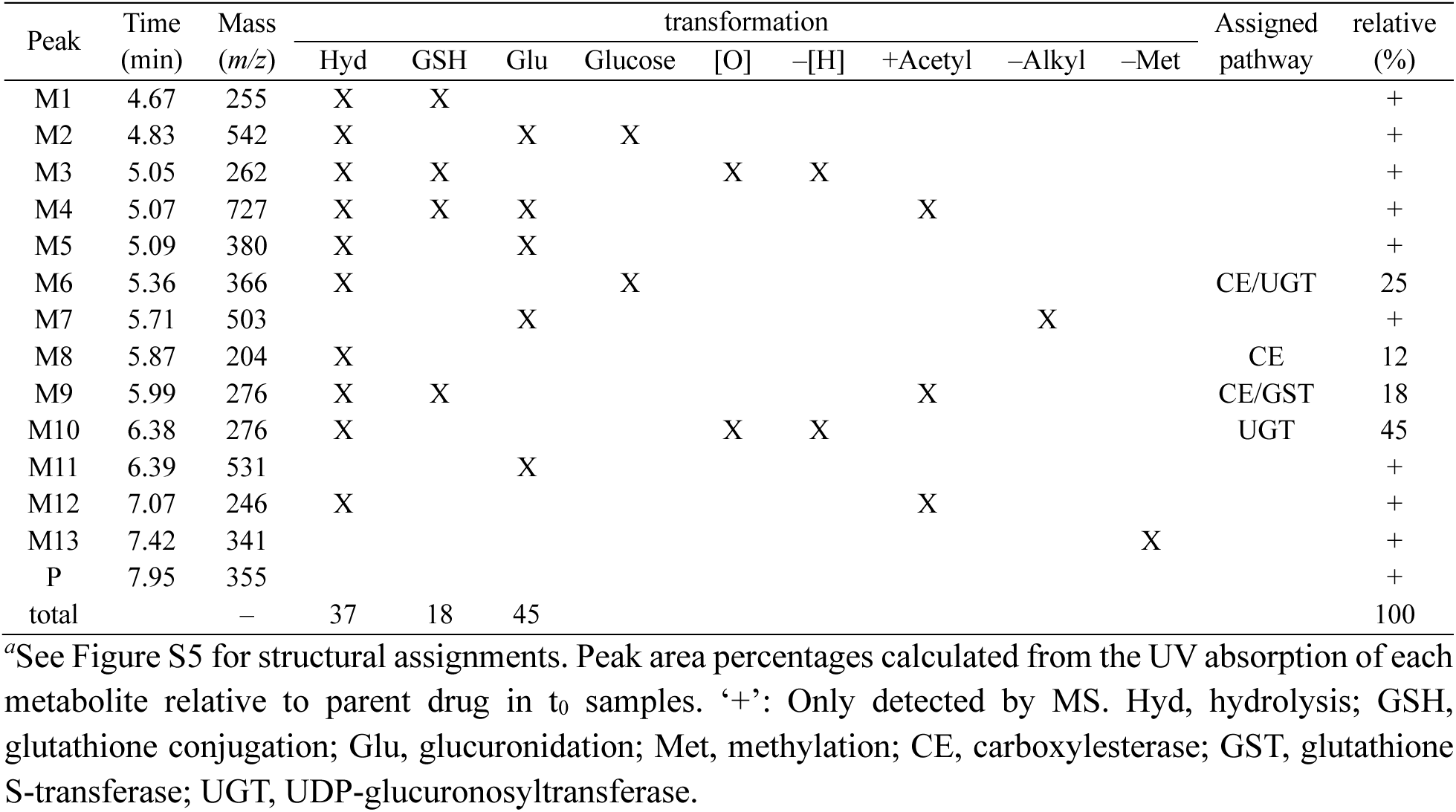
**Metabolites of 5 in Mouse Hepatocytes co-dosed with 1-ABT*^a^***

Four major metabolites of **5** were identified (Figure 5), corresponding to glucuronidation (M10, 45%), amide hydrolysis (M8, 12%), and amide hydrolysis followed by glucose (M6, 25%) or GSH (M9, 18%) conjugation. Additional minor pathways of metabolism were assigned from metabolites that could be detected by MS but were not quantifiable by UV peak area (Table 4 and Figure S5). The major metabolite M10 was most likely formed by the action of UDP-glucuronosyltransferase (UGT) enzymes directly on **5**.^28^ The exact site of glucuronidation in M10 could not be assigned from the MS fragmentation (Figure S5) but is likely to occur on one of the three free NH sites. Amide hydrolysis to form M8 was the other major pathway of metabolism (Figure 5). There are multiple hepatic enzymes that have been reported to hydrolyze other heteroaromatic amides, most commonly carboxyl esterases^29^ but occasionally amine oxidases.^30^

### Modified Mouse Liver Microsome Assay

Together P450 and UGT enzymes account for the majority of hepatic drug metabolism^31^ and the use of liver microsomes supplemented with cofactors has been successful in predicting the metabolism of several known drugs that are substrates for these enzymes.^32^ The metabolic stability of **5** was evaluated in the presence of nicotinamide adenine dinucleotide phosphate (NADPH; cofactor for P450 metabolism) and uridine diphosphate glucuronic acid (UDPGA; cofactor for UGT metabolism) in mouse liver microsomes (MLM, Figure 6). In the presence of NADPH, **5** exhibited a half-life of ∼15 min, demonstrating rapid P450 metabolism most likely on the 2-ethoxy group based on prior analysis of vinyl sulfone **1** and its analogs.^14^ Remarkably, when incubated in MLM supplemented with UDPGA alone, <3% of **5** remained within 15 min, indicating that UGT-mediated glucuronidation contributes substantially to the biotransformation of **5** and was much faster than its P450 metabolism in MLM. Interestingly, when both NADPH and UDPGA were present, the metabolic profile mirrored that observed with UDPGA alone, further suggesting that glucuronidation was the dominant metabolic pathway for **5** in MLM (Figure 6). In the absence of cofactors, **5** showed only a slow decline over 60 min, confirming that metabolic enzymes primarily drive its clearance in MLM. These observations indicated that UGT-catalyzed glucuronidation was a major cause of metabolic liability in *N*-methyl sulfonamide **5**. Furthermore, the use of UDPGA-supplemented MLM provided a simple assay to further study the site of glucuronidation of **5** and potentially develop analogs with improved metabolic stability.

**Figure 6.**
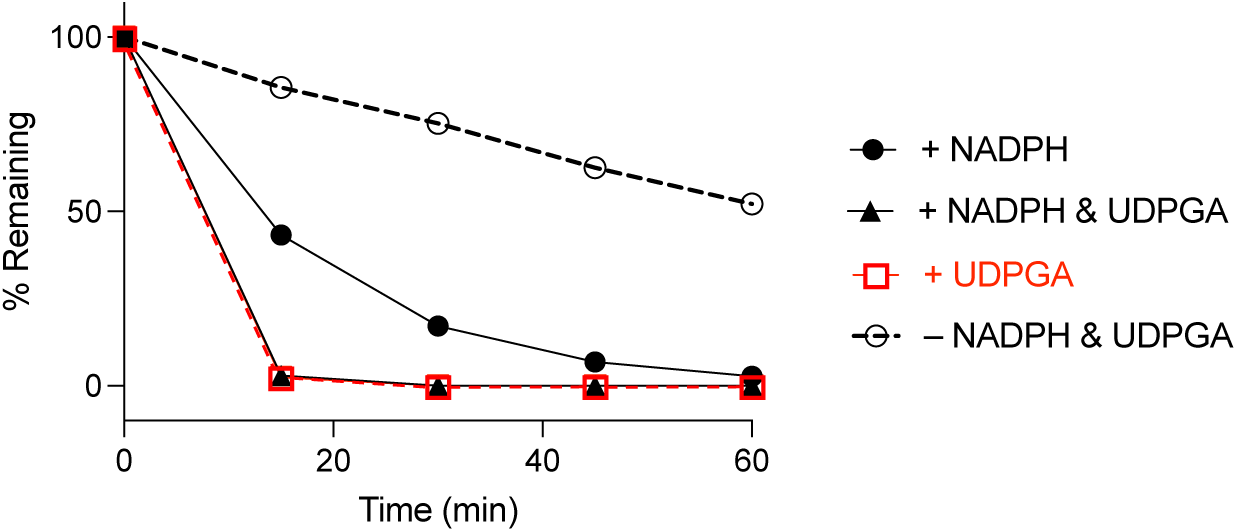
MLM stability assay of **5** (1 mM) in presence and absence of NADPH (10mM) and UDPGA (20 mM) cofactors. Reactions were carried out in triplicate and quenched at designated time points (0, 15, 30, 45, and 60 min). Mass spectrometric analysis with multiple reaction monitoring in negative ion mode was used to detect the remaining quantity of **5**.

### Structure–activity Studies

A systematic study of the sulfamate chemotype was performed to define the structure–activity for antiviral activity and metabolic stability to UGT-catalyzed glucuronidation. The sulfamate acetamide **5** was modified in three regions: a) the reverse acetamide/sulfamate warhead, b) the 5-membered heterocycle, and c) the aryl substituent (Figure 7).

**Figure 7.**
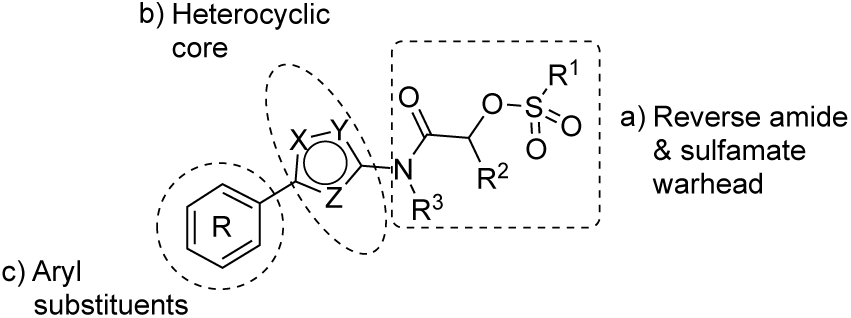
Three regions of structure-activity study

London reported^22^ that sulfamate warheads react with cysteine with release sulfamic acid, which then dissociates into sulfur trioxide and a free amine.^33^ The initial report^22^ demonstrated that the reactivity of the warhead could be tuned by changing the amine substituent to optimize the balance between target selectivity and potency. To test the reactivity of a range of sulfamate warheads for nsP2pro, analogs **6a**–**d** were synthesized (Table 5).

**Table 5.**
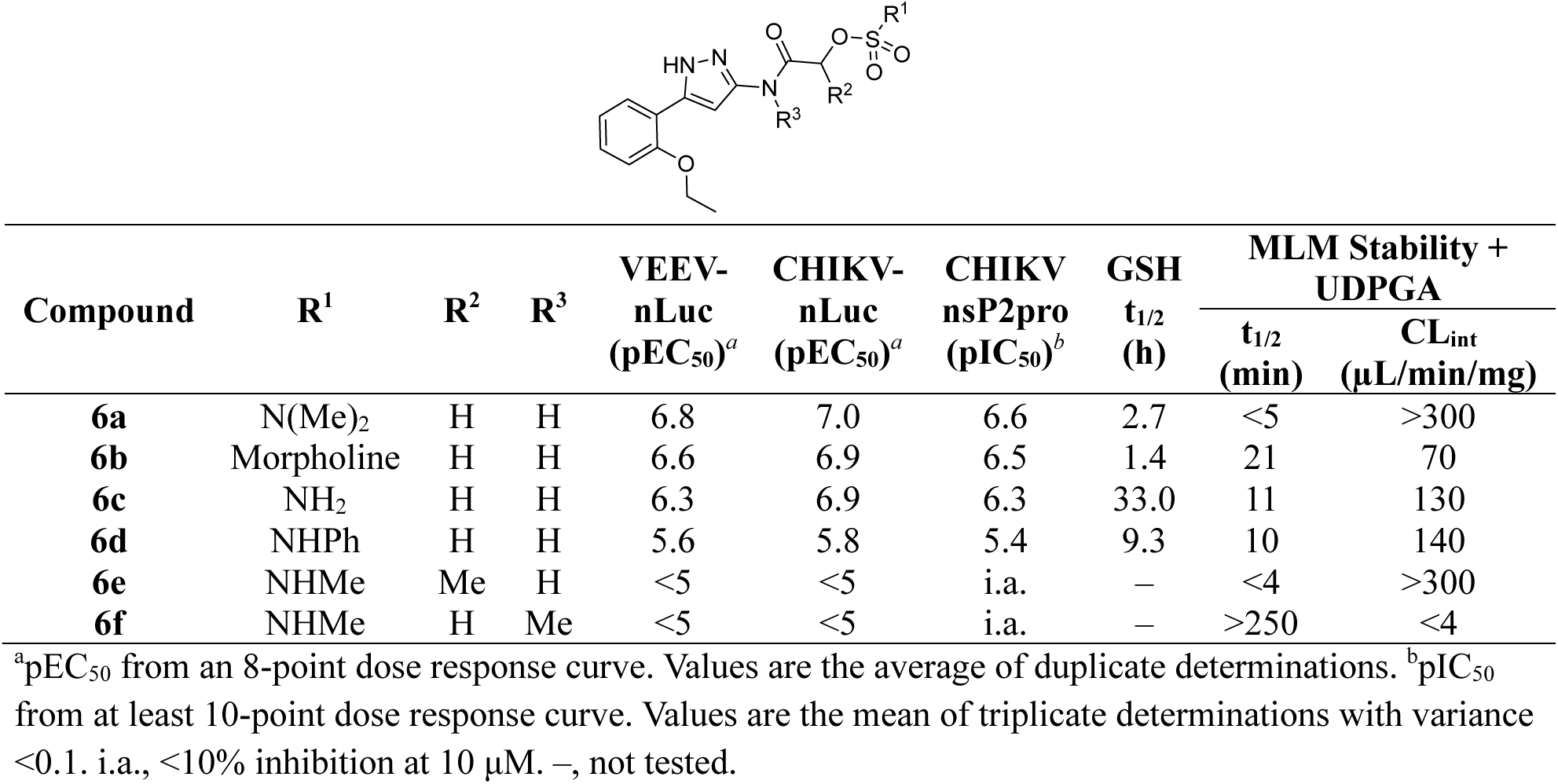
Antiviral Activity, nsP2pro Inhibition, GSH Reactivity, Metabolic Stability of 6a–f.

| Compound | R<sup>1</sup> | R<sup>2</sup> | R<sup>3</sup> | VEEV-nLuc<br>(pEC<sub>50</sub>)<sup>a</sup> | CHIKV-nLuc<br>(pEC<sub>50</sub>)<sup>a</sup> | CHIKV<br>nsP2pro<br>(pIC<sub>50</sub>)<sup>b</sup> | GSH<br>t<sub>1/2</sub><br>(h) | MLM Stability +<br>UDPGA |  |
| --- | --- | --- | --- | --- | --- | --- | --- | --- | --- |
|  |  |  |  |  |  |  |  | t<sub>1/2</sub><br>(min) | CL<sub>int</sub><br>(μL/min/mg) |
| **6a** | N(Me)<sub>2</sub> | H | H | 6.8 | 7.0 | 6.6 | 2.7 | <5 | >300 |
| **6b** | Morpholine | H | H | 6.6 | 6.9 | 6.5 | 1.4 | 21 | 70 |
| **6c** | NH<sub>2</sub> | H | H | 6.3 | 6.9 | 6.3 | 33.0 | 11 | 130 |
| **6d** | NHPh | H | H | 5.6 | 5.8 | 5.4 | 9.3 | 10 | 140 |
| **6e** | NHMe | Me | H | <5 | <5 | i.a. | – | <4 | >300 |
| **6f** | NHMe | H | Me | <5 | <5 | i.a. | – | >250 | <4 |

<sup>a</sup>pEC<sub>50</sub> from an 8-point dose response curve. Values are the average of duplicate determinations. <sup>b</sup>pIC<sub>50</sub> from at least 10-point dose response curve. Values are the mean of triplicate determinations with variance <0.1. i.a., <10% inhibition at 10 μM. –, not tested.

The antiviral activity, nsP2pro inhibition, GSH reactivity, and metabolic stability were assessed (Table 5). *N*-methyl sulfamate **5** had demonstrated potent inhibition of alphaviral replication using the VEEV-nLuc and CHIKV-nLuc reporter assays with EC50 ∼100 nM (Table 1). The *N*,*N*-demethyl (**6a**) and morpholino (**6b**) sulfamates retained activity on both alphaviruses, but the amino analog (**6c**) was 5-fold less potent on VEEV. The anilino sulfamate (**6d**) was >10-fold less active on both alphaviruses. CHIKV nsP2pro inhibition matched the CHIKV antiviral activity. In the GSH reactivity assay, *N*-methyl sulfamate **5** showed a half-life of 14 h. Only the amino analog (**6c**) showed greater stability to GSH, with the *N*,*N*- demethyl (**6a**) and morpholino (**6b**) sulfamates displaying much greater GSH reactivity like the vinyl sulfone **1**. The anilino sulfamate (**6d**) showed similar GSH stability to **5**. Interestingly, the relative GSH stability of the NHMe (**5**) > NHPh (**6d**) > NMe2 (**6a**) sulfamate acetamides was exactly opposite of the trend reported by London et al,^22^ demonstrating that while the chemical reactivity of the sulfamate warhead was tunable the outcome was not predictable. In the UGT-catalyzed glucuronidation assay, using MLM supplemented with UDPGA, the morpholino sulfamate (**6b**) proved to be the most stable with a half-life of 21 min. The *N*,*N*-demethyl sulfamate (**6a**) was cleared as fast as **5**. The amino (**6c**) and anilino (**6d**) sulfamates showed intermediate stability in MLM. From this initial series of analogs, the morpholino sulfamate (**6b**) stood out with the best balance of antiviral potency and metabolic stability despite its greater GSH reactivity. Importantly, the data showed that modification of the sulfamate amino group could modulate the rate of UGT-catalyzed glucuronidation as well as general chemical reactivity.

Two analogs (**6e**–**f**) were synthesized to explore the effect of methyl substitution on the reverse acetamide. Methyl substitution on the methylene of the warhead resulted in an analog (**6e**) that lacked bioactivity but was still rapidly metabolized in MLM. Methyl substitution on the amide NH in **6f** also led to a loss of bioactivity but, notably, the compound was now resistant to glucuronidation in MLM. This observation suggested that the secondary amide NH in **5** might be a primary site of UGT-catalyzed glucuronidation.

Four analogs (**7a**–**d**) were synthesized to explore the effect of varying the heterocyclic core (Table 6). In the structure–activity studies of vinyl sulfone **1**, we found that a wide range of 5-membered heterocycles could substitute for its pyrazole core.^11^ However, we were surprised to find that the corresponding substitutions in the sulfamate series led to analogs devoid of nsP2pro inhibition or antiviral activity (Table 6). Even the conservative pyrazole (**5**) to isoxazole (**7a**) substitution resulted in >100-fold loss in bioactivity. This unanticipated result suggested that the sulfamate acetamide series of covalent inhibitors binds to the nsP2pro enzyme in a different mode or conformation than the vinyl sulfone series. Notably, the analogs (**7a**–**d**) still provided some insight into the structure–activity for metabolic stability. The three of the alternative 5-membered heterocycles: isoxazole **7a**, *N*-methyl pyrazole **7b**, and thiazole **7c** had improved MLM stability, suggesting that the free NH in the pyrazole core of **5** may be a secondary site of glucuronidation.

**Table 6.**
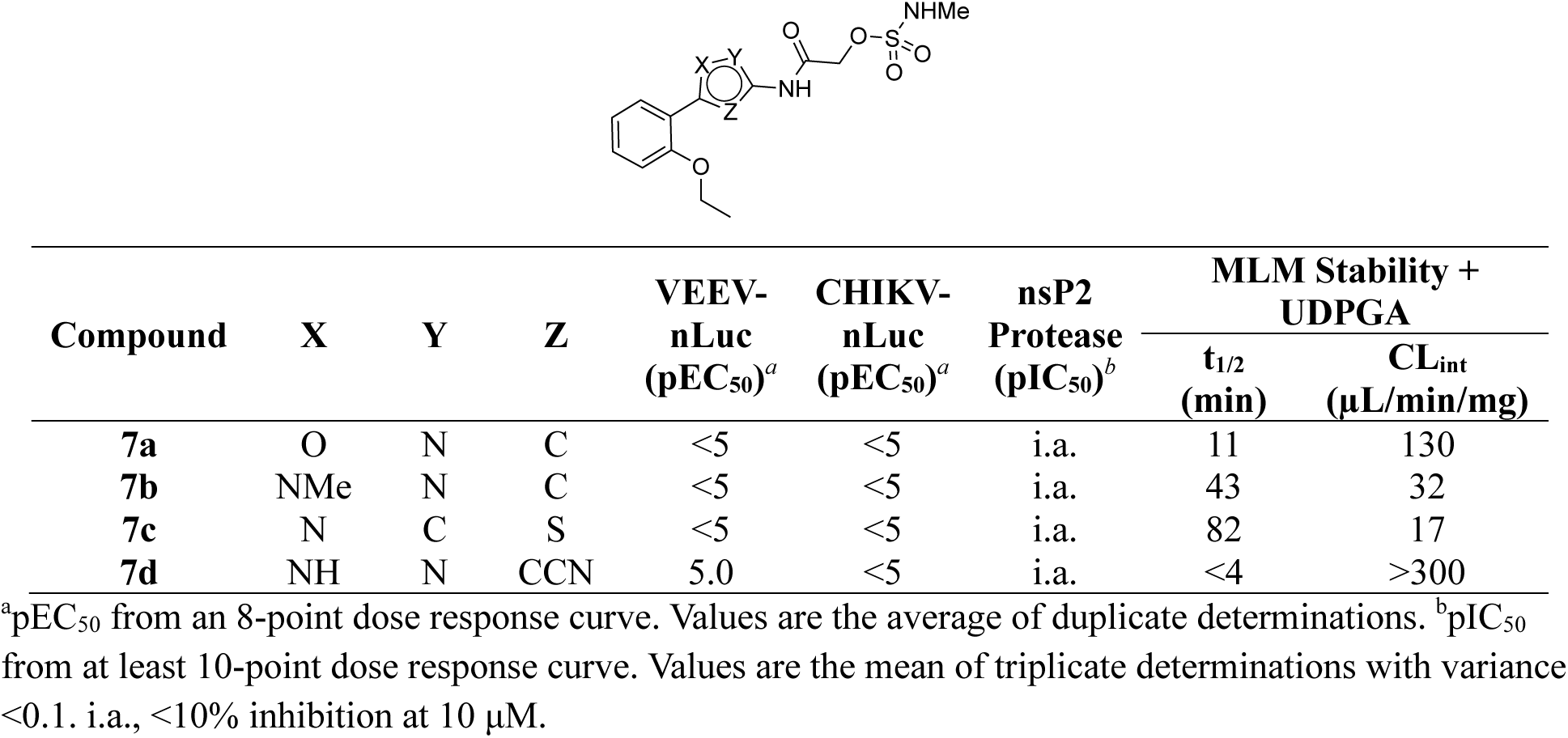
Antiviral Activity, nsP2pro Inhibition, and Metabolic Stability of 7a–d.

| Compound | X | Y | Z | VEEV-nLuc<br>(pEC<sub>50</sub>)<sup>a</sup> | CHIKV-nLuc<br>(pEC<sub>50</sub>)<sup>a</sup> | nsP2<br>Protease<br>(pIC<sub>50</sub>)<sup>b</sup> | MLM Stability +<br>UDPGA |  |
| --- | --- | --- | --- | --- | --- | --- | --- | --- |
|  |  |  |  |  |  |  | t<sub>1/2</sub><br>(min) | CL<sub>int</sub><br>(μL/min/mg) |
| **7a** | O | N | C | <5 | <5 | i.a. | 11 | 130 |
| **7b** | NMe | N | C | <5 | <5 | i.a. | 43 | 32 |
| **7c** | N | C | S | <5 | <5 | i.a. | 82 | 17 |
| **7d** | NH | N | CCN | 5.0 | <5 | i.a. | <4 | >300 |

<sup>a</sup>pEC<sub>50</sub> from an 8-point dose response curve. Values are the average of duplicate determinations. <sup>b</sup>pIC<sub>50</sub> from at least 10-point dose response curve. Values are the mean of triplicate determinations with variance <0.1. i.a., <10% inhibition at 10 μM.

Three fused 5,6-heterocycles (**8a**–**c**) were synthesized to identify an alternative core that would retain bioactivity and be resistant to glucuronidation (Table 7). Unfortunately, the benzoisothiazole (**8a**) was only weakly activity on CHIKV, whereas the benzoisoxazole (**8b**) was only active on VEEV. The pyridoisoxazole (**8c**) was inactive in the antiviral assays but showed activity in the nsP2pro inhibition assay, suggesting that its lack of antiviral activity may be due to poor cell penetration. All three analogs showed only slightly improved metabolic stability compared to pyrazole **5**.

**Table 7.**
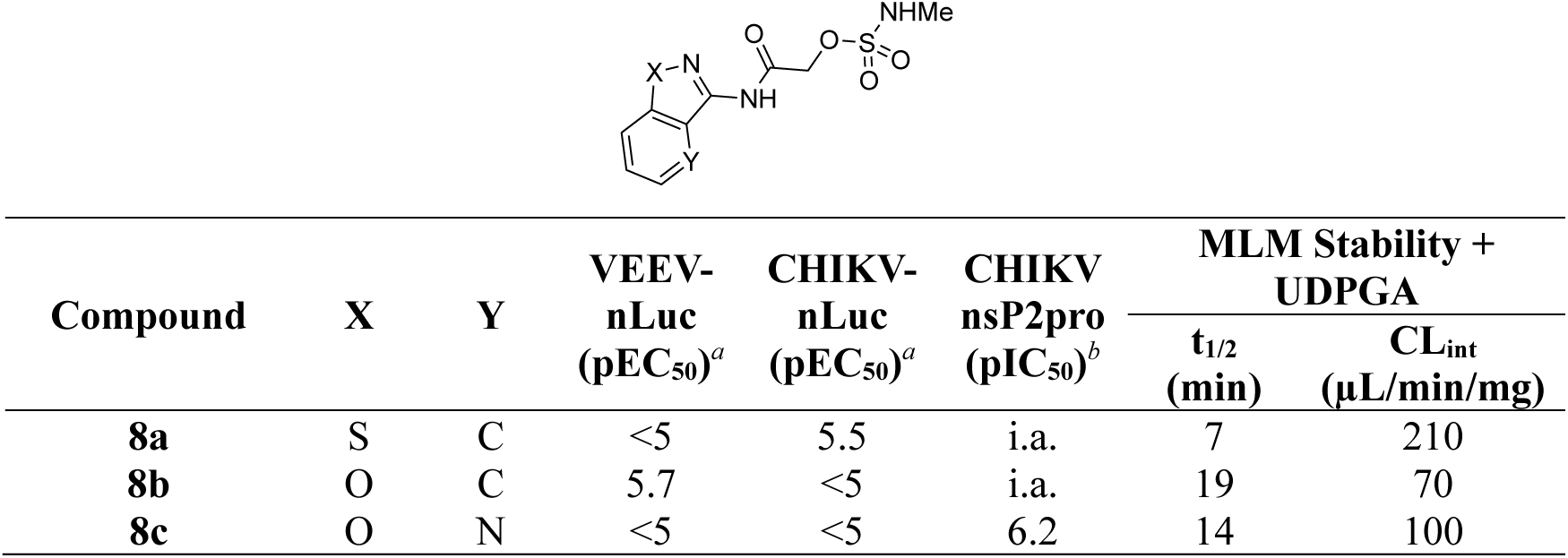
Antiviral Activity, nsP2pro Inhibition, and Metabolic Stability of 8a–c.

Since the 2-ethoxy substituent was a known P450 liability from our prior studies,^14^ the unsubstituted phenyl analog of **5** was synthesized. Unfortunately, **9a** was inactive in the antiviral and nsP2pro assays (Table 8). Unsurprisingly, **9a** was still a good substrate for glucuronidation. Analogs **9b** (**NMS** in Figure 4A) and **9c** with alkyne substituents at the 4-poisition of the phenyl group were synthesized to support the chemoproteomics experiments (Figure 4B). Both analogs showed antiviral activity on VEEV and CHIKV and were also active in the nsP2pro enzyme assay with an IC50 ∼10 μM (Table 8).

**Table 8.** Antiviral Activity, nsP2pro Inhibition, Metabolic Stability of 9a–c.

| Compound | R | VEEV-nLuc<br>(pEC<sub>50</sub>)<sup>a</sup> | CHIKV-nLuc<br>(pEC<sub>50</sub>)<sup>a</sup> | CHIKV<br>nsP2pro<br>(pIC<sub>50</sub>)<sup>b</sup> | MLM Stability +<br>UDPGA |  |
| --- | --- | --- | --- | --- | --- | --- |
| | | | | | t<sub>1/2</sub><br>(min) | CL<sub>int</sub><br>(μL/min/mg) |
| **9a** | H | <5 | <5 | i.a. | 7 | 190 |
| **9b (NMS)** | OCH<sub>2</sub>C≡CH | 5.9 | 5.8 | 5.0 | 31 | 40 |
| **9c** | C≡CH | 5.7 | 5.8 | 5.0 | 43 | 30 |

<sup>a</sup>pEC<sub>50</sub> from an 8-point dose response curve. Values are the average of duplicate determinations. <sup>b</sup>pIC<sub>50</sub> from at least 10-point dose response curve. Values are the mean of triplicate determinations with variance <0.1. i.a. = inactive at 10 μM.

The metabolic ID experiments (Table 4 and Figure 5) identified glucuronidation as a major pathway for hepatic first pass metabolism of **5** but mass spectrometry was unable to identify the exact site of modification (Figure S5). The combined structure–activity in Tables 5–8 showed that the free NH on the secondary acetamide of **5** was likely to be the primary liability for UGT-catalyzed glucuronidation. However, the observation that modification of the free NH groups on either the sulfamate warhead or the pyrazole core also led to small improvements in metabolic stability suggests that these may be secondary sites for glucuronidation.

### Chemistry

The compounds were synthesized following the synthetic routes shown in Schemes 1–3. Acrylamide (**2**), chloroacetamide (**3**), and sulfonate (**4**) electrophiles were introduced by SN2 reaction of 5-(2-ethoxyphenyl)-1*H*-pyrazol-3-amine **10** with acryloyl chloride, 2-chloroacetyl chloride, and 2- ((methylsulfonyl)oxy)acetic acid respectively (Scheme 1). Treatment of **10** with methyl 2-hydroxyacetate afforded *N*-(5-(2-ethoxyphenyl)-1*H*-pyrazol-3-yl)-2-hydroxyacetamide **11**, subsequent sulfamoylation afforded **6c**. Synthesis of sulfamates **5**, **6a**⎼**b**, **6d**⎼**f**, **7a**–**d**, **8a**–**c**, and **9a**–**c** was accomplished using TBTU (for **5**), triazabicyclodecene (for **6a**), triethylaluminum (for **6b**, **6d**, **6e**), and T3P (**6f**, **7a**–**d**, **8a**–**c**, and **9a**– **c)** as coupling reagents (Scheme 1). Amines **12**, **19**, **20**, **21**, **26**, and **27** were synthesized from available building blocks (Scheme 2). Methylation of pyrazole amine **10** afforded a mixture of 5-(2-ethoxyphenyl)- 1-methyl-1*H*-pyrazol-3-amine **12** and 5-(2-ethoxyphenyl)-*N*-methyl-1*H*-pyrazol-3-amine **19** respectively. Azidation of 2-(2-ethoxyphenyl)thiazole-5-carboxylic acid **28** using diphenylphosphoryl azide (DPPA) in *t*-BuOH, followed by TFA-mediated deprotection yielded 2-(2-ethoxyphenyl)thiazol-5-amine **20**. 3-amino- 1*H*-pyrazole-4-carbonitrile **29** was subjected to bromination and subsequent Suzuki coupling with 2- ethoxyphenylboronic acid to furnish 3-amino-5-(2-ethoxyphenyl)-1*H*-pyrazole-4-carbonitrile **21**. 5-(4- (prop-2-yn-1-yloxy)phenyl)-1*H*-pyrazol-3-amine **26** was synthesized over a 7-step reaction sequence from methyl 4-hydroxybenzoate **30**. PMB-protection of hydroxy group, followed by treatment with ACN/*n-*BuLi afforded the ß-ketonitrile intermediate, which underwent a cyclocondensation reaction with hydrazine hydrate to afford pyrazole **31**. Subsequent Boc protection, followed by catalytic hydrogenation furnished intermediate **32**. Alkylation with propargyl bromide, and subsequent Boc-deprotection furnished terminal alkyne **26**. 5-(4-ethynylphenyl)-1*H*-pyrazol-3-amine **27** was synthesized from methyl 4-iodobenzoate **33** over a 4-step reaction sequence. Sonogashira coupling, ß-ketonitrile formation, and subsequent TMS deprotection afforded intermediate **34**. Cyclocondensation with hydrazine hydrate provided **27**. Sulfamate esters **13**–**16** was synthesized by treating methyl 2-hydroxyacetate **35** or methyl 2-hydroxypropanoate **36** with corresponding sulfamoyl chlorides **37**–**40** under basic conditions (Scheme 3). Saponification of ester **16** afforded free acid **17**.

## Conclusions

*N*-Methyl sulfamate **5** represents is a promising new chemotype of covalent nsP2 cysteine protease inhibitors. **5** shows an improved pan-antialphaviral inhibition profile compared to the vinyl sulfone **1**. In addition, **5** does not suffer from the rapid GST-catalyzed GSH conjugation that has complicated the *in vivo* testing of **1** and its analogs.^14^ Surprisingly, **5** has its own metabolic liability resulting from the reverse acetamide, which was identified as a primary site of glucuronidation and was also subject to amide hydrolysis in primary hepatocytes. Our structure–activity studies suggest that glucuronidation can be tempered by modification of the sulfamate warhead or the heterocyclic core. However, a bioisosteric replacement of the amide will ultimately be required to fully address the metabolic liabilities.

Pyrazole **5** is a new example of the use of a sulfamate acetamide as a tunable covalent warhead. Yet, it is clear that additional work is required to better understand the rules for modulating the chemical reactivity of these electrophiles, since our results studying the GSH stability of several sulfamate acetamides are the exact opposite of those reported previously in another series.^22^ Importantly, however, by expanding the range of amines utilized in these warheads we have identified the morpholino sulfamate (e.g. **6b**) as a particularly interesting analog that has potent bioactivity and is more resistant to UGT-catalyzed glucuronidation.

## Experimental Section

### General Information

All reactions were conducted in oven-dried glassware under a dry nitrogen atmosphere unless otherwise specified. All reagents and solvents were obtained from commercial sources and used without further purification. No unexpected safety hazards were encountered during the synthesis. Analytical thin layer chromatography (TLC) was performed on pre-coated silica gel plates (200 μm, F254 indicator), visualized under UV light or by staining with iodine and KMnO₄. Column chromatography utilized pre-loaded silica gel cartridges on a Biotage automated purification system. ¹H and ¹³C NMR spectra were recorded in DMSO-*d*₆ at 400 MHz and 101/126/214 MHz respectively, on a Bruker spectrometer. Chemical shifts (*δ*) are reported in parts per million (ppm) downfield from tetramethylsilane for ¹H NMR, with major peaks designated as s (singlet), d (doublet), t (triplet), q (quartet), and m (multiplet). High-resolution mass spectrometry (HRMS) analyses were performed at the UNC Department of Chemistry Mass Spectrometry Core Laboratory using a Q Exactive HF-X mass spectrometer. Liquid chromatography-mass spectrometry (LC-MS) was conducted on an Agilent 1290 Infinity II LC System with an Agilent Infinity Lab PoroShell 120 EC-C18 column (30 °C, 2.7 μm, 2.1 × 50 mm), employing a 5−95% CH₃CN in water eluent, with 0.2% (v/v) formic acid as the modifier and a flow rate of 1 mL/min. Preparative high-performance liquid chromatography (HPLC) was executed using an Agilent 1260 Infinity II LC System equipped with a Phenomenex C18 column (PhenylHexyl, 30 °C, 5 μm, 75 × 30 mm), with a 5−95% CH₃CN in water eluent and 0.05% (v/v) trifluoroacetic acid as the modifier, at a flow rate of 30 mL/min. Analytical HPLC data were recorded on a Waters Alliance HPLC with a PDA detector or an Agilent 1260 Infinity II series with a PDA detector (EC-C18, 100 mm × 4.6 mm, 3.5 μm), using a 10−90% CH₃CN in water eluent at a flow rate of 1 mL/min. The final compounds were confirmed to be >95% pure by HPLC analysis.

### Chemistry

(*E*)-5-(2-ethoxyphenyl)-*N*-(3-(methylsulfonyl)allyl)-1*H*-pyrazole-3-carboxamide (**1**), 5-(2-ethoxyphenyl)-1*H*-pyrazol-3-amine (**10**), **CA**, **TVS**, and **VSC** were synthesized following the reported procedures.^16, 35^ 5-(2-ethoxyphenyl)isoxazol-3-amine (**18**), benzo[*d*]isothiazol-3-amine (**22**), benzo[*d*]isoxazol-3-amine (**23**), isoxazolo[4,5-*b*]pyridin-3-amine (**24**), 5-phenyl-1*H*-pyrazol-3-amine (**25**) were obtained from commercial sources.

### General Methods

*General Procedure-I.* To a stirring solution of sulfamoyl ester (1.0 eq.) and amine (1.0 eq.) in toluene at 0 °C was added triethylaluminum solution (1.0 M in hexanes) (1.5 eq.). The reaction mixture was stirred at 25 °C for 2 h. On completion, the reaction mixture was poured into water and extracted with EtOAc. The organic layers were dried over anhydrous Na2SO4, filtered, concentrated in vacuo, and the resulting crude product was purified by automated column chromatography or preparative HPLC to afford **6b**, **6d**, **6e**.

*General Procedure-II.* To a stirred solution of amine (1.0 eq.) in THF were added 2-((*N*- methylsulfamoyl)oxy)acetic acid (**17**, 1.5 eq.) and propylphosphonic anhydride (T3P, 50 % in EtOAc) (1.0 eq.) and the reaction was stirred at 70 °C for 1–16 h. On completion, the reaction mixture was poured into water and extracted with EtOAc. The organic layers were dried over anhydrous Na2SO4, filtered, concentrated in vacuo, and the resulting crude product was purified by automated column chromatography or preparative HPLC to afford **6f**, **7a**⎼**d**, **8a**⎼**c**, **9a**⎼**c**.

*General Procedure-III.* To a stirred solution of sodium hydride (60% dispersion in mineral oil, 1.2 eq.) in THF/DMF or triethylamine (3.0 eq.) in CH2Cl2, methyl 2-hydroxyacetate (1.0 eq.) and alkyl/aryl sulfonyl chloride (1.0 eq) were added at 0 °C. The reaction mixture was stirred at 25 °C for 2–6 h. On completion, the reaction was poured into 1N citric acid solution and extracted with EtOAc. Combined organic layers were dried over anhydrous Na2SO4, filtered, concentrated, and the resulting crude product was purified by automated column chromatography to afford **13**–**16**.

*N-(3-(2-ethoxyphenyl)-1H-pyrazol-5-yl)acrylamide (**2**).* To a stirred solution of 3-(2- ethoxyphenyl)-1*H*-pyrazol-5-amine (**10**, 0.200 g, 0.985 mmol, 1.0 eq.) and *N,N*-diisopropylethylamine (0.381 g, 2.95 mmol, 3.0 eq.) in CH2Cl2 was added acryloyl chloride (0.04 mL, 0.492 mmol, 0.5 eq.) at 0 °C. The reaction was stirred at 0 °C for 2 h. On completion, the reaction mixture was diluted with water and the organic layer was extracted with CH2Cl2. Combined organic layers were dried over anhydrous Na2SO4, filtered, concentrated, and the resulting crude product was purified by preparative HPLC to afford **2** as a white solid (0.027 g, 11% yield, m.p. 97 °C). HPLC purity >99%. ^1^H NMR (400 MHz, DMSO-*d*6): *δ* 10.63 (s, 1H), 7.66 (d, *J* = 7.7 Hz, 1H), 7.32 (t, *J* = 7.8 Hz, 1H), 7.12 (d, *J* = 8.3 Hz, 1H), 7.07 (s, 1H), 7.01 (t, *J* = 7.5 Hz, 1H), 6.50 (dd, *J* = 16.8, 10.3 Hz, 1H), 6.25 (dd, *J* = 17.0, 2.2 Hz, 1H), 5.75 – 5.66 (m, 1H), 4.16 (q, *J* = 7.0 Hz, 3H), 1.40 (t, *J* = 6.9 Hz, 3H). ^13^C NMR (101 MHz, DMSO-*d*6): *δ* 162.2, 154.9, 147.2, 138.9, 131.5, 129.4, 127.3, 126.5, 120.6, 118.3, 112.9, 96.8, 63.7, 14.6. HRMS (ESI) *m/z*: [M+H]^+^ calculated for C13H15ClN3O2: 280.0853, found 280.0846.

*2-chloro-N-(3-(2-ethoxyphenyl)-1H-pyrazol-5-yl)acetamide (**3**).* To a stirred solution of 3-(2- ethoxyphenyl)-1*H*-pyrazol-5-amine (**10** , 0.200 g, 0.985 mmol, 1.0 eq.) in CH2Cl2 was added triethylamine (0.1 mL, 1.477 mmol, 1.5 eq.) and 2-chloroacetyl chloride (0.131 g, 1.182 mmol, 1.2 eq.) dropwise at 0 °C. The reaction was stirred at 25 °C for 2 h. On completion, the reaction mixture was diluted with water and extracted with EtOAc. Combined organic layers were dried over anhydrous Na2SO4, filtered, concentrated, and the crude product was purified by preparative HPLC to afford **3** as a white solid (0.063 g, 23% yield, m.p. 193 °C). HPLC purity >99%. ^1^H NMR (400 MHz, DMSO-*d*6): *δ* 12.52 (s, 1H), 10.75 (s, 1H), 7.65 (dd, *J* = 7.7, 1.7 Hz, 1H), 7.32 (ddd, *J* = 8.7, 7.3, 1.7 Hz, 1H), 7.12 (d, *J* = 8.3 Hz, 1H), 7.04 – 6.98 (m, 2H), 4.25 (s, 2H), 4.15 (q, *J* = 6.9 Hz, 2H), 1.40 (t, *J* = 6.9 Hz, 3H). ^13^C NMR (101 MHz, DMSO-*d*6): *δ* 163.7, 154.9, 146.9, 138.7, 129.4, 127.3, 120.6, 117.9, 112.9, 96.6, 63.7, 43.0, 14.5. HRMS (ESI) *m/z*: [M+H]^+^calculated for C13H15ClN3O2: 280.0853, found 280.0846.

*2-((5-(2-ethoxyphenyl)-1H-pyrazol-3-yl)amino)-2-oxoethyl methanesulfonate (**4**).* To a stirring solution of 5-(2-ethoxyphenyl)-1*H*-pyrazol-3-amine (**10**, 0.158 g, 0.78 mmol, 1.2 eq.) and 2-((methylsulfonyl)oxy)acetic acid (0.100 g, 0.65 mmol, 1.0 eq.) in DMF/EtOAc (1:1, 3 mL) were added diisopropylethylamine (0.2 mL, 1.3 mmol, 2.0 eq.) and propylphosphonic anhydride (50% by wt., 0.764 mL, 2.0 eq, 1.30 mmol) and the reaction was stirred at 25 °C for 2 h. On completion, the reaction mixture was quenched with water and extracted with EtOAc. Combined organic layers were dried over anhydrous Na2SO4, filtered, concentrated, and the crude product was purified using automated column chromatography followed by reverse phase HPLC to afford **4** as a white solid (0.103 g, 35% yield, m.p. 120 °C). HPLC purity >99%. ^1^H NMR (500 MHz, DMSO-*d*6): *δ* 7.90 (dd, *J* = 7.7, 1.7 Hz, 1H), 7.36 (ddd,

*J* = 8.3, 7.3, 1.8 Hz, 1H), 7.10 (dd, *J* = 8.4, 1.0 Hz, 1H), 6.99 (td, *J* = 7.4, 1.1 Hz, 1H), 6.63 (s, 1H), 5.98 (s, 1H), 5.61 (s, 2H), 4.11 (q, *J* = 6.9 Hz, 2H), 3.34 (s, 3H), 1.40 (t, *J* = 6.9 Hz, 3H). ^13^C NMR (126 MHz, DMSO-*d*6): *δ* 167.2, 156.7, 152.6, 151.1, 130.5, 128.4, 120.4, 120.3, 112.9, 88.9, 67.1, 63.7, 37.7, 14.7. HRMS (ESI) *m/z*: [M+H]^+^ calculated for C14H18N3O5S: 340.0967, found 340.0955.

*2-((5-(2-ethoxyphenyl)-1H-pyrazol-3-yl)amino)-2-oxoethyl methylsulfamate (**5**).* To a stirred solution of 2-((*N*-methylsulfamoyl)oxy)acetic acid (**17**, 0.244 g, 1.477 mmol, 1.0 eq.) in pyridine (3.0 mL) was added 5-(2-ethoxyphenyl)-1*H*-pyrazol-3-amine (**10**, 0.300 g, 1.477 mmol, 1.0 eq) and TBTU (0.711 g, 2.216 mmol, 1.5 eq) at 0 °C. The reaction was stirred at 25 °C for 2 h. On completion, the reaction mixture was quenched with water and extracted with EtOAc. The combined organic layers were dried over anhydrous Na2SO4, filtered, concentrated, and the crude product was purified by automated column chromatography (eluent 50% EtOAc /hexane) followed by reverse phase HPLC to afford **5** (0.130 g, 25% yield, m.p. 172–174 °C). HPLC purity >99%. ^1^H NMR (401 MHz, DMSO-*d*6): *δ* 12.50 (s, 1H), 10.60 (s, 1H), 7.88 (d, *J* = 4.4 Hz, 1H), 7.65 (dd, *J* = 7.7, 1.7 Hz, 1H), 7.36 – 7.28 (m, 1H), 7.13 (s, 1H), 7.05 – 6.96 (m, 2H), 4.64 (s, 2H), 4.16 (q, *J* = 6.9 Hz, 2H), 2.64 (d, *J* = 4.7 Hz, 3H), 1.40 (t, *J* = 6.9 Hz, 3H). ^13^C NMR (101 MHz, DMSO-*d*6): *δ* 163.1, 154.9, 146.7, 138.7, 129.4, 127.3, 120.6, 117.9, 112.9, 96.8, 66.4, 63.7, 28.8, 14.5. HRMS (ESI) *m/z*: [M+H]^+^ calculated for C14H19N4O5S: 355.1076, found 355.1064.

*2-((5-(2-ethoxyphenyl)-1H-pyrazol-3-yl)amino)-2-oxoethyl dimethylsulfamate (**6a**).* Step-1: preparation according to General Procedure-III using methyl 2-hydroxyacetate (2.0 g, 22.20 mmol, 1.0 eq.), sodium hydride (60% dispersion in mineral oil) (1.3 g, 33.30 mmol, 1.5 eq.), and dimethylsulfamoyl chloride (3.18 g, 22.20 mmol, 1.0 eq.) in DMF at 25 °C for 6 h. The crude product was purified using automated column chromatography (eluent 25% EtOAc/hexane) to afford methyl 2-((*N*,*N*- dimethylsulfamoyl)oxy)acetate (**13**) as a white solid (0.300 g, 7% yield). LCMS: *m/z* = 198.1 [M+1]^+^. Step- 2: to a stirring solution of methyl 2-((*N*,*N*-dimethylsulfamoyl)oxy)acetate **13** (0.300 g, 1.521 mmol, 1.0 eq.) in THF (10 mL) were added 5-(2-ethoxyphenyl)-1*H*-pyrazol-3-amine **10** (0.309 g, 1.521 mmol, 1.0 eq) and triazabicyclodecene (0.190 g, 1.369 mmol, 0.9 eq.). The reaction mixture was stirred at 25 °C for 2 h. On completion, the reaction mixture was diluted with water and extracted with EtOAc. Combined organic layers were dried over anhydrous Na2SO4, filtered, concentrated, and the crude product was purified by column chromatography (eluent 10% acetone/ CH2Cl2) to afford **6a** as a white solid (0.010 g, 2% yield, m.p 137–139 °C). HPLC purity >99%. ^1^H NMR (400 MHz, DMSO-*d*6): *δ* 12.53 (s, 1H), 10.65 (s, 1H), 7.65 (d, *J* = 7.7 Hz, 1H), 7.32 (t, *J* = 7.6 Hz, 1H), 7.12 (d, *J* = 8.4 Hz, 1H), 7.02 (d, *J* = 9.1 Hz, 2H), 4.75 (s, 2H), 4.16 (q, *J* = 6.9 Hz, 2H), 2.87 (s, 6H), 1.40 (t, *J* = 6.9 Hz, 3H). ^13^C NMR (101 MHz, DMSO-*d*6): *δ* 163.1, 154.9, 146.7, 138.7, 129.4, 127.4, 120.6, 117.5, 112.9, 96.7, 67.1, 63.7, 37.9, 14.5. HRMS (ESI) *m/z*: [M+H]^+^ calculated for C15H21N4O5S: 369.1233, found 369.1223.

*2-((5-(2-ethoxyphenyl)-1H-pyrazol-3-yl)amino)-2-oxoethyl morpholine-4-sulfonate (**6b**).* Step-1: preparation according to General Procedure-III using methyl 2-hydroxyacetate (1.0 g, 11.10 mmol, 1.0 eq.), sodium hydride (60% dispersion in mineral oil) (0.532 g, 13.32 mmol, 1.2 eq.), and morpholine-4-sulfonyl chloride (2.06 g, 11.10 mmol, 1.0 eq.) in THF at 25 °C for 2 h. The crude product was purified using automated column chromatography (eluent 30% EtOAc/hexane) to afford methyl 2- ((morpholinosulfonyl)oxy)acetate (**14**) as a white solid (1.2 g, 45% yield). LCMS: *m/z* = 240.1 [M+H]^+^. Step-2: Preparation according to General Procedure-I using 5-(2-ethoxyphenyl)-1*H*-pyrazol-3-amine (**10** , 0.170 g, 0.836 mmol, 1.0 eq.) and methyl 2-((morpholinosulfonyl)oxy)acetate (**14**, 0.200 g, 0.836 mmol, 1.0 eq.) in toluene (2.0 mL) at 25 °C for 2 h. The product was purified using automated column chromatography (eluent 80% EtOAc/hexane), followed by preparative HPLC to afford **6b** as a white solid (0.040 mg, 12% yield, m.p 140–142 °C). HPLC purity >97%. ^1^H NMR (400 MHz, DMSO-*d*6): *δ* 12.59 (s, 1H), 10.71 (s, 1H), 7.69 – 7.63 (m, 1H), 7.37 – 7.27 (m, 1H), 7.12 (d, *J* = 8.3 Hz, 1H), 7.05 – 6.95 (m, 2H), 4.80 (s, 2H), 4.15 (q, *J* = 6.9 Hz, 2H), 3.65 (dd, *J* = 5.9, 3.5 Hz, 4H), 3.25 (t, *J* = 4.5 Hz, 4H), 1.43 – 1.36 (m, 3H). ^13^C NMR (214 MHz, DMSO-*d*6): *δ* 162.9, 154.9, 129.4, 127.4, 120.6, 118.1, 112.9, 96.7, 67.5, 65.3, 63.7, 63.3, 47.2, 46.2(9), 46.2(7), 42.9, 14.5. HRMS (ESI) *m/z*: [M+H]^+^ calculated for C17H23N4O6S: 411.1338, found 411.1332.

*2-((5-(2-ethoxyphenyl)-1H-pyrazol-3-yl)amino)-2-oxoethyl sulfamate (**6c**).* Step-1: to a stirred solution of 5-(2-ethoxyphenyl)-1*H*-pyrazol-3-amine (0.200 g, 0.985 mmol, 1 eq.) in toluene (5.0 mL) were added methyl 2-hydroxyacetate (0.089 mg, 0.985 mmol, 1 eq.) and triethylaluminium (1.0 M in hexane) (2.46 mL, 2.46 mmol, 2.5 eq.) at 0 °C. The reaction was stirred at 80 °C for 16 h. On completion, the reaction mixture was quenched with NH4Cl solution and extracted with EtOAc. The organic layers were dried over anhydrous Na2SO4, filtered, concentrated in vacuo, and the resulting crude was purified using automated column chromatography (eluent 85% EtOAc/hexane) to afford *N*-(5-(2-ethoxyphenyl)-1*H*- pyrazol-3-yl)-2-hydroxyacetamide (0.130 g, 51% yield). LCMS: *m/z =* 262.1 [M–H]^+^. Step-2: to a stirred solution of *N*-(5-(2-ethoxyphenyl)-1*H*-pyrazol-3-yl)-2-hydroxyacetamide (0.120 g, 0.459 mmol, 1.0 eq.) in DMA (4.0 mL) was added sulfamoyl chloride (0.106 g, 0.919 mmol, 2.0 eq.) and the reaction was stirred at 25 °C for 2 h. On completion, the reaction mixture was poured into water and extracted with EtOAc. The organic layers were dried over anhydrous Na2SO4, filtered, concentrated in vacuo, and the crude product was purified using automated column chromatography (eluent 40% EtOAc/hexane) followed by preparative HPLC to afford **6c** as a white solid (0.028 g, 18% yield, m.p. 112–114 °C). HPLC purity >99%. ^1^H NMR (401 MHz, DMSO-*d*6): *δ* 12.51 (s, 1H), 10.49 (s, 1H), 7.65 (d, *J* = 8.2 Hz, 3H), 7.36 – 7.27 (m, 1H), 7.12 (d, *J* = 8.3 Hz, 1H), 7.05 – 6.96 (m, 2H), 4.64 (s, 2H), 4.15 (q, *J* = 6.9 Hz, 2H), 1.40 (t, *J* = 6.9 Hz, 3H). ^13^C NMR (101 MHz, DMSO-*d*6): *δ* 163.3, 154.9, 146.7, 138.7, 129.4, 127.3, 120.6, 117.9, 112.9, 96.8, 66.2, 63.7, 14.5. HRMS (ESI) *m/z*: [M+Na]^+^ calculated for C13H16N4NaO5S: 363.0739, found 363.0733.

*2-((5-(2-ethoxyphenyl)-1H-pyrazol-3-yl)amino)-2-oxoethyl phenylsulfamate (**6d**).* Step-1: preparation according to General Procedure-III using methyl 2-hydroxyacetate (4.0 g, 20.87 mmol, 1.0 eq.), triethylamine (9.03 mL, 62.62 mmol, 3.0 eq.), and phenylsulfamoyl chloride (3.76 g, 41.74 mmol, 2.0 eq.) in CH2Cl2 at 25 °C for 2 h. The crude was purified using automated column chromatography (eluent 30% EtOAc/hexane) to afford methyl 2-((*N*-phenylsulfamoyl)oxy)acetate (**15**) as a white solid (0.15 g, 7% yield). LCMS: *m/z* = 244.0 [M–H]^−^. Step-2: preparation according to General Procedure-I using 5-(2- ethoxyphenyl)-1*H*-pyrazol-3-amine (**10**, 0.43 g, 2.14 mmol, 1.0 eq.) and methyl 2-((*N*-phenylsulfamoyl)oxy)acetate (**15**, 0.350 g, 1.42 mmol, 1.0 eq) in THF (5 mL) at 25 °C for 2 h. The product was purified using automated column chromatography (eluent 10% Acetone: CH2Cl2) to afford **6d** as a white solid (0.015 g, 2.5% yield, m.p. 148–150 °C). HPLC purity >97%. ^1^H NMR (400 MHz, DMSO-*d*6): *δ* 12.51 (s, 1H), 10.69 (d, *J* = 28.7 Hz, 2H), 7.64 (d, *J* = 7.7 Hz, 1H), 7.39 – 7.29 (m, 3H), 7.23 (d, *J* = 7.8 Hz, 2H), 7.12 (d, *J* = 7.9 Hz, 2H), 7.04 – 6.95 (m, 2H), 4.74 (s, 2H), 4.15 (q, *J* = 6.9 Hz, 2H), 1.39 (t, *J* = 6.9 Hz, 3H). ^13^C NMR (101 MHz, DMSO-*d*6): *δ* 162.4, 154.9, 146.6, 138.7, 137.1, 129.4, 129.2, 127.3, 124.1, 120.6, 119.7, 117.9, 112.9, 96.8, 67.1, 63.7, 14.5. HRMS (ESI) *m/z*: [M+H]^+^ calculated for C19H21N4O5S: 417.1233, found 417.1224.

*1-((5-(2-ethoxyphenyl)-1H-pyrazol-3-yl)amino)-1-oxopropan-2-yl methylsulfamate (**6e**).* Step-1: preparation according to General Procedure-III using methyl 2-hydroxypropanoate (0.500 g, 4.807 mmol, 1.0 eq.), triethylamine (0.984 mL, 7.210 mmol, 1.5 eq.), and methylsulfamoyl chloride (0.744 g, 5.769 mmol, 1.2 eq.) in CH2Cl2 at 25 °C for 2 h. The crude was purified using automated column chromatography (eluent 50% EtOAc/hexane) to afford methyl 2-((*N*-methylsulfamoyl)oxy)propanoate (**16**) as a colorless liquid (0.400 g, 42% yield). LCMS: *m/z* = 198.1 [M+H]^+^ Step-2: Preparation according to General Procedure-I using 5-(2-ethoxyphenyl)-1*H*-pyrazol-3-amine (**10**, 0.412 g, 2.03 mmol, 1.0 eq.) and methyl 2-((*N*-methylsulfamoyl)oxy)propanoate (**16**, 0.400 g, 2.03 mmol, 1.0 eq.) in toluene at 25 °C for 2 h. The product was purified by preparative HPLC to afford **6e** as a white solid (0.015 g, 2% yield, m.p 158–160 °C). HPLC purity >96%. ^1^H NMR (400 MHz, DMSO-*d*6): *δ* 12.52 (s, 1H), 10.56 (s, 1H), 7.80 (s, 1H), 7.64 (d, *J* = 7.7 Hz, 1H), 7.32 (t, *J* = 7.8 Hz, 1H), 7.12 (d, *J* = 8.4 Hz, 1H), 7.06 – 6.95 (m, 2H), 4.98 (q, *J* = 6.7 Hz, 1H), 4.15 (q, *J* = 6.5 Hz, 2H), 2.59 (s, 3H), 1.48 (d, *J* = 6.7 Hz, 3H), 1.40 (t, *J* = 6.9 Hz, 3H). ^13^C NMR (101 MHz, DMSO-*d*6): *δ* 167.8, 155.6, 146.9, 139.7, 130.5, 128.1, 121.5, 118.3, 113.6, 97.3, 75.5, 64.5, 29.3, 19.2, 15.1. HRMS (ESI) *m/z*: [M+H]^+^ calculated for C15H21N4O5S: 369.1233, found 369.1225.

*2-((5-(2-ethoxyphenyl)-1H-pyrazol-3-yl)(methyl)amino)-2-oxoethyl methylsulfamate (**6f**).* Step-1: to a stirred solution of 5-(2-ethoxyphenyl)-1*H*-pyrazol-3-amine (**10**, 2.5 g, 12.31 mmol, 1.0 eq.) in DMF were added potassium carbonate (5.09 g, 36.94 mmol, 3.0 eq.), methyl iodide (2.55 g, 18.47 mmol, 1.5 eq.) and the reaction was stirred at 25 °C for 12 h. On completion, the reaction mixture was poured into water and extracted with EtOAc. The combined organic layers were washed with brine, dried over anhydrous Na2SO4, concentrated in vacuo, and the crude product was purified using automated column chromatography (eluent 10% EtOAc/hexane) to afford a mixture of 5-(2-ethoxyphenyl)-*N*-methyl-1*H*- pyrazol-3-amine (**12**) and 5-(2-ethoxyphenyl)-1-methyl-1*H*-pyrazol-3-amine (**19**). Total yield 1.21 g, 45%. LCMS: *m/z* = 218.2 [M+H]^+^. Step-2: to a stirred solution of the mixture of 5-(2-ethoxyphenyl)-*N*-methyl- 1*H*-pyrazol-3-amine (**12**) and 5-(2-ethoxyphenyl)-1-methyl-1*H*-pyrazol-3-amine (**19**) (0.8 g, 3.68 mmol, 1.0 eq.) in CH2Cl2 was added Boc anhydride (8.2 g, 37.69 mmol, 1.2 eq.), trimethylamine (5.4 mL, 37.69 mmol, 3.0 eq.), and the reaction was stirred at 25 °C for 5 h. On completion, the reaction mixture was poured into water and extracted with EtOAc. The combined organic layers were washed with brine, dried over anhydrous Na2SO4, concentrated in vacuo, and the resulting crude was purified using automated column chromatography (eluent 20% EtOAc/hexane) to afford *tert*-butyl 5-(2-ethoxyphenyl)-3- (methylamino)-1*H*-pyrazole-1-carboxylate as a colorless oil (0.45 g, 39% yield). LCMS: *m/z* = 318.3 [M+H]^+^. Step-3: Preparation according to General Procedure-II using 5 *tert*-butyl 5-(2-ethoxyphenyl)-3- (methylamino)-1*H*-pyrazole-1-carboxylate (0.300 g, 0.945 mmol, 1.0 eq.) and 2-((*N*-methylsulfamoyl)oxy)acetic acid (**17**, 0.159 g, 0.945 mmol, 1.0 eq.) in T3P (0.3 mL) at 70 °C for 1 h. The product was purified using automated column chromatography (eluent 60% EtOAc/hexane), followed by preparative HPLC to afford **6f** as a white sticky solid (0.018 g, 5% yield). HPLC purity >96%. ^1^H NMR (401 MHz, DMSO-*d*6): *δ* 12.96 (s, 1H), 7.79 (d, *J* = 5.0 Hz, 1H), 7.72 – 7.66 (m, 1H), 7.36 (t, *J* = 7.5 Hz, 1H), 7.15 (d, *J* = 8.3 Hz, 1H), 7.06 – 7.00 (m, 1H), 6.72 (s, 1H), 4.70 (s, 2H), 4.17 (q, *J* = 6.9 Hz, 2H), 3.23 (s, 3H), 2.57 (d, *J* = 5.0 Hz, 3H), 1.41 (t, *J* = 7.0 Hz, 3H). ^13^C NMR (101 MHz, DMSO-*d*6): *δ* 165.1, 154.9, 149.6, 140.2, 129.9, 127.4, 120.6, 117.3, 112.8, 99.7, 65.8, 63.7, 35.5, 28.7, 14.4. HRMS (ESI) *m/z*: [M+H]^+^ calculated for C15H21N4O5S: 369.1233, found 369.1225.

*2-((5-(2-ethoxyphenyl)isoxazol-3-yl)amino)-2-oxoethyl methylsulfamate (**7a**).* Preparation according to General Procedure-II using 5-(2-ethoxyphenyl)isoxazol-3-amine (**18**, 120 mg, 0.59 mmol, 1.0 eq.) and 2-((*N*-methylsulfamoyl)oxy)acetic acid (**17**, 99.4 mg, 0.59 mmol, 1.0 eq.) in T3P (1.2 mL) at 70 °C for 2 h. The product was purified using automated column chromatography (eluent 40% EtOAc/hexane), followed by preparative HPLC to afford **7a** as a white solid (0.059 g, 28% yield, m.p. 164 °C). HPLC purity >98%. ^1^H NMR (400 MHz, DMSO-*d*6): *δ* 11.32 (s, 1H), 7.95 (q, *J* = 4.8 Hz, 1H), 7.86 (dd, *J* = 7.8, 1.7 Hz, 1H), 7.49 (ddd, *J* = 8.9, 7.5, 1.7 Hz, 1H), 7.37 (s, 1H), 7.22 (dd, *J* = 8.6, 1.0 Hz, 1H), 7.10 (td, *J* = 7.5, 1.0 Hz, 1H), 4.72 (s, 2H), 4.22 (q, *J* = 6.9 Hz, 2H), 2.63 (d, *J* = 4.8 Hz, 3H), 1.45 (t, *J* = 6.9 Hz, 3H). ^13^C NMR (101 MHz, DMSO-*d*6): *δ* 165.1, 164.7, 158.2, 155.3, 131.9, 126.7, 120.8, 115.2, 113.0, 97.8, 66.2, 64.1, 28.8, 14.5. HRMS (ESI) *m/z*: [M+H]^+^ calculated for C14H18N3O6S: 356.0916, found 356.0907.

*2-((5-(2-ethoxyphenyl)-1-methyl-1H-pyrazol-3-yl)amino)-2-oxoethyl methylsulfamate (**7b**).* Preparation according to General Procedure-II using 5-(2-ethoxyphenyl)-1-methyl-1*H*-pyrazol-3-amine (**19**, 0.2 g, 0.92 mmol, 1.0 eq.) and 2-((*N*-methylsulfamoyl)oxy)acetic acid (**17**, 0.155 g, 0.92, 1.0 eq.) in T3P (2.0 mL) at 70 °C for 2 h. The product was purified by preparative HPLC to afford **7b** as a white solid (0.110 g, 33% yield, m.p. 119–121 °C). HPLC purity >99%. ^1^H NMR (401 MHz, DMSO-*d*6): *δ* 10.23 (s, 1H), 7.98 (q, *J* = 4.8 Hz, 1H), 7.87 (dd, *J* = 7.7, 1.8 Hz, 1H), 7.25 (ddd, *J* = 8.9, 7.3, 1.8 Hz, 1H), 7.06 (dd, *J* = 8.4, 1.1 Hz, 1H), 6.96 (td, *J* = 7.5, 1.1 Hz, 1H), 6.75 (s, 1H), 4.72 (s, 2H), 4.10 (q, *J* = 6.9 Hz, 2H), 3.71 (s, 3H), 2.65 (d, *J* = 4.8 Hz, 3H), 1.39 (t, *J* = 6.9 Hz, 3H). ^13^C NMR (101 MHz, DMSO-*d*6): *δ* 164.8, 155.5, 145.2, 135.5, 128.6, 127.1, 121.8, 120.4, 112.8, 100.9, 66.3, 63.5, 35.7, 28.9, 14.8. HRMS (ESI) *m/z*: [M+H]^+^ calculated for C15H21N4O5S: 369.1233, found 369.1219.

*2-((2-(2-ethoxyphenyl)thiazol-5-yl)amino)-2-oxoethyl methylsulfamate (**7c**).* Step-1: 2- bromothiazole-5-carboxylic acid (5.0 g, 24.03 mmol, 1.0 eq.) and Cs2CO3 (15.62 g, 48.06 mmol, 2.0 eq.) were weighed into a two necked flask precharged with argon and dissolved in a degassed 4:1 mixture of 1,4-dioxane and H2O (50 mL). 2-ethoxyphenylboronic acid (5.98 g, 36.05 mmol, 1.5 eq.) and Pd(dppf)Cl2•CH2Cl2 (1.96 g, 2.4 mmol, 0.1 eq.) were added and the mixture was heated at 100 °C for 12 h. On completion, the reaction mixture was filtered, concentrated and washed with EtOAc–H2O. Combined organic layers were dried over anhydrous Na2SO4, filtered, concentrated, and the crude product was purified using automated column chromatography (eluent 5% MeOH/CH2Cl2) to afford 2-(2-ethoxyphenyl)thiazole-5-carboxylic acid (**28**) as a brown solid (1.45 g, 20% yield). LCMS: *m/z* = 250.0 [M+H] ^+^. Step-2: to a stirred solution of 2-(2-ethoxyphenyl)thiazole-5-carboxylic acid (**28**, 1.0 g, 4.01 mmol, 1.0 eq.) in *tert*-butyl alcohol (15 mL), diphenylphosphoryl azide (1.65 g, 6.01 mmol, 1.5 eq.) and triethylamine (16.7 mL, 12.03 mmol) were added and the reaction was heated at 80 °C for 10 h. On completion, the reaction mixture was cooled at room temperature and solvent removed in vacuo. The crude was diluted with EtOAc and washed with water. Combined organic layers were dried over anhydrous Na2SO4, filtered, concentrated, and the crude was purified using automated column chromatography (eluent 30% EtOAc/hexane) to afford *tert*- butyl (2-(2-ethoxyphenyl)thiazol-5-yl)carbamate as a yellow solid (0.36 g, 28% yield). LCMS: *m/z* = 321.3 [M+1] ^+^. Step-3: to a stirring solution of *tert*-butyl (2-(2-ethoxyphenyl)thiazol-5-yl)carbamate (0.35 g, 1.09 mmol, 1.0 eq.) in CH2Cl2 (3.5 mL) was added TFA (1.0 mL) and the reaction was stirred at 25 °C for 2 h. On completion, the precipitated solid was filtered to obtain 2-(2-ethoxyphenyl)thiazol-5-amine (**20**) as a white solid (0.02 g, 83% yield). LCMS: *m/z* = 220.8 [M–H]^+^. Step-4: Preparation according to General Procedure-II using 2-(2-ethoxyphenyl)thiazol-5-amine (**20**, 0.2 g, 0.91 mmol, 1.0 eq.) and 2-((*N*-methylsulfamoyl)oxy)acetic acid (**17**, 0.2 g, 0.91 mmol, 1.0 eq.) in T3P (2.0 mL) at 70 °C for 2 h. The product was purified by preparative HPLC to afford **7c** as a white solid (0.105 g, 31% yield, m.p. 178 °C). HPLC purity >99%. ^1^H NMR (401 MHz, DMSO-*d*6): *δ* 11.55 (s, 1H), 8.22 (dd, *J* = 7.9, 1.8 Hz, 1H), 8.02 (s, 1H), 7.75 (s, 1H), 7.38 (ddd, *J* = 8.3, 7.2, 1.8 Hz, 1H), 7.19 (dd, *J* = 8.5, 1.1 Hz, 1H), 7.06 (ddd, *J* = 8.1, 7.3, 1.1 Hz, 1H), 4.72 (s, 2H), 4.29 (q, *J* = 7.0 Hz, 2H), 2.64 (s, 3H), 1.51 (t, *J* = 7.0 Hz, 3H). ^13^C NMR (101 MHz, DMSO-*d*6): *δ* 162.9, 154.6, 153.4, 135.9, 130.1, 128.2, 126.9, 122.2, 120.9, 112.9, 66.3, 64.6, 28.8, 14.7. HRMS (ESI) *m/z*: [M+H]^+^ calculated for C14H18N3O5S2: 372.0688, found 372.0674.

*2-((4-cyano-5-(2-ethoxyphenyl)-1H-pyrazol-3-yl)amino)-2-oxoethyl methylsulfamate (**7d**).* Step-1: to a stirring solution of 3-amino-1*H*-pyrazole-4-carbonitrile (1.0 g, 9.2 mmol, 1.0 eq.) in DMF, *N*- Bromosuccinimide (1.6 g, 9.2 mmol, 1.0 eq.) was added and the reaction stirred at 25 °C for 2 h. On completion, the reaction mixture was poured into sodium bicarbonate solution and extracted with EtOAc. The combined organic layers were washed with brine, dried over anhydrous sodium sulfate, filtered, concentrated, and the obtained crude was purified using automated column chromatography (eluent 40% EtOAc/hexane) to afford 3-amino-5-bromo-1*H*-pyrazole-4-carbonitrile as a brown solid (1.6 g, 47% yield). LCMS: *m/z* = 185.0 [M–2] ^+^. Step-2: to a stirred solution of 3-amino-5-bromo-1*H*-pyrazole-4-carbonitrile (1.0 g, 5.3 mmol, 1.0 eq.) in DME were added 2-ethoxyphenylboronic acid (1.06 g, 6.4 mmol, 1.2 eq.), 2M K2CO3 in water, and purged with argon for 5 minutes. Tetrakis(triphenylphosphine)palladium(0) (0.617 g, 0.5 mmol, 0.1 eq.) was added and the reaction was stirred at 130 °C for 3 h. On completion, the reaction mixture was poured into water and extracted with EtOAc. The combined organic layers were washed with brine, dried over anhydrous sodium sulfate, filtered, concentrated, and the crude product was purified using automated column chromatography (eluent 40% EtOAc/hexane) to afford 3-amino-5-(2-ethoxyphenyl)-1*H*-pyrazole-4-carbonitrile (**21**) as a yellow solid (0.5 g, 41% yield). LCMS: *m/z* = 229.2 [M+H] ^+^. Step-3: preparation according to General Procedure-II using 3-amino-5-(2-ethoxyphenyl)-1*H*-pyrazole-4- carbonitrile (**21**, 0.2 g, 0.87 mmol, 1.0 eq.) and 2-((*N*-methylsulfamoyl)oxy)acetic acid (**17**, 0.148 g, 0.87 mmol, 1.0 eq.) in T3P (2.0 mL) at 70 °C for 16 h. The product was purified using automated column chromatography (eluent 55% EtOAc/hexane), followed by preparative HPLC to afford **7d** as a white solid (0.040 g, 12% yield, m.p. 157 °C). HPLC purity >98%. ^1^H NMR (401 MHz, DMSO-*d*6): *δ* 13.53 (s, 1H), 10.65 (s, 1H), 7.95 (d, *J* = 5.1 Hz, 1H), 7.51 (dd, *J* = 9.5, 6.9 Hz, 2H), 7.21 (d, *J* = 8.2 Hz, 1H), 7.14 – 7.07 (m, 1H), 4.71 (s, 2H), 4.13 (q, *J* = 6.9 Hz, 2H), 2.65 (d, *J* = 4.8 Hz, 3H), 1.36 (t, *J* = 6.9 Hz, 3H). ^13^C NMR (101 MHz, DMSO-*d*6): *δ* 164.7, 155.7, 146.9, 144.9, 131.9, 130.0, 120.5, 115.4, 113.6, 112.7, 86.4, 66.0, 63.8, 28.8, 14.2. HRMS (ESI) *m/z*: [M+H]^+^ calculated for C15H18N5O5S: 380.1029, found 380.1019.

*2-(benzo[d]isothiazol-3-ylamino)-2-oxoethyl methylsulfamate (**8a**).* Preparation according to General Procedure-II using benzo[*d*]isothiazol-3-amine (**22**, 0.089 mg, 0.473 mmol, 1.0 eq.) and 2-((*N*-methylsulfamoyl)oxy)acetic acid (**17**, 0.1 mg, 0.591 mmol, 1.0 eq.) in T3P at 70 °C for 4 h. The product was purified using automated column chromatography (eluent 30% EtOAc/hexane) to afford **8a** as a white solid (0.085 g, 46% yield, m.p. 147 °C). HPLC purity >99%. ^1^H NMR (401 MHz, DMSO-*d*6): *δ* 11.08 (s, 1H), 8.17 (d, *J* = 8.3 Hz, 2H), 7.96 (d, *J* = 5.0 Hz, 1H), 7.63 (ddd, *J* = 8.1, 6.9, 1.1 Hz, 1H), 7.51 (ddd, *J* = 8.0, 7.0, 1.1 Hz, 1H), 4.92 (s, 2H), 2.65 (d, *J* = 4.6 Hz, 3H). ^13^C NMR (101 MHz, DMSO-*d*6): *δ* 165.6, 151.9, 151.5, 128.5, 124.9, 124.2, 120.8, 66.6, 28.9. HRMS (ESI) *m/z*: [M+H]^+^ calculated for C10H12N3O4S2: 302.0269, found 302.0261.

*2-(benzo[d]isoxazol-3-ylamino)-2-oxoethyl methylsulfamate (**8b**).* Preparation according to General Procedure-II using benzo[*d*]isoxazol-3-amine (**23**, 0.238 mg, 1.775 mmol, 1.0 eq.) and 2-((*N*-methylsulfamoyl)oxy)acetic acid **17** (0.300 mg, 1.775 mmol, 1.0 eq.) in T3P at 70 °C for 16 h. The product was purified using automated column chromatography (eluent 30% EtOAc/hexane) to afford **8b** as a white solid (0.120 g, 23% yield, m.p. 126 °C). HPLC purity >99%. ^1^H NMR (401 MHz, DMSO-*d*6): *δ* 11.37 (s, 1H), 8.05 – 7.96 (m, 2H), 7.74 – 7.64 (m, 2H), 7.40 (ddd, *J* = 8.1, 6.8, 1.2 Hz, 1H), 4.86 (s, 2H), 2.65 (s, 3H). ^13^C NMR (101 MHz, DMSO-*d*6): *δ* 165.1, 163.0, 152.9, 130.9, 124.1, 123.5, 115.9, 109.9, 66.2, 28.9.

HRMS (ESI) *m/z*: [M+H]^+^ calculated for C10H12N3O5S: 286.0498, found 286.0491.

*2-(isoxazolo[4,5-b]pyridin-3-ylamino)-2-oxoethyl methylsulfamate (**8c**).* Preparation according to General Procedure-II using isoxazolo[4,5-*b*]pyridin-3-amine (**24**, 0.119 mg, 0.887 mmol, 1.0 eq.) and 2-((*N*-methylsulfamoyl)oxy)acetic acid **17** (0.150 mg, 0.887 mmol, 1.0 eq.) in T3P at 70 °C for 16 h. The product was purified using automated column chromatography (eluent 60% EtOAc/hexane) to afford **8c** as a white solid (0.040 g, 15% yield, m.p. 128–130 °C). HPLC purity >98%. ^1^H NMR (401 MHz, DMSO-*d*6): *δ* 11.47 (s, 1H), 8.76 (dd, *J* = 4.4, 1.2 Hz, 1H), 8.26 (dd, *J* = 8.7, 1.2 Hz, 1H), 7.96 (s, 1H), 7.74 (dd, *J* = 8.6, 4.5 Hz, 1H), 4.90 (s, 2H), 2.65 (s, 3H). ^13^C NMR (101 MHz, DMSO-*d*6): *δ* 164.4, 155.3, 152.7, 147.8, 134.7, 125.2, 118.6, 66.3, 28.9. HRMS (ESI) *m/z*: [M+H]^+^ calculated for C9H11N4O5S: 287.0450, found 287.0441.

*2-oxo-2-((5-phenyl-1H-pyrazol-3-yl)amino)ethyl methylsulfamate (**9a**).* Step-1: to a stirred solution of 5-phenyl-1*H*-pyrazol-3-amine **25** (0.800 g, 5.025 mmol, 1.0 eq) in CH2Cl2 were added triethylamine (0.7 mL, 10.05 mol, 2.0 eq) and Boc anhydride (2.19 g, 10.05 mole, 2.0 eq) and the reaction was stirred at 25 °C for 4 h. On completion, the reaction mixture was poured into water and extracted with EtOAc. The organic layers were dried over anhydrous Na2SO4, concentrated in vacuo, and the crude product was purified using automated column chromatography (eluent 25% EtOAc/hexane) to afford *tert*-butyl 3- amino-5-phenyl-1*H*-pyrazole-1-carboxylate (0.500 g, 38% yield). LCMS: *m/z* [M–100]^−^ 160.2. Step-2: preparation according to General Procedure-II using *tert*-butyl 3-amino-5-phenyl-1*H*-pyrazole-1- carboxylate (0.320 g, 1.234 mmol, 1.0 eq.) and 2-((*N*-methylsulfamoyl)oxy)acetic acid (**17**, 0.208 g, 1.234 mmol, 1.5 eq.) in T3P (0.785 g, 1.234 mmol, 1.0 eq.) at 70 °C for 2 h. The product was purified by preparative HPLC to afford **9a** as a white solid (0.014 g, 4% yield, m.p. 135–137 °C). HPLC purity >99%. ^1^H NMR (401 MHz, DMSO-*d*6): *δ* 12.93 (s, 1H), 10.69 (s, 1H), 7.89 (d, *J* = 4.9 Hz, 1H), 7.72 (d, *J* = 7.7 Hz, 2H), 7.45 (t, *J* = 7.6 Hz, 2H), 7.35 (t, *J* = 7.4 Hz, 1H), 6.90 (s, 1H), 4.65 (s, 2H), 2.64 (d, *J* = 4.8 Hz, 3H). ^13^C NMR (101 MHz, DMSO-*d*6): *δ* 163.3, 147.3, 142.0, 128.9 (2C), 128.2, 124.98 (2C), 93.8, 66.3, 28.8. HRMS (ESI) *m/z*: [M+H]^+^ calculated for C12H15N4O4S: 311.0814, found 311.0805.

*2-oxo-2-((3-(4-(prop-2-yn-1-yloxy)phenyl)-1H-pyrazol-5-yl)amino)ethyl methylsulfamate (**9b**).* Step-1: to a stirred solution of methyl 4-hydroxybenzoate (**33**, 10.0 g, 65.78 mmol, 1.0 eq) in DMF (100 mL) were added K2CO3 (13.61 g, 98.68 mmol, 1.0 eq), 4-methoxybenzyl chloride (15.45 g, 98.67 mmol, 1.5 eq.) and the reaction was stirred at 50 °C for 16 h. On completion, the reaction mixture was poured into water and extracted with EtOAc. The organic layers were dried over anhydrous Na2SO4, concentrated in vacuo, and the resulting crude was purified using automated column chromatography (eluent 0–30% EtOAc/hexane) to afford methyl 4-((4-methoxybenzyl)oxy) benzoate (15.0 g, 84% yield). LCMS *m/z* 273.1 [M+1]^+^. Step-2: to a solution of acetonitrile (2.26 g, 55.14 mmol, 1.0 eq) in THF (75 mL) at –78 °C under nitrogen was slowly added *n*-butyl lithium (44.11 mL, 110.29 mmol, 2.0 eq) and the reaction was stirred at –78 °C for 30 min. A solution of methyl 4-((4-methoxybenzyl)oxy)benzoate (15.0 g, 55.14 mmol, 1.0 eq) in THF (75 mL) was added slowly to the reaction mixture and the reaction was stirred at 25 °C for 3 h. On completion, the reaction mixture was quenched with saturated NH4Cl solution and extracted with EtOAc. The organic layers were dried over anhydrous Na2SO4, concentrated in vacuo, and the resulting crude was purified using automated column chromatography (eluent 0–30% EtOAc/hexane) to afford 3-(4-((4- methoxybenzyl) oxy) phenyl)-3-oxopropanenitrile (10.0 g, 65% yield). LCMS: *m/z* = 280 [M–H]^−^. Step-3: to a stirring solution of 3-(4-((4-methoxybenzyl) oxy) phenyl)-3-oxopropanenitrile (10.0 g, 35.58 mmol, 1.0 eq) in ethanol (100 mL), hydrazine hydrate (1.14 mL, 35.58 mmol, 1.0 eq.) was added and the reaction stirred at 90 °C for 16 h. On completion, the reaction mixture was concentrated in vacuo and the crude product was purified using automated column chromatography (80% EtOAc/hexane) to furnish 5-(4-((4- methoxybenzyl)oxy)phenyl)-1*H*-pyrazol-3-amine (**32**) as a white solid (4.2 g, 40% yield). LCMS: *m/z* = 296.2 [M+H]^+^. Step-4: to a stirring solution of 5-(4-((4-methoxybenzyl)oxy)phenyl)-1*H*-pyrazol-3-amine (**32**) in CH2Cl2 (50 mL) were added TEA (6.9 mL, 42.71 mmol, 3.0 eq.) and DMAP (0.347 g, 2.847 mmol, 0.2 eq.). After 10 min, Boc anhydride (4.75 mL, 21.35 mmol, 1.5 eq.) was added and the reaction stirred at 25 °C for 12 h. On completion, the reaction mixture was poured into water and extracted with EtOAc. The combined organic layers were washed with brine, dried over anhydrous Na2SO4, and concentrated in vacuo to afford *tert*-butyl 3-(bis(*tert*-butoxycarbonyl)amino)-5-(4-((4-methoxybenzyl)oxy)phenyl)-1*H*-pyrazole- 1-carboxylate as a white solid (2.0 g, 29% yield). LCMS: *m/z* = 596.5 [M+H]^+^. Step-5: to a stirred solution of *tert*-butyl 3-(bis(*tert*-butoxycarbonyl)amino)-5-(4-((4-methoxybenzyl)oxy)phenyl)-1*H*-pyrazole-1- carboxylate (2.0 g, 3.36 mmol, 1.0 eq) in MeOH (20 mL) was added 5% Pd/C (1.0 g) under H2 gas. The reaction was stirred at 25 °C for 16 h. On completion, the reaction mixture was filtered through celite, washed with MeOH, and concentrated in vacuo to afford *tert*-butyl 3-(bis(*tert*-butoxycarbonyl)amino)-5- (4-hydroxyphenyl)-1*H*-pyrazole-1-carboxylate (**33**, white solid, 1.5 g, 99% yield). LCMS: *m/z* = 476.3 [M+H]^+^. Step-6: to a solution of (**33**, 1.5 g, 3.16 mmol, 1.0 eq) in DMF were added propargyl bromide (0.45 g, 3.79 mmol, 1.2 eq) and K2CO3 (0.65 g, 4.74 mmol, 1.5 eq) and the reaction was stirred at 60 °C for 3 h. On completion, the reaction mixture was diluted with EtOAc and washed with saturated aqueous NaHCO3 solution and brine. Combined organic layers were dried over anhydrous Na2SO4, filtered, concentrated, and purified by automated column chromatography to afford *tert*-butyl 3-(bis(*tert*- butoxycarbonyl)amino)-5-(4-(prop-2-yn-1-yloxy)phenyl)-1*H*-pyrazole-1-carboxylate (white solid, 0.7 g, 43.2%). LCMS: *m/z* = 514.3 [M+H]^+^. Step-7: boc-deprotection of *tert*-butyl 3-(bis(*tert*- butoxycarbonyl)amino)-5-(4-(prop-2-yn-1-yloxy)phenyl)-1*H*-pyrazole-1-carboxylate (0.7 g, 1.36 mmol, 1.0 eq) using TFA (0.25 mL) in CH2Cl2 (10 mL) afforded 5-(4-(prop-2-yn-1-yloxy)phenyl)-1*H*-pyrazol-3- amine (**26**) as a white solid (0.37 g). LCMS: *m/z* = 214.1 [M+H]^+^. Step-8: preparation according to General Procedure-II using 5-(4-(prop-2-yn-1-yloxy)phenyl)-1*H*-pyrazol-3-amine (**26**, 0.15 g, 0.70 mmol, 1.0 eq.) and 2-((*N*-methylsulfamoyl)oxy)acetic acid (**17**, 0.240 g, 1.42 mmol, 1.0 eq.) in T3P (2.0 mL) at 70 °C for 16 h. The product was purified by preparative HPLC to afford **9b** as a white solid (0.02 g, 8% yield, m.p. 146–148 °C). HPLC purity >97%. ^1^H NMR (400 MHz, DMSO-*d*6): *δ* 10.66 (s, 1H), 7.89 (d, *J* = 5.0 Hz, 1H), 7.67 (d, *J* = 8.4 Hz, 2H), 7.06 (d, *J* = 8.5 Hz, 2H), 6.79 (s, 1H), 4.84 (d, *J* = 2.4 Hz, 2H), 4.64 (s, 2H), 3.60 (t, *J* = 2.4 Hz, 1H), 2.64 (d, *J* = 4.8 Hz, 3H). ^13^C NMR (214 MHz, DMSO-*d*6): *δ* 163.3, 158.1, 157.9, 157.1, 126.4 (2C), 115.3 (2C), 93.1, 79.1, 78.4, 66.3, 55.5, 28.8. HRMS (ESI) *m/z*: [M+H]^+^ calculated for C15H17N4O5S: 365.0920, found 365.0911.

*2-((5-(4-ethynylphenyl)-1H-pyrazol-3-yl)amino)-2-oxoethyl methylsulfamate (**9c**).* Step-1: to a stirred solution of methyl 4-iodobenzoate (25 g, 90.42 mmol, 1.0 eq) in DMF (250 mL) were added triethylamine (40 mL, 286.257 mmol, 3 eq), PdCl2(PPh3)2 (6.69 g, 9.54 mmol, 0.1 eq), CuI (1.8 g, 9.54 mmol, 0.1 eq), and TMS (27.1 mL, 190.84 mmol, 2.0 eq). The reaction mixture was stirred at 25 °C for 1 h. On completion, the reaction mixture was filtered, washed with water, and the organic layer was extracted with ethyl acetate. Combined organic layers were dried over anhydrous Na2SO4, filtered, concentrated, and the crude product was purified by column chromatography (eluent: 5% EtOAc/hexanes) to afford methyl 4-((trimethylsilyl)ethynyl)benzoate as a brown solid (22 g, 99% yield). Step-2: to a stirred solution of acetonitrile (6.4 mL, 121.767 mmol, 2.5 eq.) in THF (120 mL) was added *n*-BuLi (2.5 M in hexane, 48.7 mL, 121.77 mmol, 2.5 eq) dropwise at –78 °C and stirred for 30 min. Methyl 4- ((trimethylsilyl)ethynyl)benzoate (11.3 g, 48.71 mmol, 1.0 eq) was added at –78 °C and the reaction was stirred for 1 h. On completion, the reaction mixture was quenched with 1N HCl solution till pH∼3. Organic layer was extracted with EtOAc, dried over anhydrous Na2SO4, concentrated, and the resulting crude was purified by column chromatography (eluent 10% EtOAc/hexanes) to afford 3-oxo-3-(4- ((trimethylsilyl)ethynyl)phenyl)propanenitrile as a colorless oil, 7 g, 59% yield). LCMS: *m/z* = 240.1 [M– H]^+^. Step-3: to a stirred solution of 3-oxo-3-(4-((trimethylsilyl)ethynyl)phenyl)propanenitrile (14 g, 58.09 mmol, 1.0 eq) in MeOH (140 mL) was added K2CO3 (16 g, 116.18 mmol, 2.0 eq) and the reaction was stirred at 25 °C for 2 h. On completion, the reaction mixture was quenched with 1N HCl solution till pH∼3 and extracted with EtOAc. Combined organic layers were dried over anhydrous Na2SO4, filtered, concentrated, and the crude product was purified by column chromatography (eluent 10% EtOAc/hexanes) to afford 3-(4-ethynylphenyl)-3-oxopropanenitrile as a colorless oil (9.5 g, 96% yield). LCMS: *m/z* = 168 [M–H]^+^. Step-4: to a stirred solution of 3-(4-ethynylphenyl)-3-oxopropanenitrile (4.5 g, 26.63 mmol, 1.0 eq.) in IPA (45 mL) was added hydrazine hydrate (6.65 g, 133.14 mmol, 5.0 eq.) and the reaction stirred at 80 °C for 6 h. On completion, the reaction mixture was concentrated in vacuo and the crude product was purified using automated column chromatography (eluent 10% EtOAc/hexane) to afford 5-(4- ethynylphenyl)-1*H*-pyrazol-3-amine (**27**, 2.8 g, 58% yield) was obtained as a white solid. LCMS: *m/z* = 184.1 [M+H]^+^. Step-5: preparation according to General Procedure-II using 5-(4-ethynylphenyl)-1*H*- pyrazol-3-amine (**27**, 0.207 g, 1.14 mmol, 0.8 eq) and 2-((*N*-methylsulfamoyl)oxy)acetic acid (**17**, 0.240 g, 1.42 mmol, 1.0 eq.) in T3P (2.0 mL) at 70 °C for 16 h. The product was purified by preparative HPLC to afford **9c** as a white solid (0.040 g, 8% yield, m.p. 154–156 °C). HPLC purity >98%. ^1^H NMR (400 MHz, DMSO-*d*6): *δ* 13.03 (s, 1H), 10.74 (s, 1H), 7.91 (d, *J* = 4.2 Hz, 1H), 7.75 (d, *J* = 8.0 Hz, 2H), 7.54 (d, *J* = 8.1 Hz, 2H), 6.92 (s, 1H), 4.65 (s, 2H), 4.29 (s, 1H), 2.64 (d, *J* = 4.7 Hz, 3H). ^13^C NMR (214 MHz, DMSO- *d*6): 163.4, 147.6, 141.9, 132.3 (2C), 129.7, 125.1 (2C), 121.2, 94.2, 83.2, 81.8, 66.3, 28.8. HRMS (ESI) *m/z*: [M+H]^+^ calculated for C14H15N4O4S: 335.0814, found 335.0805.

### Biological assays

The CHIKV nsP2pro, CHIKV-nLuc viral replication, VEEV-nLuc viral replication, and alphavirus viral titer reduction (VTR), assays were performed using the reported protocols.^10^

### GSH Capture Assay

The GSH reactivity of compounds **1**–**5**, **6a**–**d** was determined using the reported protocol.^12^

### Enzyme Binding Kinetics

*K*i and *k*inact were determined by combining different concentrations of an internally quenched protease substrate with a *N*-terminal CyLyte Fluor5 and C-terminal QXL^®^670 (Anaspec) with compound **5** diluted in a 2-fold 16-point dose response starting from 100 µM in DMSO. 150 nM of CHIKV nsP2 was added to this mixture and relative fluorescence was measured immediately following addition every minute for 30 min. Relative percent activity was determined by measuring the relative activity (RFU/min) at each timepoint and comparing it to the relative activity of DMSO treated sample at the given timepoint. *k*obs, *K*i, and *k*inact were determined using GraphPrism. Relative percent activity was plotted over time and fit to a one-phase decay non-linear regression to calculate *K*obs for each concentration and *k*inact and *K*i were determined by plotting *k*obs over compound concentration fit to an irreversible inhibition non-linear regression where Relative Percent Activity = [Experimental RFU/min Time(x)/DMSO RFU/min Time(x)]*100.

### Cysteine Protease Selectivity Assay

Cysteine protease inhibition assays were performed in duplicate at BPS Bioscience Inc., San Diego, CA. A solution of compound **5** ten-fold higher than the final concentration was prepared with 10% DMSO in assay buffer and 5 µL of the dilution was added to a 50 µL reaction so that the final concentration of DMSO was 1% in all the reactions. All control samples, including background and no compound controls, also contain 1% DMSO. The compounds were pre-incubated in duplicate at room temperature for 30 min in a mixture containing assay buffer, enzyme. For cathepsin L assay, the enzymatic reactions were conducted in duplicate at room temperature for 30 min in a 100 µL mixture containing 50 mM MES buffer, pH 5.0, 100 mM NaCl, 5 mM DTT, cathepsin substrate, cathepsin enzyme, and compound **5**. Fluorescence intensity was measured at an excitation of 360 nm and an emission of 460 nm using a Tecan Infinite M1000 microplate reader. For deubiquitinase assay, the enzymatic reactions were conducted in duplicate at room temperature for 30 min in a 50 µL mixture containing 50 mM Tris-HCl, pH 7.4, 0.5 mM EDTA, 0.05% Tween 20, 1 mM DTT, 100 nM Ubiquitin-AMC substrate, a deubiquitinase enzyme, and compound **5**. Fluorescence intensity was measured at an excitation of 360 nm and an emission of 460 nm using a Tecan Infinite M1000 microplate reader. For 3CLPro (SARS-CoV- 2) assay, the mixtures were incubated for 30 min at room temperature with slow shaking. Enzymatic reactions were initiated by addition of 10 μL of 200 μM substrate to the mixtures and proceeded at room temperature with slow shaking and protected from light. Fluorescence intensity was measured using a M1000 Tecan microplate reader. The fluorescent intensity data were analyzed using the computer software, GraphPad Prism. In absence of compound **5**, the fluorescent intensity (Ft) in each data set was defined as 100% activity. In the absence of the enzyme, the fluorescent intensity (Fb) in each data set was defined as 0% activity. The percent activity in the presence of each compound was calculated according to the following equation: % activity = (F-Fb)/(Ft–Fb), where F = the fluorescent intensity in the presence of compound **5**.

### Fluorescence-based Chemoproteomics

HEK293 cells were lysed by sonication and the resulting lysates (∼3.5 mg/mL) diluted in reaction buffer (25 mM HEPES pH 7.3, 1 mM DTT, 1% DMSO). 2 µL of a 250 µM stock solution in DMSO of clickable sulfamate (**NMS**) or clickable chloroacetamide (**CA**) were added to 40 µL of cell lysate (∼140 µg total protein) and the mixtures were incubated at 24 °C for 30 min. Equal volumes of THPTA (5 mM stock in H2O) and CuSO4.H2O (10 mM stock in H2O) were premixed to generate the ligand-copper solution. For the click reaction, freshly prepared solutions of 10 µL 6-TAMRA azide purchased from Vector Laboratories, cat. CCT-1246 (50 µM stock in buffer), 20 µL ligand-copper premix, and 10 µL sodium ascorbate (10 mM stock in H2O) were added successively to cell lysates, and the mixture was incubated in the dark at 24 °C for 1 h. On completion, 45 µL of the reaction was mixed with 45 µL of SDS-PAGE sample buffer [1:5 volumes of 10x reducing agent (NuPAGE; cat. NP0009) to 2X SDS sample buffer (Novex, cat. LC2676)] and heated at 85 °C for 5 min. Samples were separated in a 4–12% Tris-Glycine SDS-PAGE (Novex, catalog number XP04125BOX). The TAMRA labeling was visualized on an Invitrogen iBrightFL1000 (515–545 nm excitation; 568–617 nm emission). Total protein was visualized following Coomassie staining. For reactions containing **VSC**, the competitor compound was added to the cell lysate (2 µL of a 240 µM stock solution in DMSO, final concentration 10 µM) and incubated at 24 °C for 30 min prior to the addition of **NMS** or **CA**. For reactions containing nsP2 protease, 4 µg of the purified protein was added to the cell lysate prior to the addition of any compound.

### Hepatocyte Stability Assay

Cryopreserved hepatocytes from mouse (TPCS, cat. no. CMH- 100CD-SQ, lot no. CMH100CD-V01422 and CMH100CD-V01699, pooled male CD-1) and human (BioIVT, cat. no. X008001, lot no. QZW and YIT, 10 pooled donors each) were used to assess compound stability. 10 mM stock solutions of compound **5** and positive control verapamil were prepared in DMSO. Thawing medium and supplement incubation medium (serum-free) were placed in a 37 °C water bath for at least 15 min prior to use. Stock solutions were diluted to 100 μM by combining 198 μL of 1:1 MeCN/water and 2 μL of 10 mM stock solution. Vials of cryopreserved hepatocytes were removed from cryogenic storage and thawed in a 37 °C water bath with gentle shaking until all ice crystals had dissolved. The contents were poured into a 50 mL thawing medium conical tube and centrifuged at 100 g for 10 min at room temperature. Thawing medium was aspirated, and hepatocytes were resuspended with serum-free incubation medium to yield ∼1.5 × 10^6^ cells/mL. Cell viability and density were counted using AO/PI fluorescence staining and then diluted with serum-free incubation medium to a working cell density of 0.5×10^6^ viable cells/mL. A portion of the hepatocytes at 0.5×10^6^ viable cells/mL was boiled for 5 min prior to adding to the plate as negative control. Aliquots of 198 μL hepatocytes were dispensed into each well of a 96-well non-coated plate. For the inhibitor plate, 0.5 μL of 80 mM EA (0.2 mM) or 400 mM 1-ABT (1 mM) were added. The plate was placed in the incubator for approximately 20 min. Aliquots of 2 μL of 100 μM compound **5** (1 μM) and positive control were added into respective wells of the non-coated 96-well plate to start the reaction. The assay was performed in duplicate. The plate was incubated for the designated time points; subsequently, 25 μL of contents were transferred and mixed with 6 volumes (150 μL) of cold MeCN with IS (100 nM alprazolam, 200 nM labetalol, 200 nM caffeine and 200 nM diclofenac) to terminate the reaction at time points of 0, 15, 30, 60, 90 and 120 min. Samples were centrifuged for 45 min at 3,220 g and an aliquot of 100 µL of the supernatant was diluted by 100 µL ultra-pure H2O, and the mixture was used for LC-MS/MS analysis. Peak areas of compound **5** were determined from extracted ion chromatograms. The *in vitro* t1/2 was determined from the slope value using the relationship: *in vitro* t1/2 = 0.693/k, where k is the rate constant. The intrinsic clearance in μL/min/10^6^ cells was calculated using the relationship CLint = kV/N where V is the incubation volume (0.2 mL) and N is the number of hepatocytes per well (0.1 × 10^6^ cells).

### Metabolite Identification

Compound **5** was incubated with cryopreserved male CD-1 mouse hepatocytes (1.0 × 10⁶ cells/mL) in the presence of 1 mM 1-ABT to inhibit cytochrome P450-mediated metabolism. Incubations were conducted at 37 °C for 0, 15, and 30 min. Incubations were quenched with 2 volumes of acetonitrile (0.1% formic acid), followed by centrifugation for 30 min at 16,000 g. 400 µL supernatants were then transferred to clean tubes and place in the evaporator under steady stream of nitrogen at room temperature until dry. The dried residues were reconstituted with 100 µL of ACN: H2O = 1:4 (v/v) solution and vortexed for proper mixing and injected into LC-UV-MS/MS. Samples for metabolite identification and profiling were analyzed using a Vanquish UHPLC system, equipped with Diode Array Detector HL and Orbitrap Exploris 240 (Thermo Fisher Scientific, USA). A XSelect HSS T3, 100 x 2.1 mm, 2.5 µm HPLC column (Waters) was used for separation of analytes. The mobile phase was a gradient of water, containing 0.1% formic acid (A) and acetonitrile, containing 0.1% formic acid (B). The flow rate was 500 µL/min. The mobile phase composition began at 5% B held for 0–1.5 min, 5–60% B at 1.5–9 min, 60–100% B at 9–12 min, 100% B at 12–14 min, 100–5% B at 14–14.3 min, and finally 14.3–15 min re-equilibration period at 5% B. UV absorbance was monitored with a photodiode array detector from 220– 400 nm. The mass spectrometer was operated in both negative and positive-ion modes with an electrospray ion source potential of 3.5 kV, and source heater temperature of 350 °C. Curtain, GS1 and GS2 gas flow settings were 25, 30, and 35 units, respectively. Full scan mass spectra were acquired over the range 120– 1200 *m/z* at a constant resolving power of ∼ 60,000. Metabolites were characterized by comparison of mass spectral fragmentation patterns with that for compound **5** (see Figure 5 and Figure S5 for structural assignments of metabolites).

### Mouse Liver Microsome Assay with Cofactors

*With NADPH*. Pooled, male CD-1 mice (cat. no. M1000, lot no. 2210246, sponsor: XENOTECH) were used for the assay. For metabolic stability assessment, a master incubation mixture was prepared by combining mouse liver microsomes (20 mg/mL stock, 2 µL; final concentration 0.556 mg/mL) and Alamethicin (5 mg/mL stock, 0.4 µL; final concentration 27.8 µg/mL) with other necessary components as per standard protocol. The final volume of each incubation was adjusted using phosphate buffer and ultra-pure water. The metabolic reaction was initiated by adding NADPH (10 mM, prepared in water) to the pre-warmed mixture and incubated at 37 °C. Reactions were carried out in triplicate and quenched at designated time points (0, 15, 30, 45, and 60 min) using two volumes of ice-cold acetonitrile containing 0.1% formic acid. Samples were vortexed and centrifuged at 16,000 g for 30 min to pellet the proteins. Supernatants (400 µL) were carefully transferred to clean tubes and dried under a gentle stream of nitrogen at room temperature. The dried residues were reconstituted in 100 µL of a 20:80 MeCN/H2O (v/v) solution, vortexed thoroughly, and transferred to autosampler vials for analysis. Chromatographic separation was conducted on a Shimadzu LC system equipped with a column maintained at 40 °C. A sample volume of 1 µL was injected into the system. Mass spectrometric analysis was performed using a SCIEX Triple Quad™ 6500+ instrument operated in appropriate ionization and acquisition modes for targeted quantification of the parent test compound.

*With UDPGA*. Pooled, male CD-1 mice (cat. no. M1000, lot no. 2310181, sponsor: XENOTECH) were used for the assay. A master incubation mixture was prepared in 100 mM phosphate buffer (pH 7.4), containing mouse liver microsomes (20 mg/mL stock, 2 µL; final concentration: 0.556 mg/mL) and Alamethicin (5 mg/mL stock, 0.4 µL; final concentration: 27.8 µg/mL). The microsomal mixture was pre-incubated on ice for 15 min to allow membrane permeabilization by Alamethicin. Uridine 5′-diphospho- glucuronic acid (UDPGA) was freshly prepared at 20 mM concentration in ultra-pure water. The glucuronidation reaction was initiated by the addition of UDPGA to the pre-warmed microsomal mixture. The final reaction volume was adjusted with ultra-pure water. Incubations were conducted at 37 °C for predetermined time points (0, 15, 30, 45, and 60 min), and each reaction was carried out in triplicate. At each time point, reactions were terminated by adding two volumes of ice-cold acetonitrile containing 0.1% formic acid. The quenched mixtures were vortexed and centrifuged at 16,000 g for 30 minutes at 4 °C. A 400 µL aliquot of the supernatant was transferred to a clean tube and dried under a gentle stream of nitrogen at room temperature. The dried residues were reconstituted in 100 µL of 20:80 MeCN/H2O (v/v), vortexed thoroughly, and transferred to autosampler vials. Samples were analyzed using Shimadzu LC30AD systems coupled with SCIEX Triple Quad™ 4500, 5500, or 6500+ mass spectrometers. Chromatographic separation was achieved at 40 °C with an injection volume of 1 µL. Multiple reaction monitoring in negative ion mode was used to detect and quantify the parent test compound.

### Kinetic Solubility Assay

Kinetic solubility of final compounds was determined in triplicate in pH 7.4 phosphate buffer using 10 mM stock solutions at 2.0 % DMSO and CLND detection at Analiza, Inc (Table S1).

## Associated Content

**Supporting Information.** Table S1. Aqueous solubility; Table S2. Inhibitory effects of **5** on human and viral cysteine proteases; Figure S1. Chemical stability of **5** at room temperature; Figure S2. DLS data for **5**; Figure S3. GSH stability of **5**; Figure S4. Cell toxicity data upon 48 h exposure of **5**; Figure S5. MS fragmentation of metabolites M1–13 of **5** from mouse hepatocytes; Figures S6– S45. NMR spectra of final compounds; Figure S46– S66. HPLC analyses (PDF). Molecular formula strings (CSV)

## Acknowledgements

The Structural Genomics Consortium (SGC) is a registered charity (no: 1097737) that receives funds from Bayer AG, Boehringer Ingelheim, Bristol Myers Squibb, Genentech, Genome Canada, through Ontario Genomics Institute [OGI-196], EU/EFPIA/OICR/McGill/KTH/Diamond Innovative Medicines Initiative 2 Joint Under-taking [EUbOPEN grant 875510], Janssen, Merck KGaA (also known as EMD in Canada and the US), Pfizer, and Takeda. We gratefully acknowledge the support of Piramal Pharma Solutions for assistance in the synthesis of intermediates and final compounds and Sumera Perveen (U. Toronto) for compound aggregation determination. The research reported in this publication was supported by NIH grant 1U19AI171292-01 (READDI-AViDD Center), in part by the NC Biotech Center Institutional Support Grant 2018-IDG-1030, and by NIH grant S10OD032476 for upgrading the 500 MHz NMR spectrometer in the UNC Eshelman School of Pharmacy NMR Facility.

## Abbreviations

CHIKV chikungunya virus; DAA directly acting antiviral; GSH, glutathione; GST glutathione *S*- transferase; MAYV Mayaro virus; MLM mouse liver microsome; nsP2 nonstructural protein 2; nLuc nanoluciferase; VEEV, Venezuelan equine encephalitis virus; VTR, viral titer reduction; UGT uridine diphosphate glucuronosyltransferase.

---

**Scheme 1.**
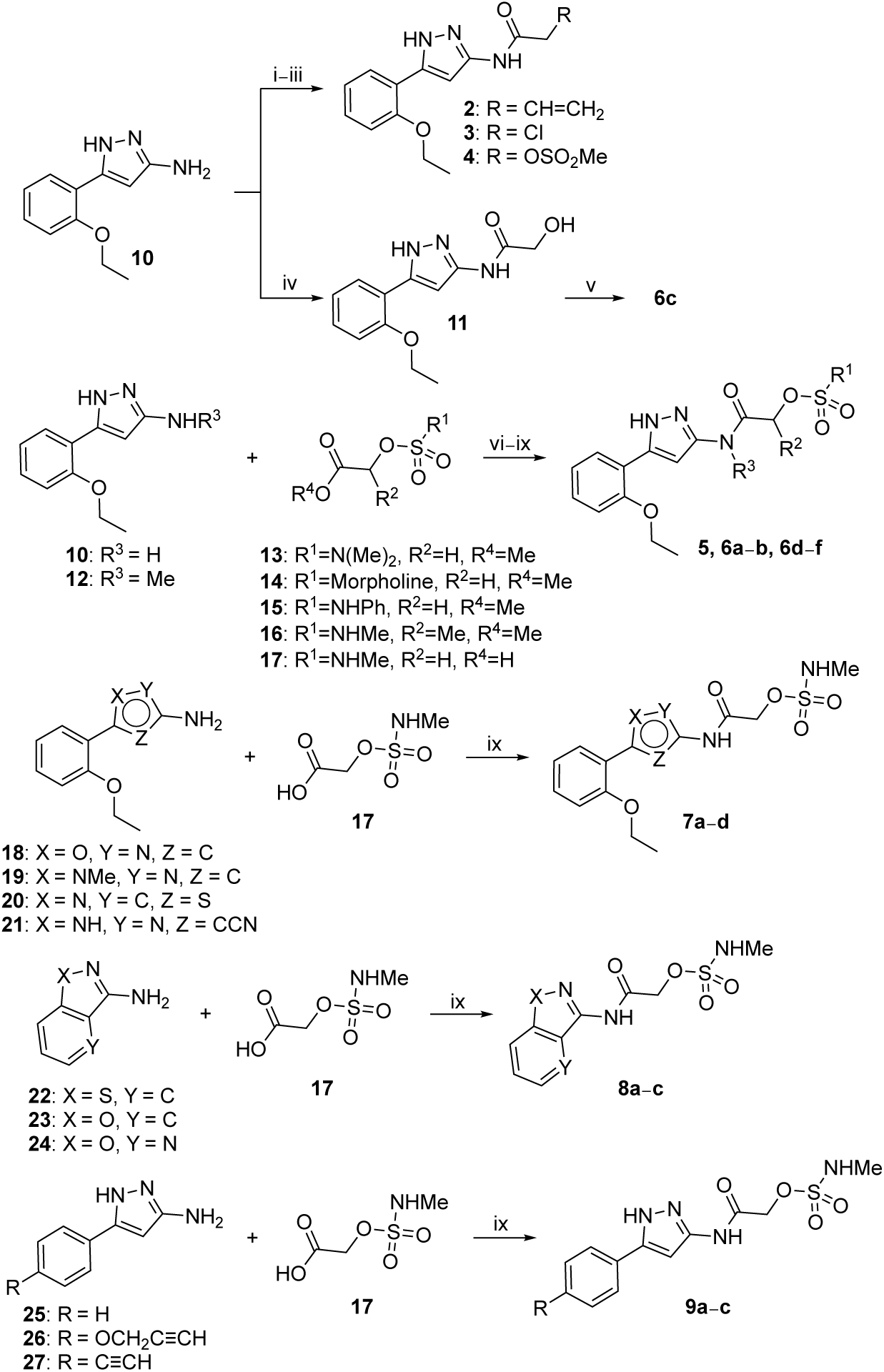
Synthesis of **2**–**5**, **6a**⎼**f**, **7a**⎼**d**, **8a**⎼**c**, **9a**⎼**c***^a a^*Reagents and conditions: (i) acryloyl chloride (for **2**); (ii) 2-chloroacetyl chloride (for **3**); (iii) 2- ((methylsulfonyl)oxy)acetic acid^34^ (for **4**); (iv) methyl 2-hydroxyacetate, triethylaluminum (1.0 M in hexanes); (v) sulfamoyl chloride; (vi) TBTU, pyridine (for **5**); (vii) triazabicyclodecene (for **6a**); (viii) triethylaluminum (1.0 M in hexanes) (for **6b**, **6d**, **6e**); (ix) T3P (for **6f, 7a**⎼**d**, **8a**⎼**c**, **9a**⎼**c**).

**Scheme 2.**
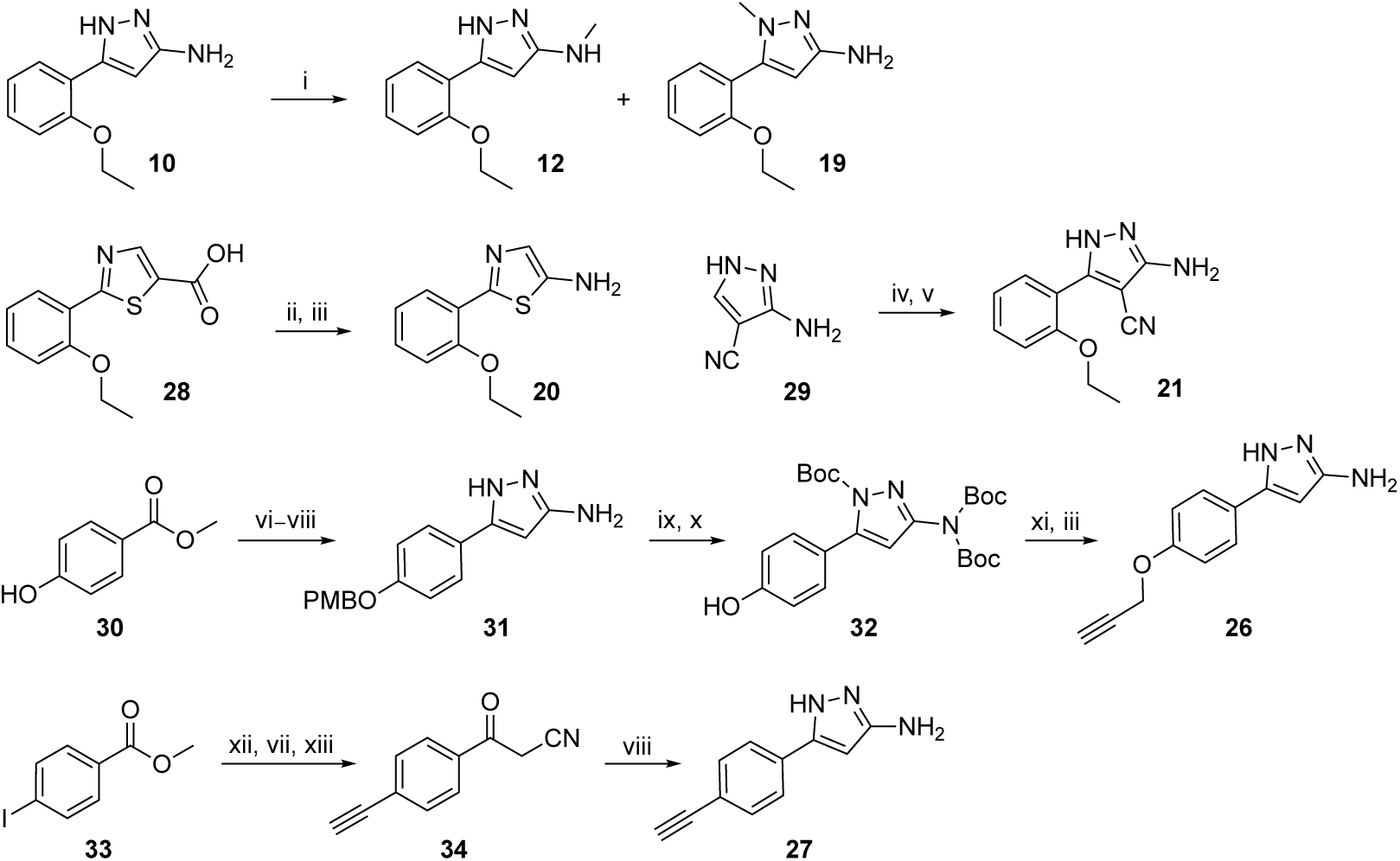
Synthesis of amines **12**, **19**⎼**21**, **26**, **27***^a a^*Reagents and conditions: (i) CH3I; (ii) *t*-BuOH, DPPA; (iii) TFA; (iv) NBS; (v) 2-ethoxyphenylboronic acid; (vi) PMBCl; (vii) ACN, *n-*BuLi; (viii) NH2–NH2.H2O; (ix) (Boc)2O; (x) Pd/C; (xi) propargyl bromide; (xii) trimethylsilylacetylene; (xiii) K2CO3.

**Scheme 3.**
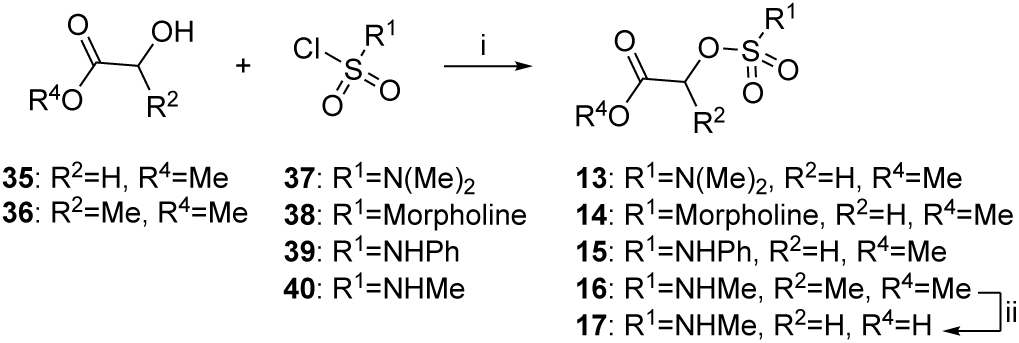
Synthesis of sulfamates **13**–**17***^a a^*Reagents and conditions: (i) dimethylsulfamoyl chloride (for **13**), morpholine-4-sulfonyl chloride (for **14**), phenylsulfamoyl chloride (for **15**), methylsulfamoyl chloride (for **16**, **17**); (ii) LiOH.

