## Supporting Information for "*N*-Alkyl Sulfamates as a New Class of nsP2 Cysteine Protease Inhibitors with Broad Spectrum Antialphaviral Activity"

Table of Contents


| | | Page |
|---|---|---|
| Table S1 | Aqueous solubility | S2 |
| Table S2 | Inhibitory effects of **5** on human and viral cysteine proteases | S3 |
| Figure S1 | Chemical stability of **5** at room temperature | S4 |
| Figure S2 | DLS data for **5** | S5 |
| Figure S3 | GSH stability of **5** | S6 |
| Figure S4 | Cell toxicity data upon 48 h exposure of **5** | S7 |
| Figure S5 | MS fragmentation of metabolites **M1–13** from mouse hepatocytes | S8–S12 |
| Figures S6–S45 | NMR spectra of final compounds | S13–S32 |
| Figures S46–S66 | HPLC analyses | S33–S42 |

**Table S1**. **Aqueous solubility**[a]

| Compound | Aqueous Solubility (μM)[a] |
|---|---|
| **1** | 150 |
| **2** | 50 |
| **3** | 5 |
| **4** | 90 |
| **5** | 70 |
| **6a** | 20 |
| **6b** | 30 |
| **6c** | 140 |
| **6d** | 40 |
| **6e** | 40 |
| **6f** | 190 |
| **7a** | 1 |
| **7b** | 190 |
| **7c** | 1 |
| **7d** | 150 |
| **8a** | 220 |
| **8b** | 220 |
| **8c** | 220 |
| **9a** | 150 |
| **9b** | 150 |
| **9c** | 70 |

[a]Kinetic solubility of a 10 mM DMSO solution at pH 7.4.

**Table S2**. Inhibitory effects of **5** on human and viral cysteine proteases

| Enzyme | Substrate | Control | % Inhibition | | | |
|---|---|---|---|---|---|---|
| | | | **5** (10 µM) | **Reference** (0.1 x $IC_{50}$)[a] | **Reference** (1 x $IC_{50}$)[a] | **Reference** (10 x $IC_{50}$)[a] |
| Cathepsin L | 5 µM Z-Leu-Arg-AMC | E-64 | 14 | 7 | 51 | 96 |
| USP7 | 100 nM Ub-AMC | Ub-Aldehyde | 3 | 2 | 37 | 95 |
| UCHL1 | 100 nM Ub-AMC | Ub-Aldehyde | 8 | 2 | 33 | 92 |
| 3CLPro | 3CL Pro Substrate | GC376 | 8 | 11 | 47 | 95 |

[a]$IC_{50}$ of the control nsP2 protease inhibitor.

**Figure S1**. Chemical stability of **5** at room temperature

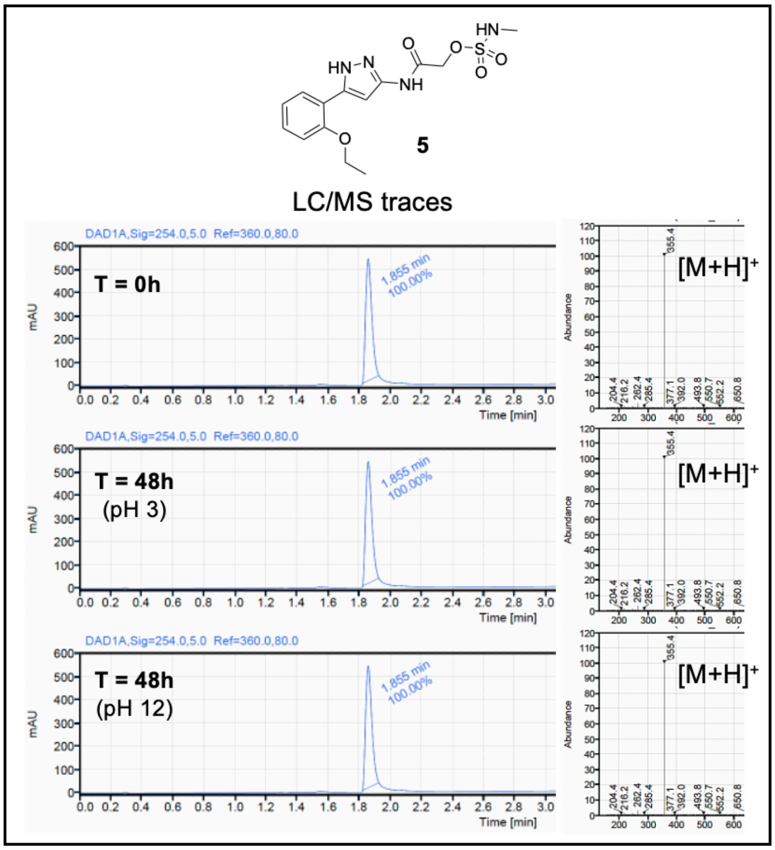

**Figure S2**. DLS data for **5**

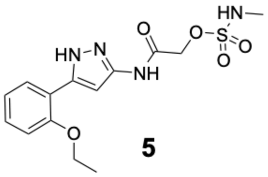


**5**

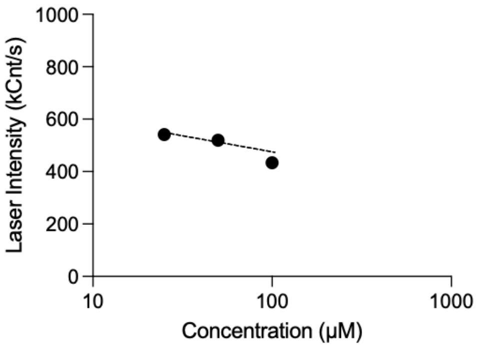


.

**Dynamic Light Scattering (DLS) Assay**. Aggregation behavior of sulfamate **5** was determined by DLS in duplicate at 25 ºC in 25 mM HEPES buffer (pH 7.4) containing 0.003% Tween-20 and 1 mM DTT, using a 10 mM stock solution containing 2% DMSO. Light scattering was measured using a DynaPro Plate Reader III. Buffer with 2% DMSO control produced an average laser intensity of 858 kCnts/s.

**Figure S3**. GSH stability of **5**

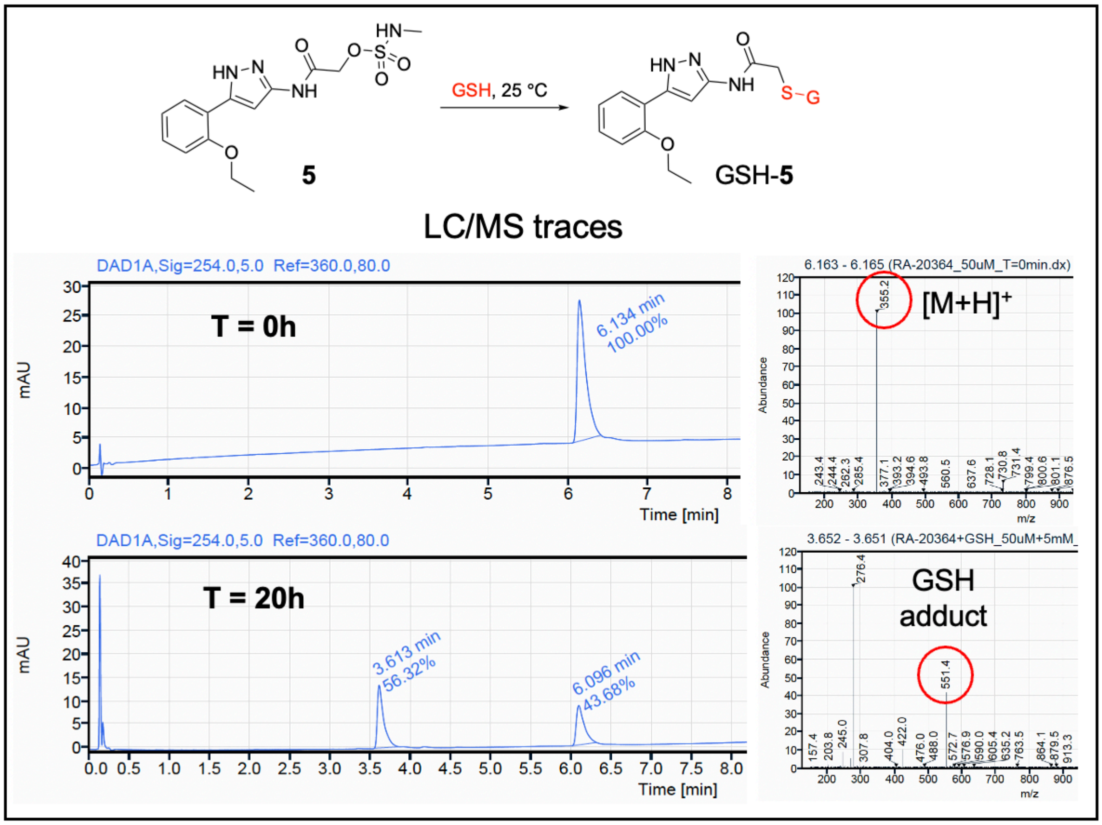


**Method for GSH stability.** A 10 mM solution of GSH (Sigma Aldrich Cat# G4251 (St. Louis, MO, USA)) was prepared in a pH 7.4 phosphate buffer. A 10 mM DMSO solution of Compound **5** was diluted in phosphate buffer to give a solution at 100 µM with 1% DMSO. At time zero (t = 0), 50 µL of the 100 µM test compound solution was added to an Eppendorf tube containing 50 µL phosphate buffer and 50 µL of 10 mM GSH solution. The final concentrations of the compound and GSH were maintained at 50 µM and 5 mM, respectively. The Eppendorf tube was vortexed, and then the sample was transferred to a high-recovery autosampler vial for LCMS analysis. Analysis was performed every 1 h interval, and the percentage of GSH adduct formation was calculated using Agilent LCMS software (OpenLab CDS Version 2.7).

**Figure S4**. Cell toxicity data upon 48 h exposure of **5**

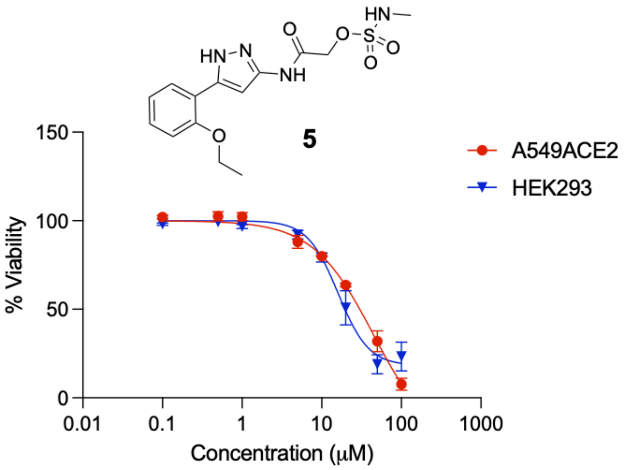


**Cell viability assay**. Cells were plated on a 384 well plate at a seeding density of 2000 cells/well for A549ACE2 (in DMEM high-glucose, 10% fetal bovine serum, 1X non-essential amino acids, and 2 mM L-glutamine) and HEK293 (in DMEM high-glucose, 10% fetal bovine serum) and allowed to adhere overnight. Cells were treated with dose response of compound, 1% DMSO (negative control), or 10% DMSO (positive control) for 48 h. CellTiter-Glo reagent (Promega) was added to each well and luminescence was detected on GloMax plate reader. There were two biological replicates containing treatments in quadruplicate. Cell viability was calculated using the following formula: ((raw RLU value – average of positive control wells) / (average of negative control wells – average of positive control wells))*100. Plots were generated using GraphPad Prism.

**Figure S5**. MS fragmentation of metabolites **M1–13** of **5** from mouse hepatocytes

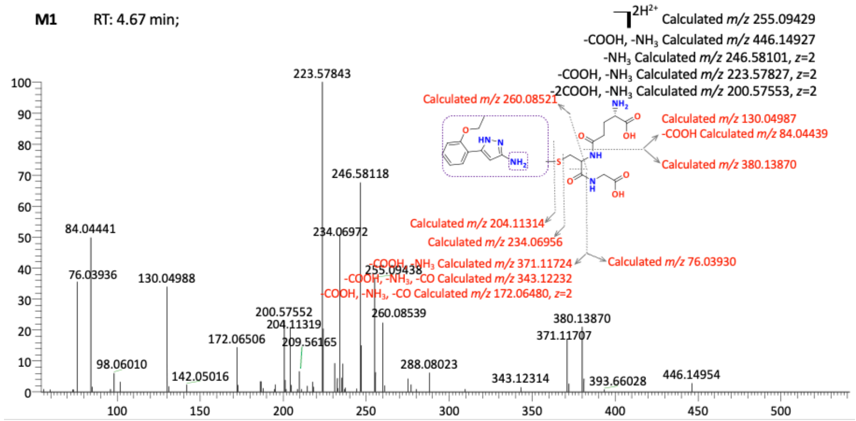


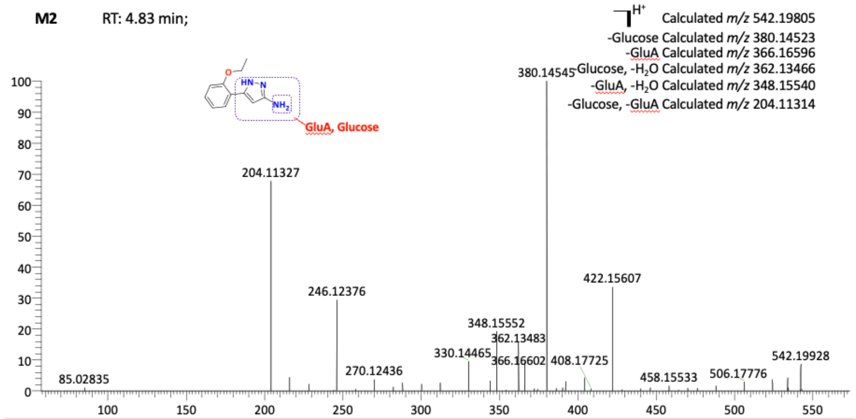


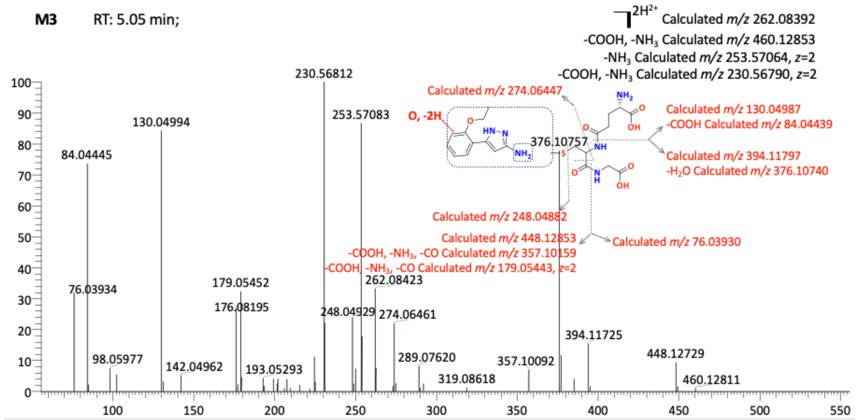

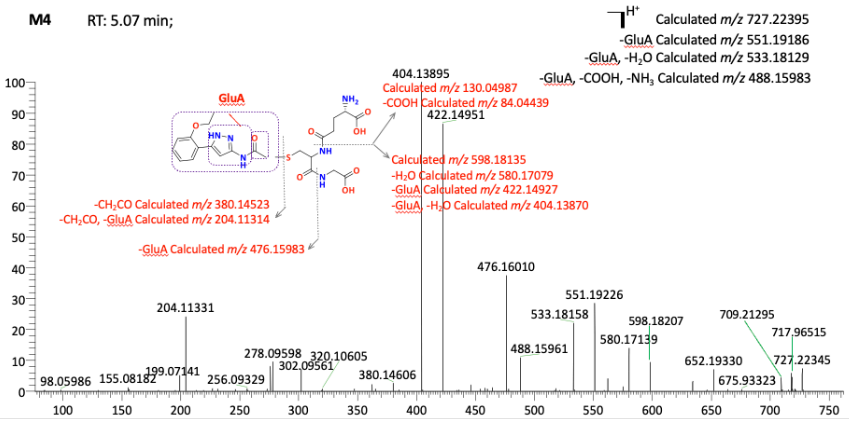
M4
RT: 5.07 min;
H+
Calculated m/z 727.22395
-GluA Calculated m/z 551.19186
-GluA, -H2O Calculated m/z 533.18129
-GluA, -COOH, -NH3 Calculated m/z 488.15983
GluA
Calculated m/z 130.04987
-COOH Calculated m/z 84.04439
Calculated m/z 598.18135
-H2O Calculated m/z 580.17079
-GluA Calculated m/z 422.14927
-GluA, -H2O Calculated m/z 404.13870
-CH2CO Calculated m/z 380.14523
-CH2CO, -GluA Calculated m/z 204.11314
-GluA Calculated m/z 476.15983
404.13895
422.14951
476.16010
551.19226
533.18158
598.18207
709.21295
717.96515
580.17139
488.15961
204.11331
652.19330
727.22345
278.09598
320.10605
302.09561
199.07141
675.93323
380.14606
98.05986
155.08182
256.09329

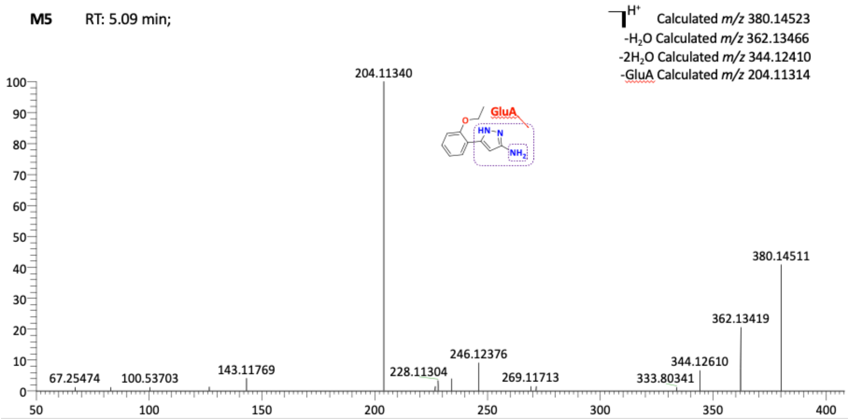
M5
RT: 5.09 min;
H+
Calculated m/z 380.14523
-H2O Calculated m/z 362.13466
-2H2O Calculated m/z 344.12410
-GluA Calculated m/z 204.11314
GluA
204.11340
380.14511
362.13419
246.12376
344.12610
67.25474
100.53703
143.11769
228.11304
269.11713
333.80341

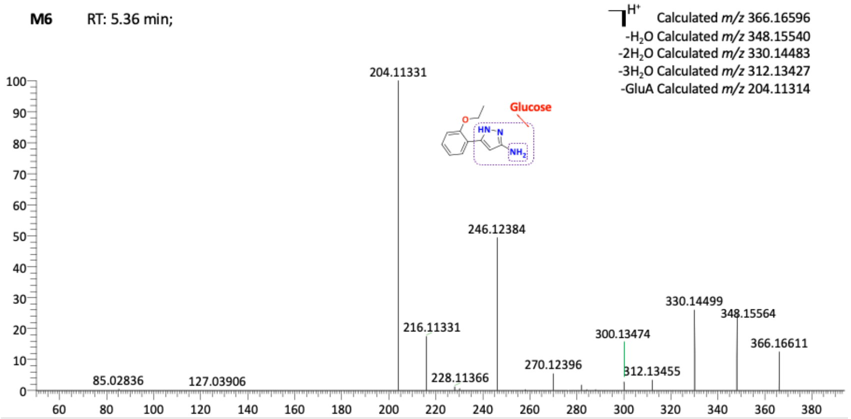
M6
RT: 5.36 min;
H+
Calculated m/z 366.16596
-H2O Calculated m/z 348.15540
-2H2O Calculated m/z 330.14483
-3H2O Calculated m/z 312.13427
-GluA Calculated m/z 204.11314
Glucose
204.11331
246.12384
330.14499
348.15564
366.16611
216.11331
300.13474
270.12396
312.13455
228.11366
85.02836
127.03906

**M7** RT: 5.71 min;

$]^{H^+}$ Calculated *m/z* 503.10785
-GluA Calculated *m/z* 327.07577

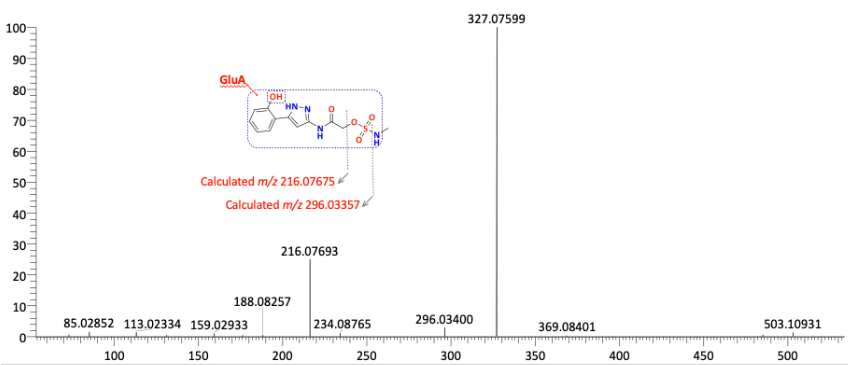


**M8** RT: 5.87 min;

$]^{H^+}$ Calculated *m/z* 204.11314

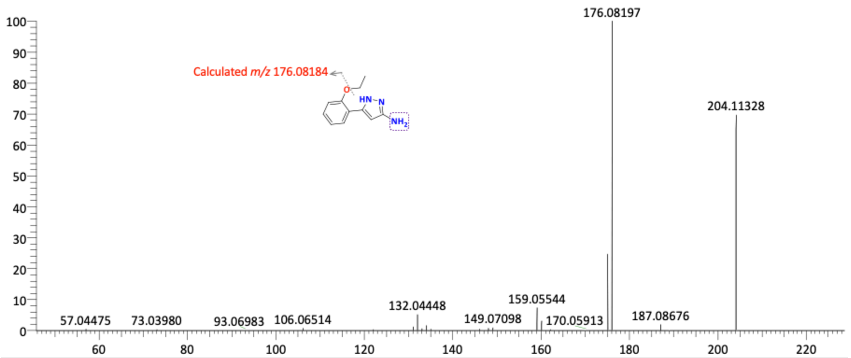


**M9** RT: 5.99 min;

$]^{2H^{2+}}$ Calculated *m/z* 276.09957
$-NH_3$ Calculated *m/z* 267.58629, *z*=2
-COOH, $-NH_3$ Calculated *m/z* 244.58355, *z*=2

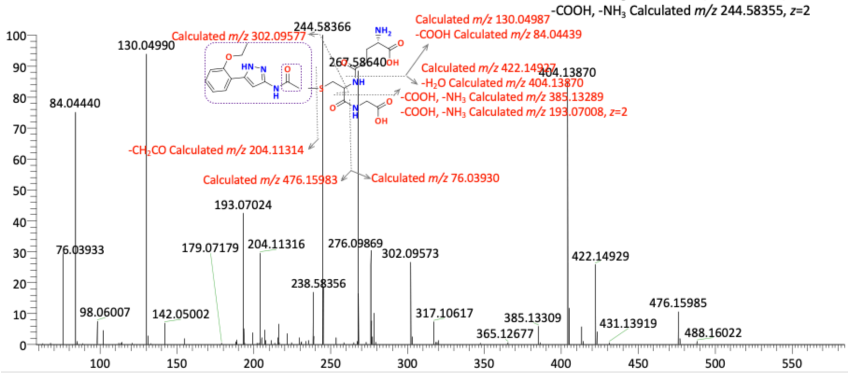

**M10** RT: 6.38 min;

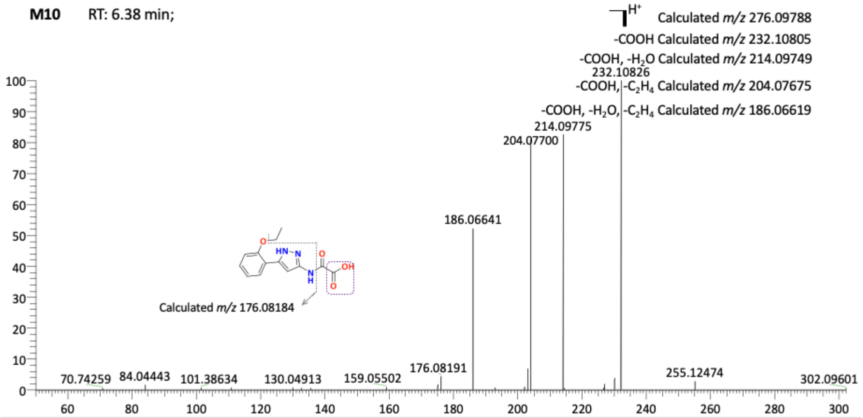


**M11** RT: 6.39 min;

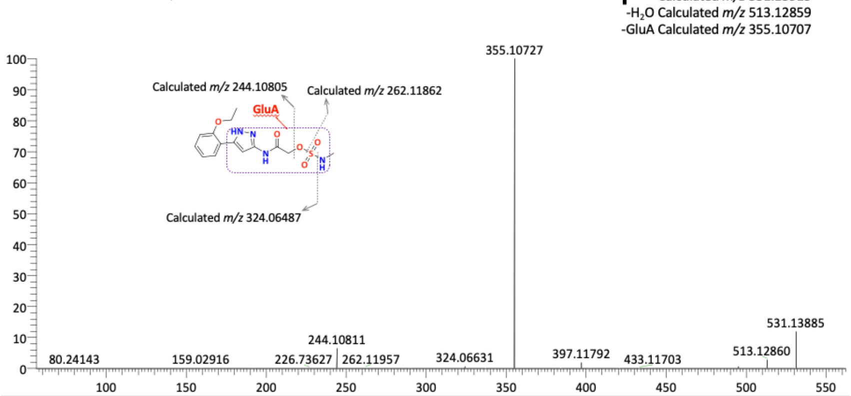


**M12** RT: 7.07 min;

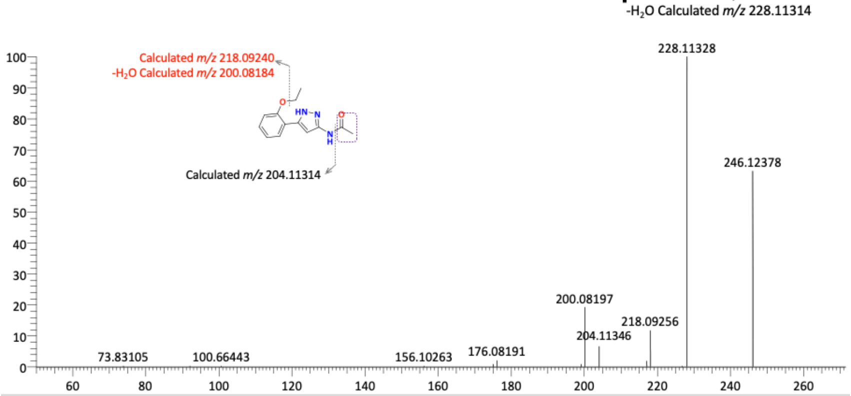

**M13** RT: 7.42 min;

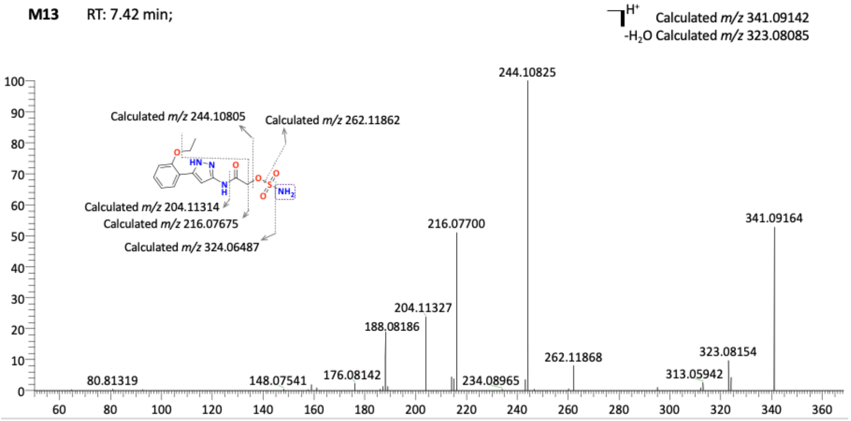

**NMR spectra of final compounds**

**Figure S6**: $^{1}H$ NMR (400 MHz, DMSO-$d_6$) for **2**

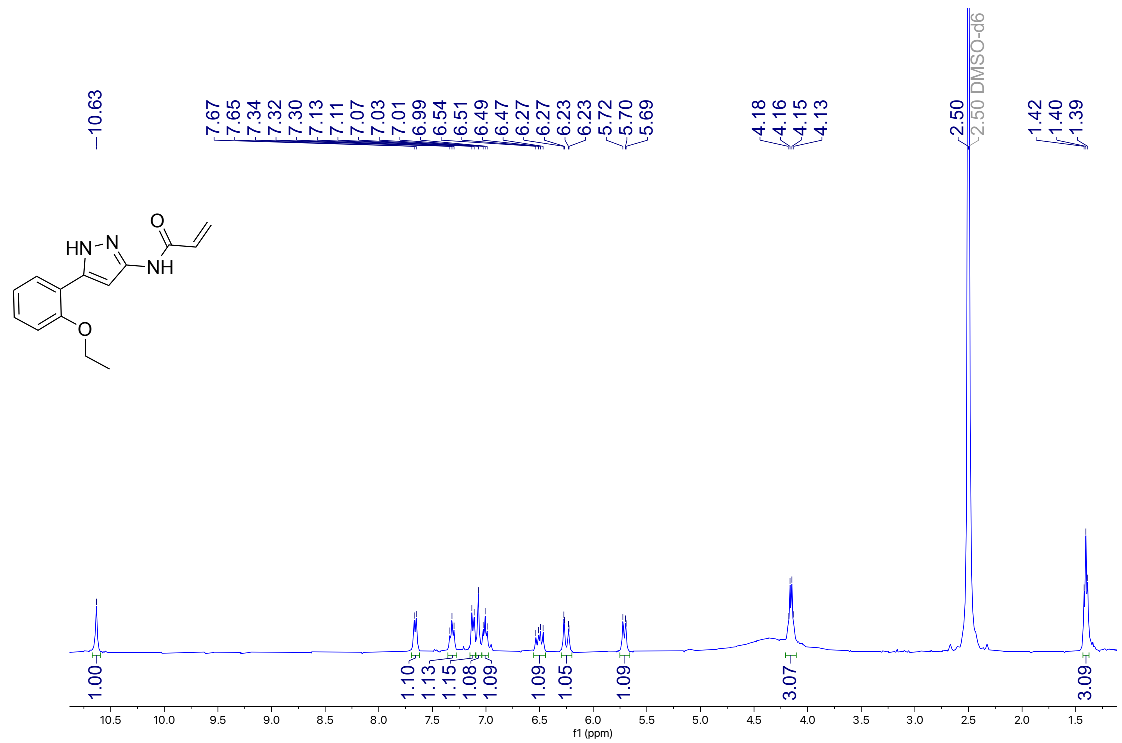


**Figure S7**: $^{13}C$ NMR (101 MHz, DMSO-$d_6$) for **2**

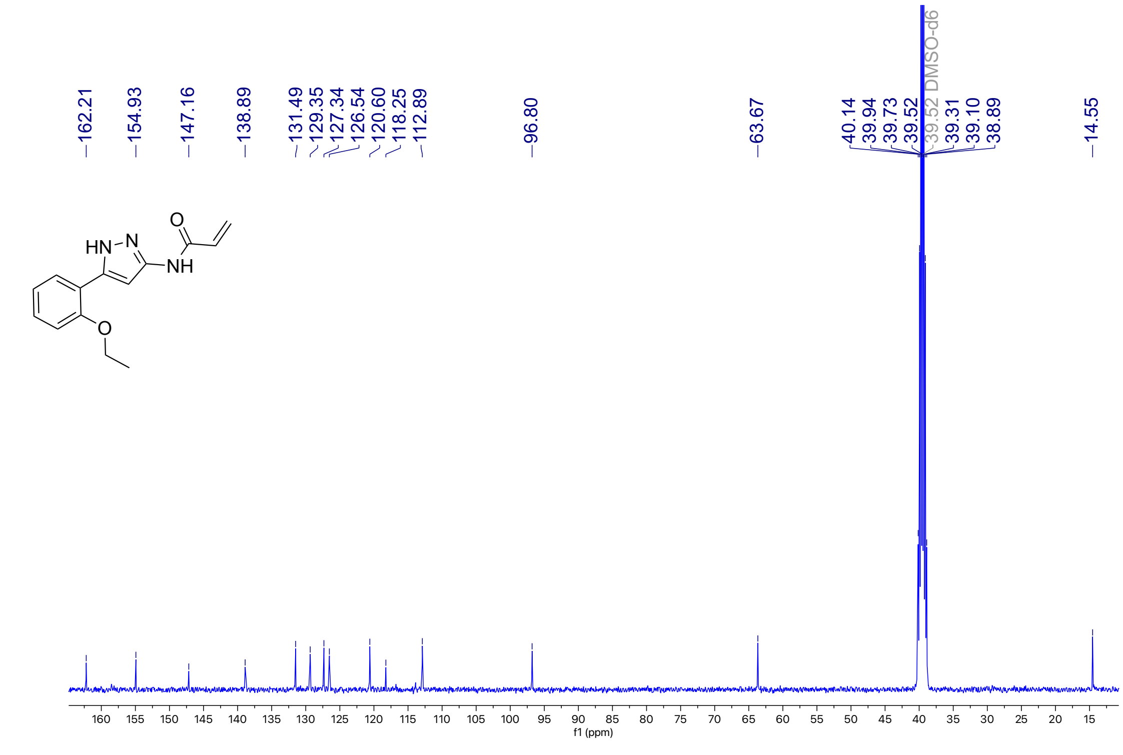

**Figure S8**: $^{1}$H NMR (400 MHz, DMSO-$d_6$) for **3**

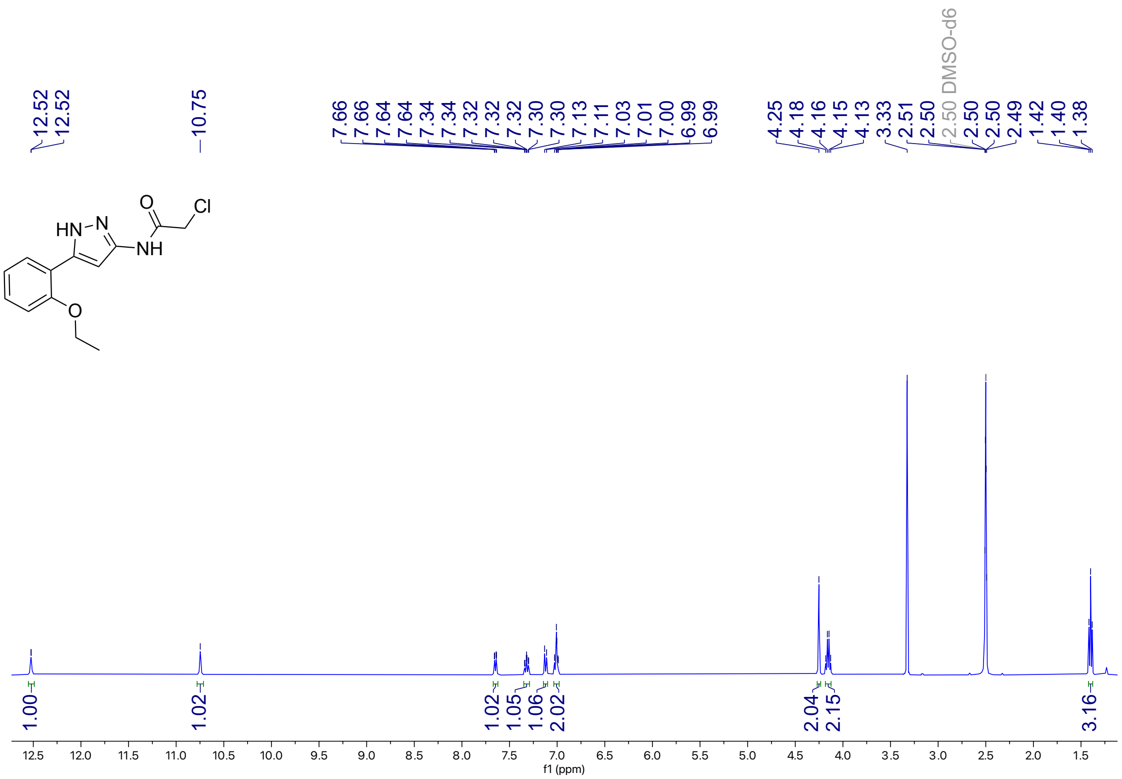

**Figure S9**: $^{13}$C NMR (101 MHz, DMSO-$d_6$) for **3**

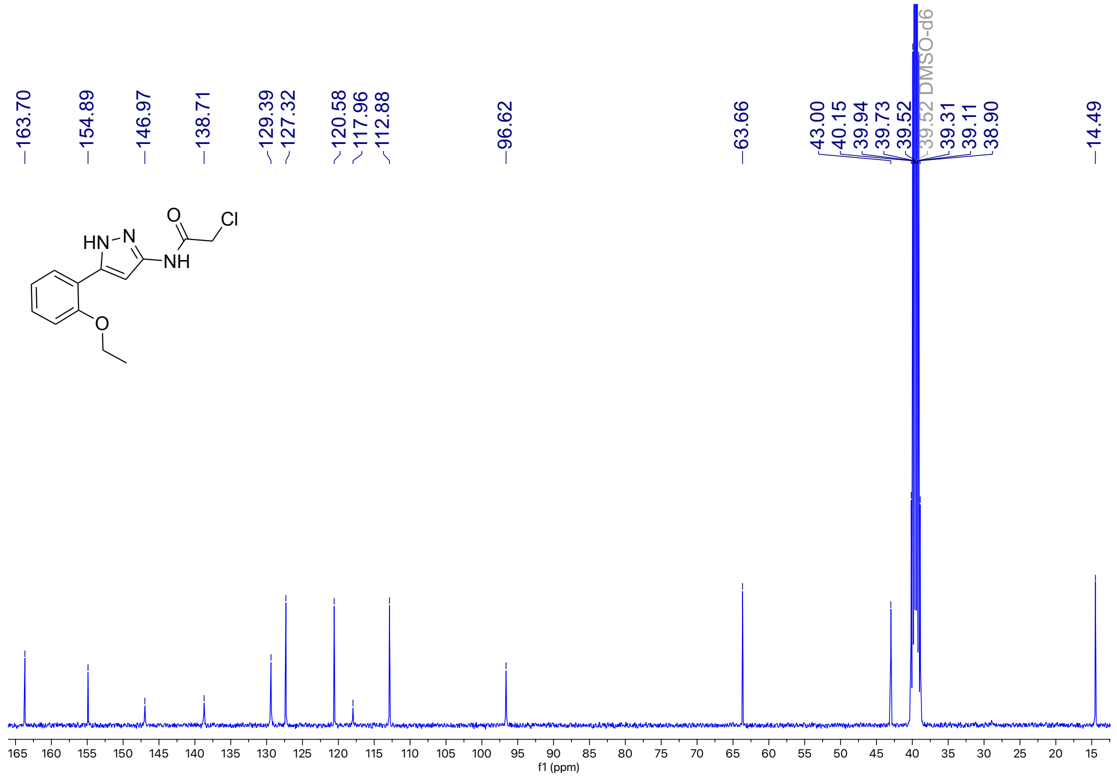

**Figure S10**: $^{1}$H NMR (500 MHz, DMSO-$d_6$) for **4**

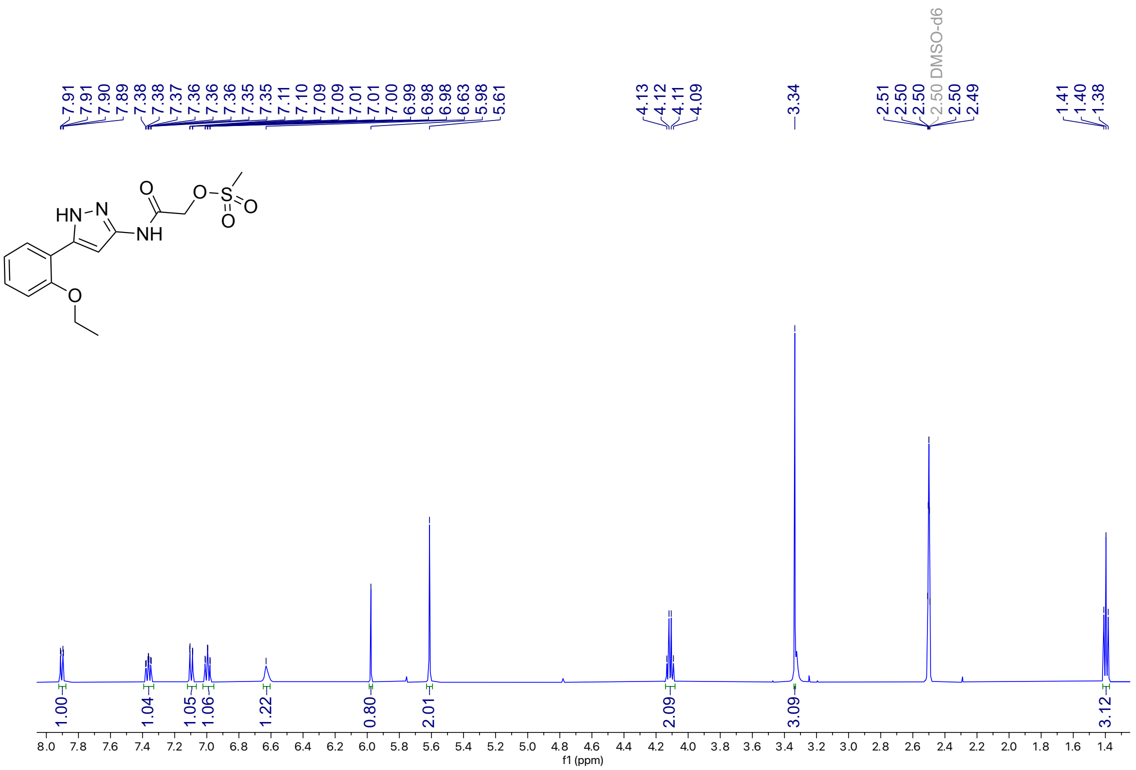

**Figure S11**: $^{13}$C NMR (126 MHz, DMSO-$d_6$) for **4**

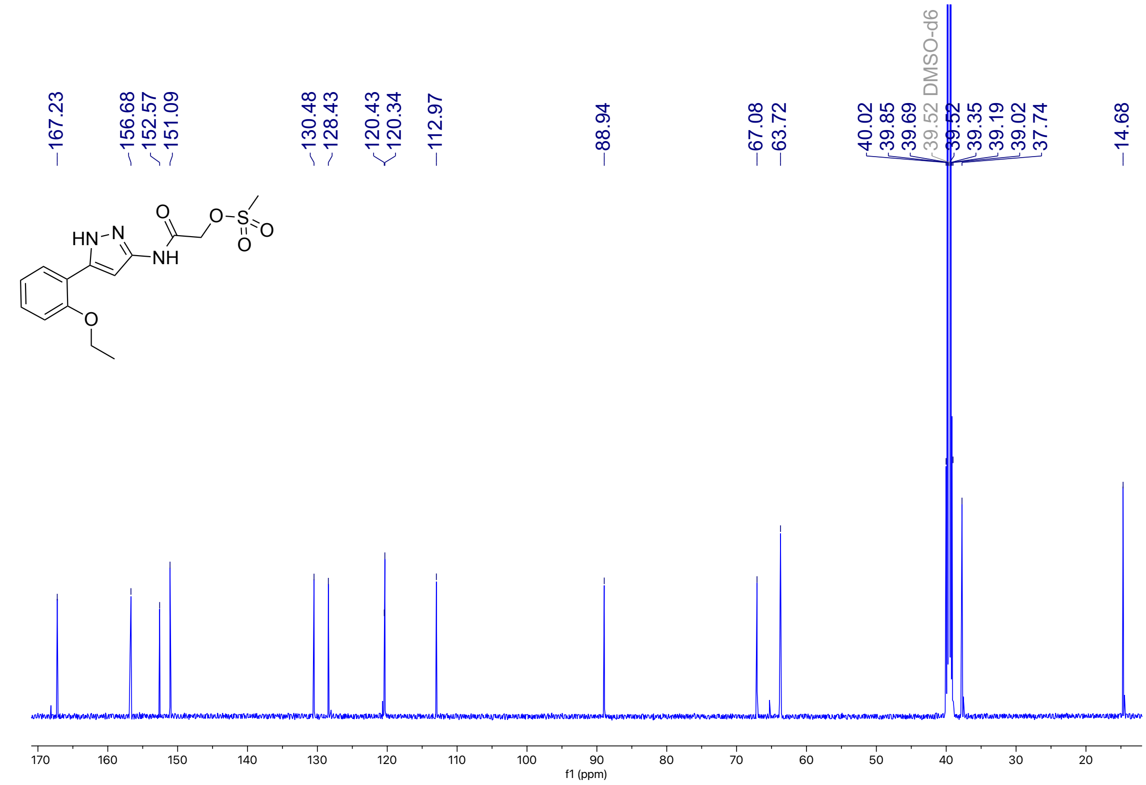

**Figure S12**: $^{1}$H NMR (401 MHz, DMSO-$d_6$) for **5**

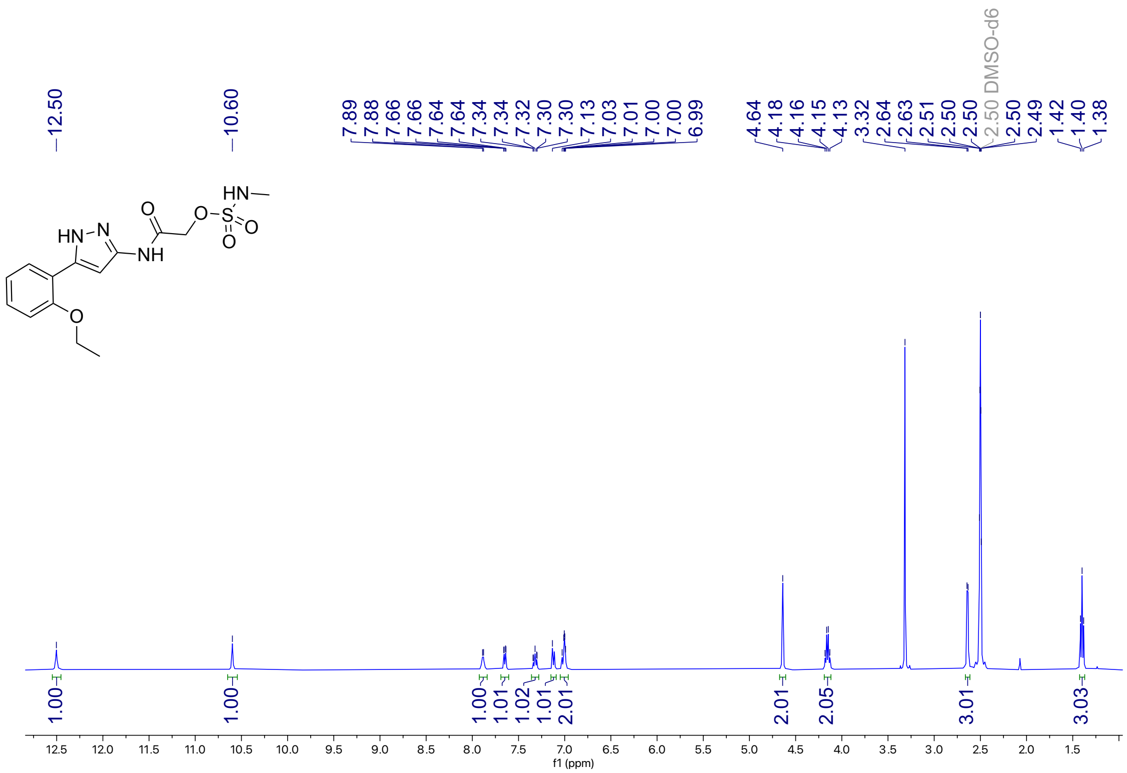

**Figure S13**: $^{13}$C NMR (101 MHz, DMSO-$d_6$) for **5**

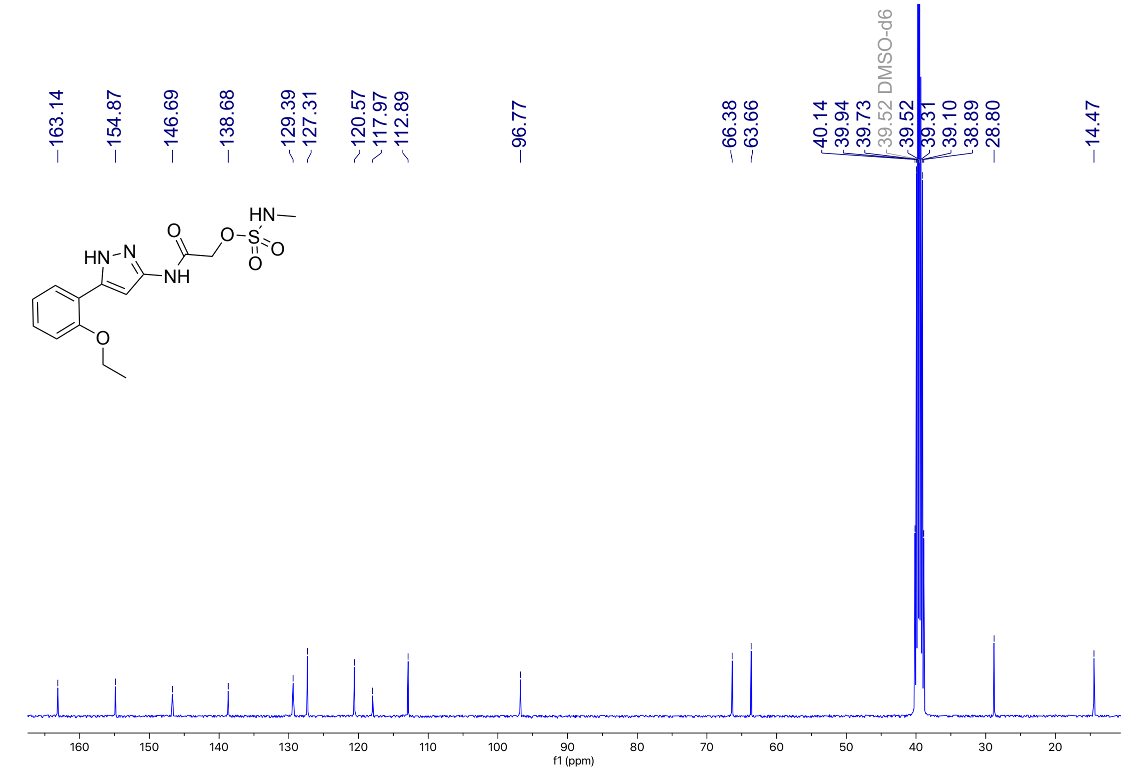

**Figure S14**: $^{1}$H NMR (400 MHz, DMSO-$d_6$) for **6a**

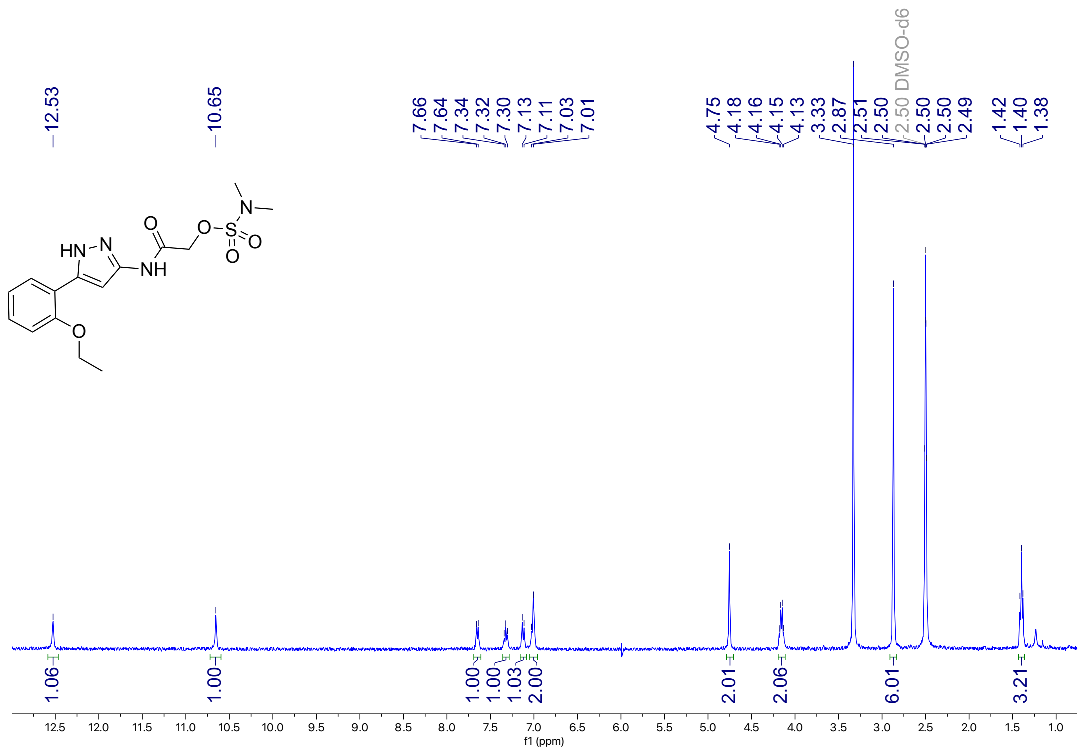


**Figure S15**: $^{13}$C NMR (101 MHz, DMSO-$d_6$) for **6a**

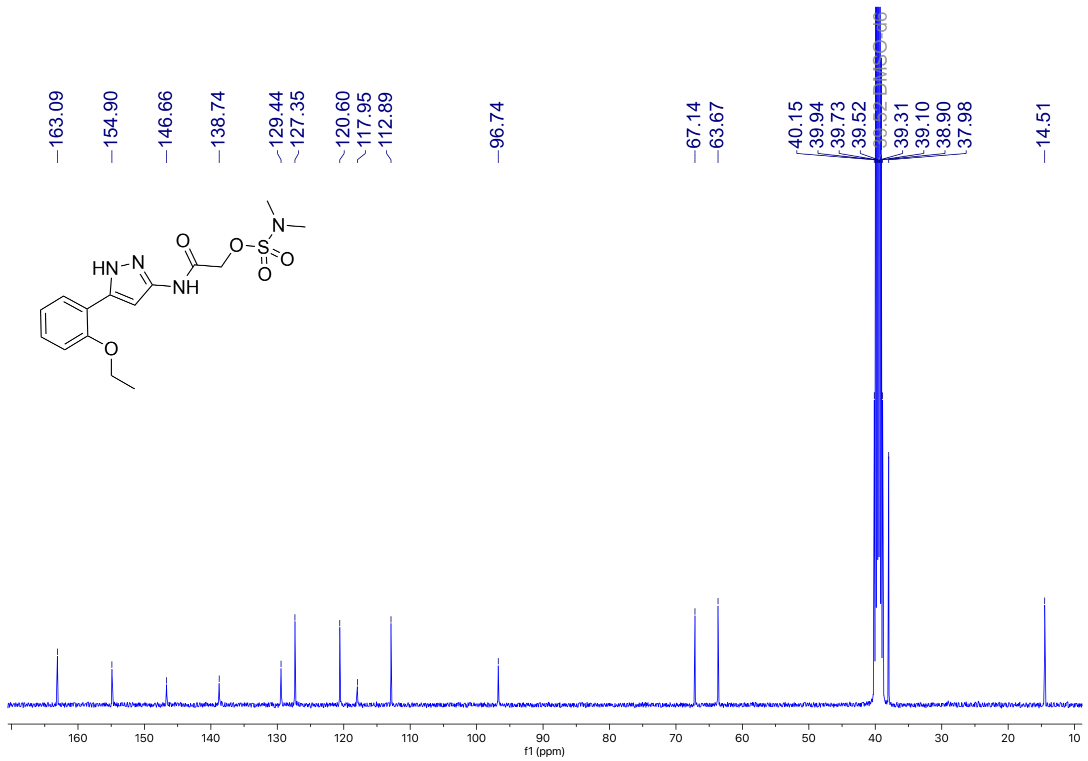

**Figure S16**: $^{1}$H NMR (400 MHz, DMSO-$d_6$) for **6b**

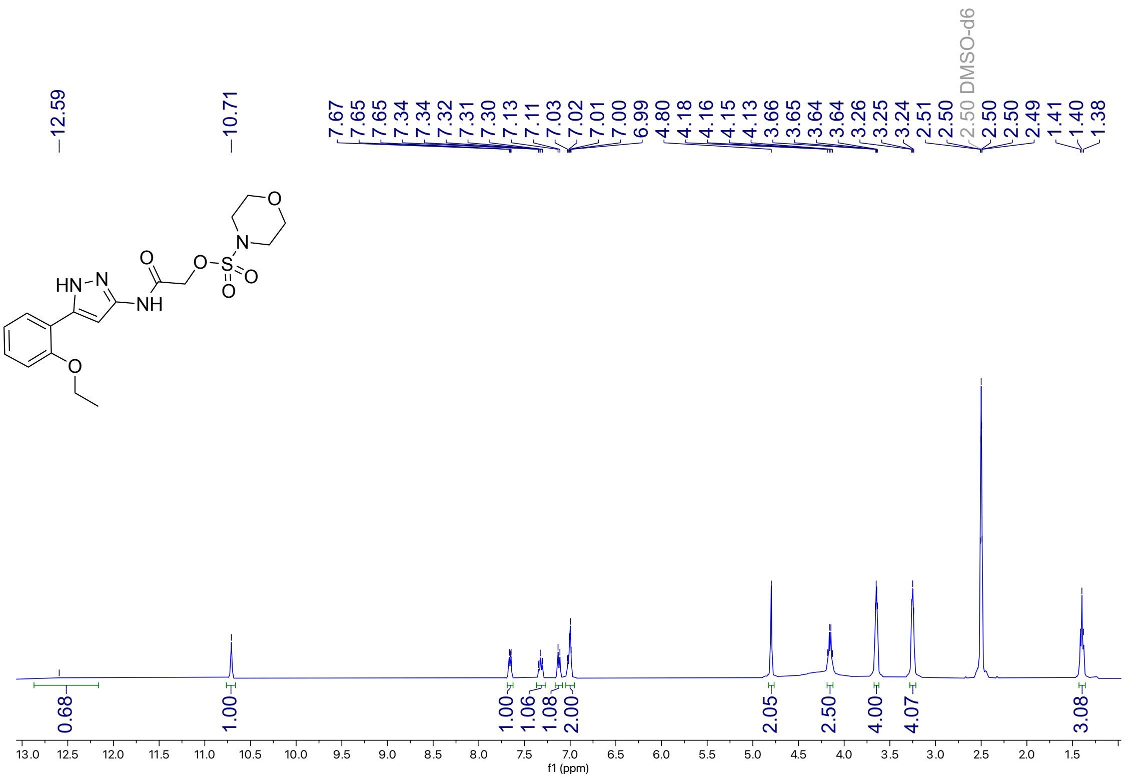

**Figure S17**: $^{13}$C NMR (214 MHz, DMSO-$d_6$) for **6b**

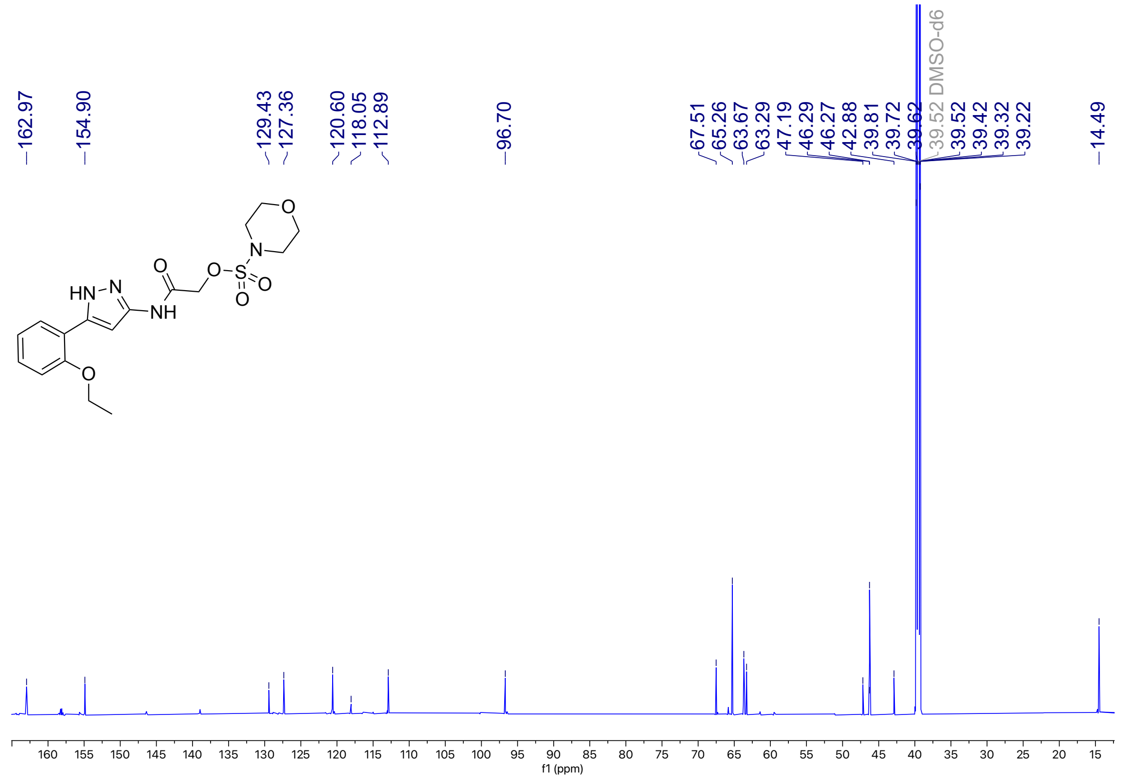

**Figure S18**: $^{1}$H NMR (401 MHz, DMSO-$d_6$) for **6c**

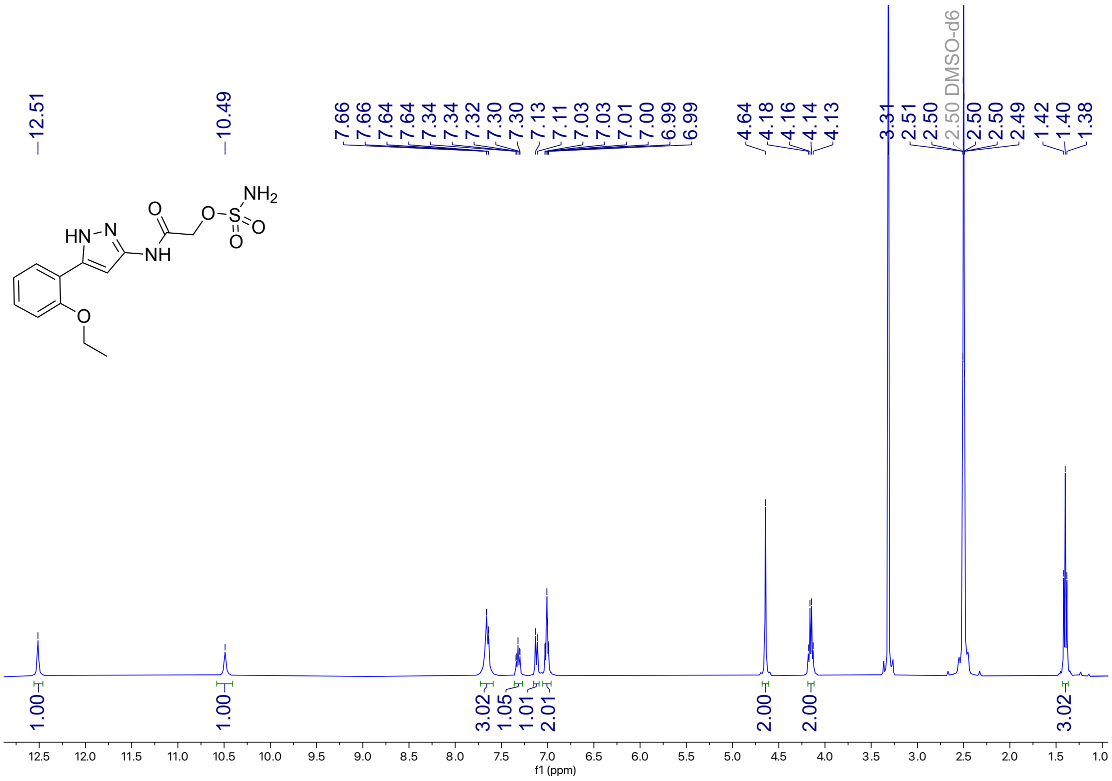


**Figure S19**: $^{13}$C NMR (101 MHz, DMSO-$d_6$) for **6c**

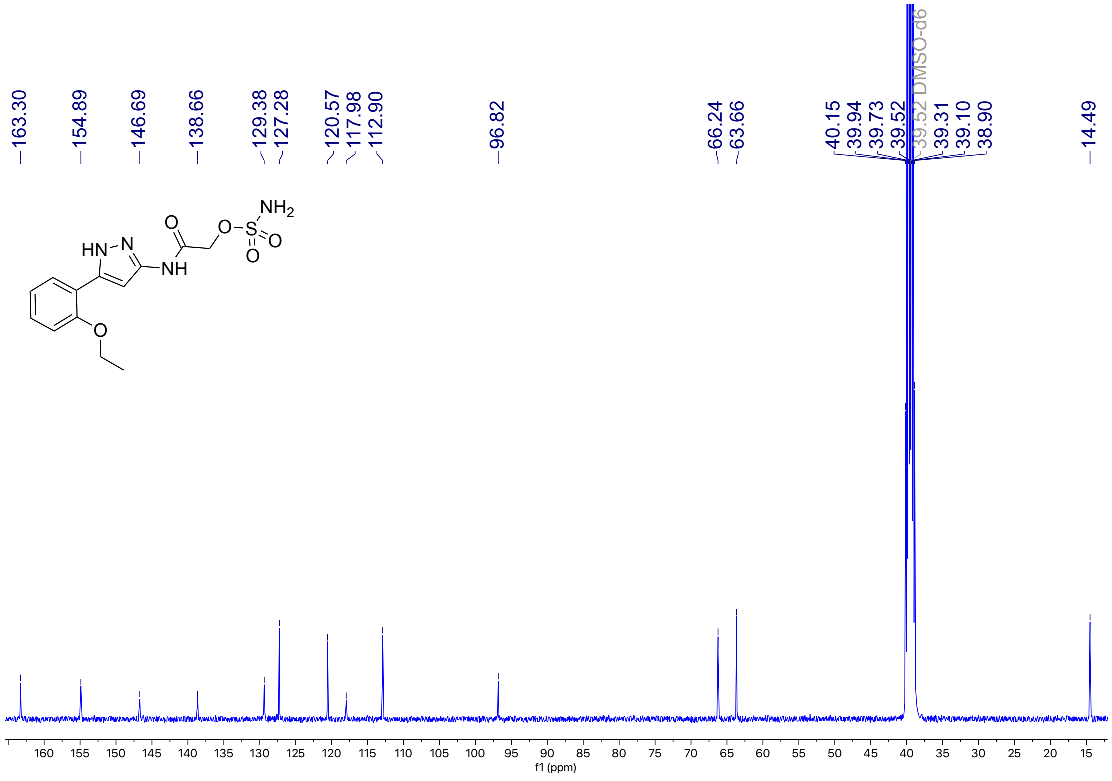

**Figure S20**: $^{1}$H NMR (400 MHz, DMSO-$d_6$) for **6d**

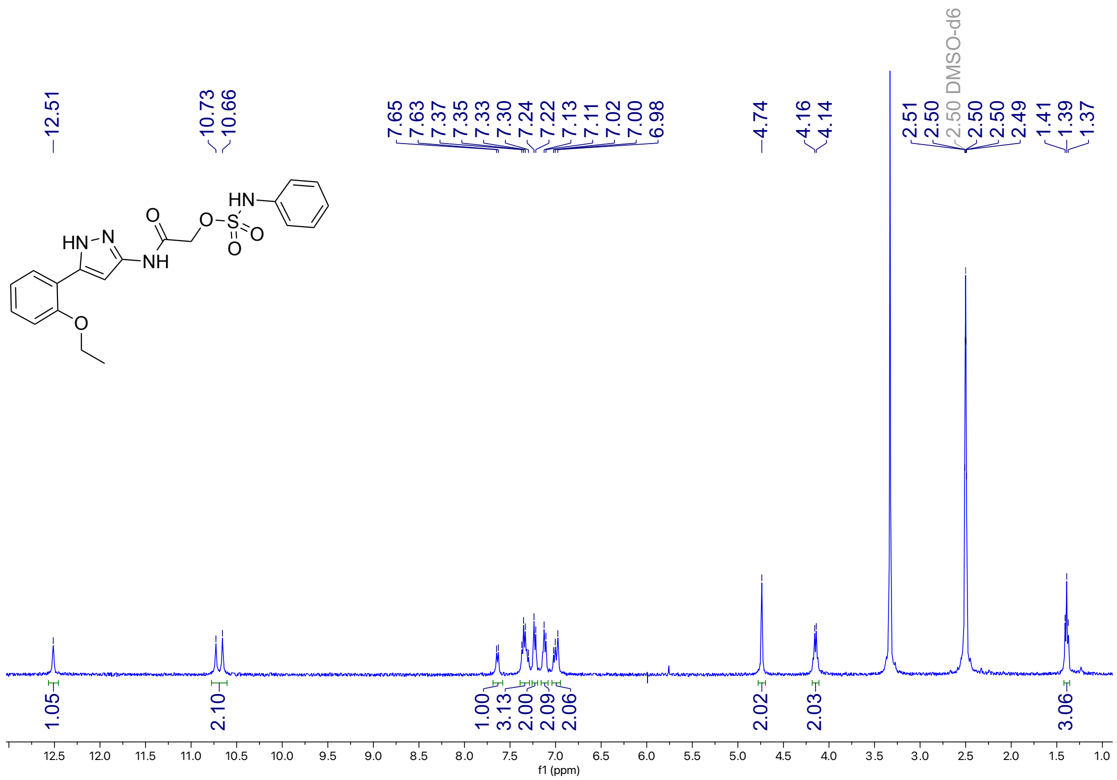


**Figure S21**: $^{13}$C NMR (101 MHz, DMSO-$d_6$) for **6d**

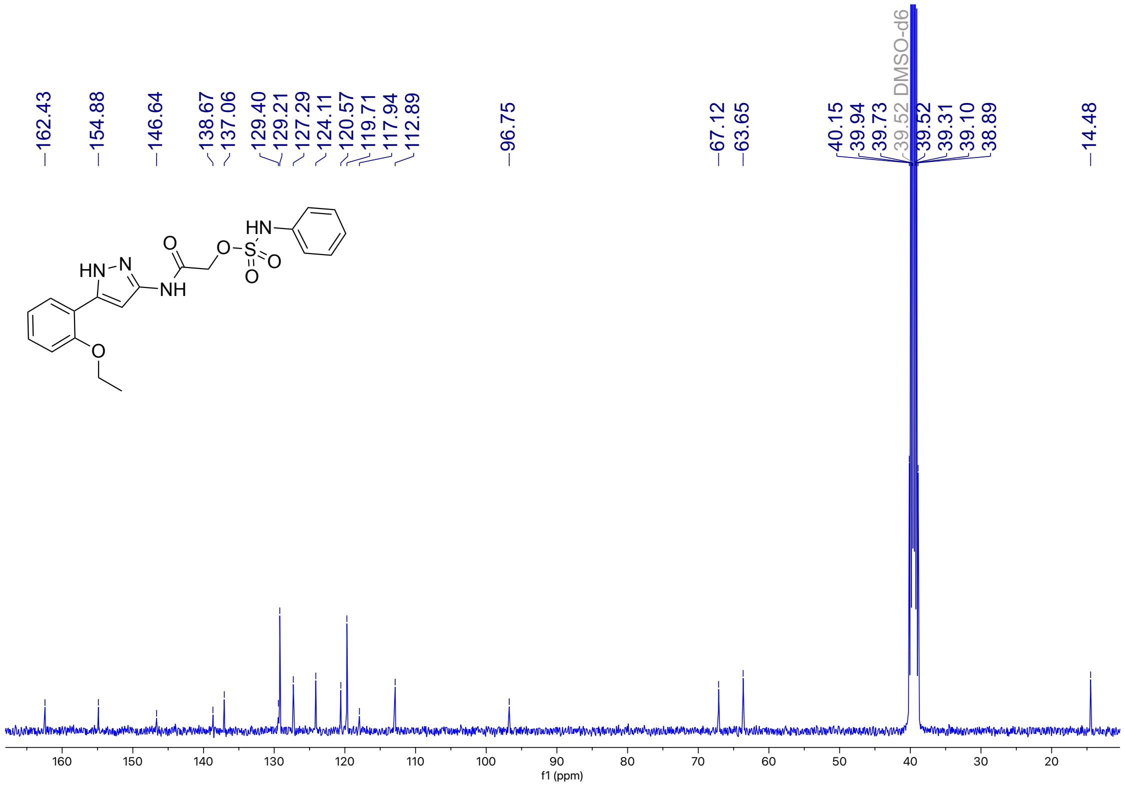

**Figure S22**: $^{1}$H NMR (400 MHz, DMSO-$d_6$) for **6e**

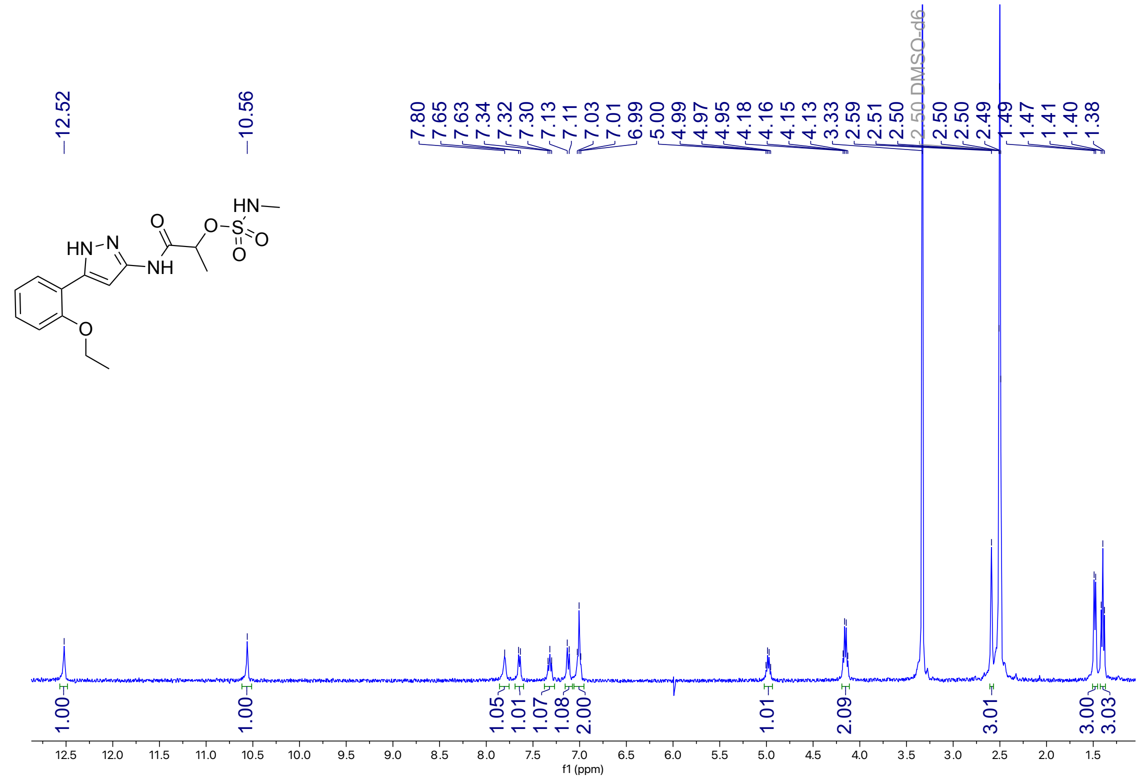

**Figure S23**: $^{13}$C NMR (101 MHz, DMSO-$d_6$) for **6e**

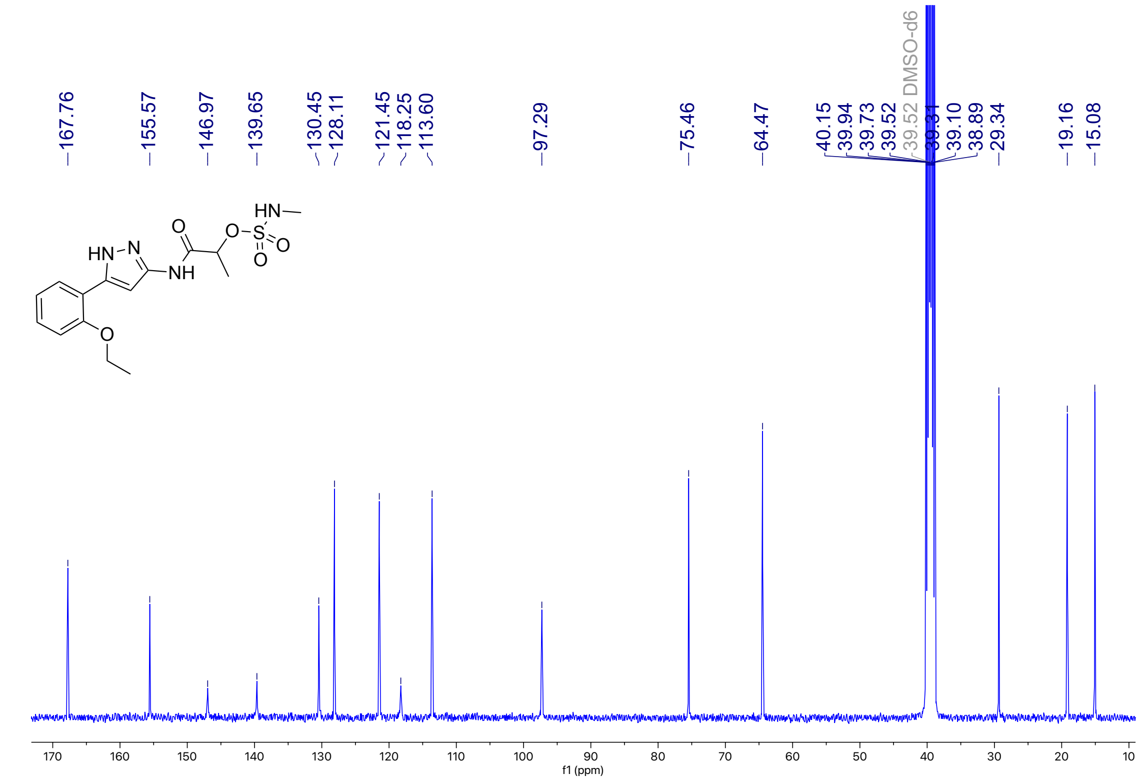

**Figure S24**: $^{1}$H NMR (401 MHz, DMSO-$d_6$) for **6f**

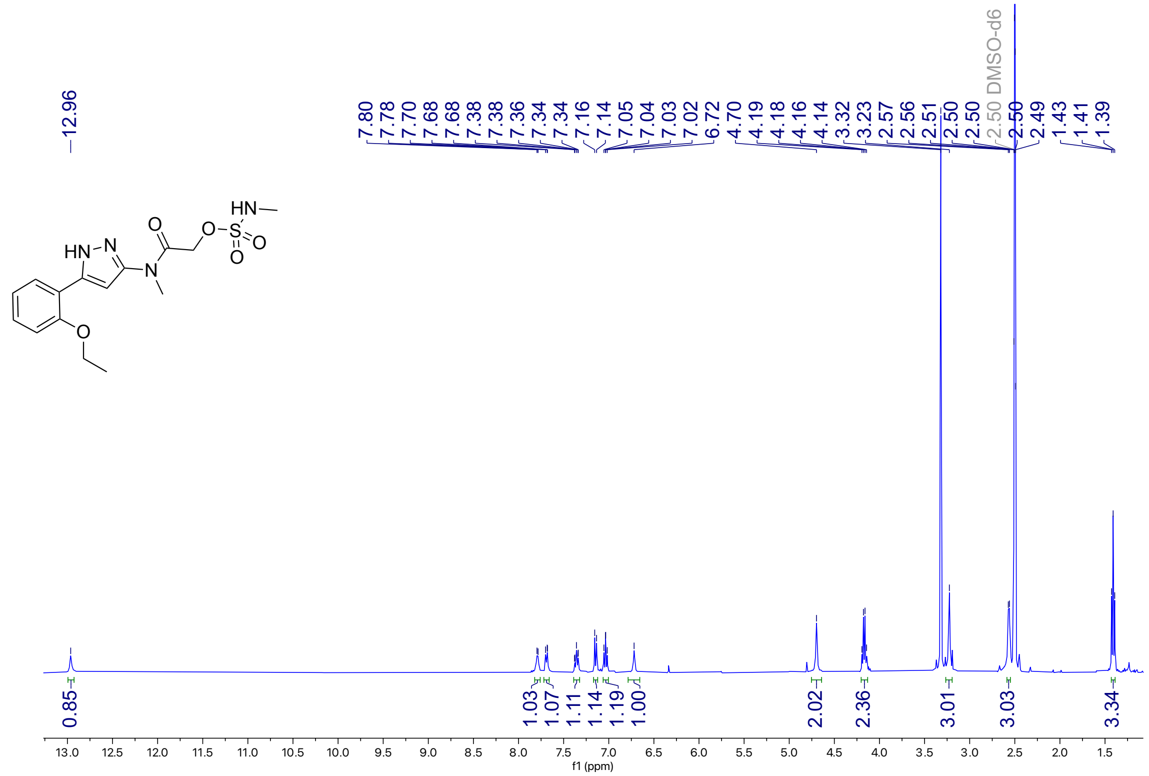


**Figure S25**: $^{13}$C NMR (101 MHz, DMSO-$d_6$) for **6f**

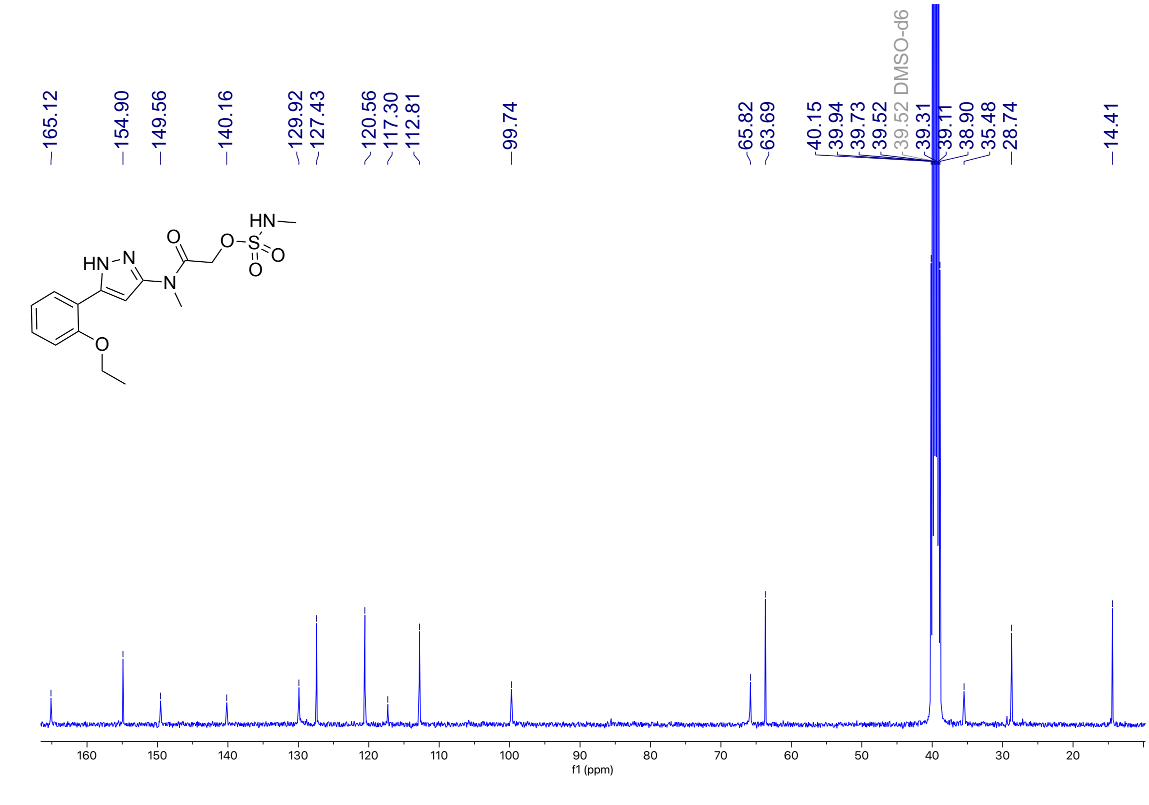

**Figure S26**: $^{1}$H NMR (400 MHz, DMSO-$d_6$) for **7a**

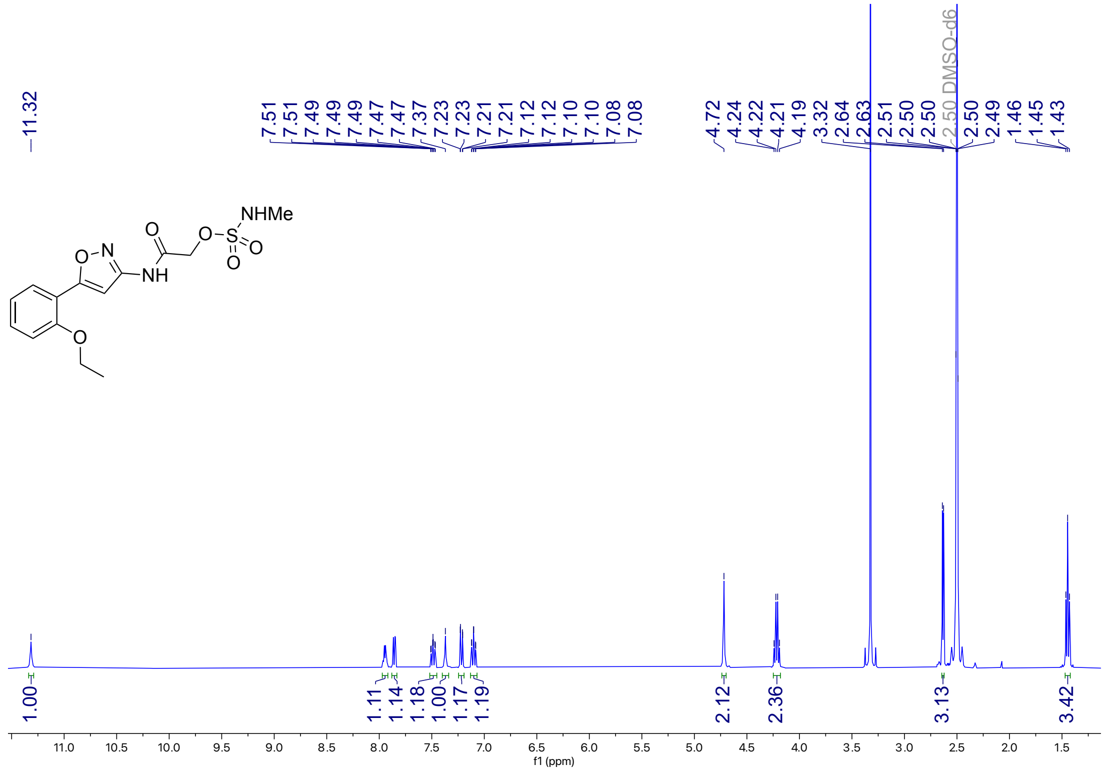

**Figure S27**: $^{13}$C NMR (101 MHz, DMSO-$d_6$) for **7a**

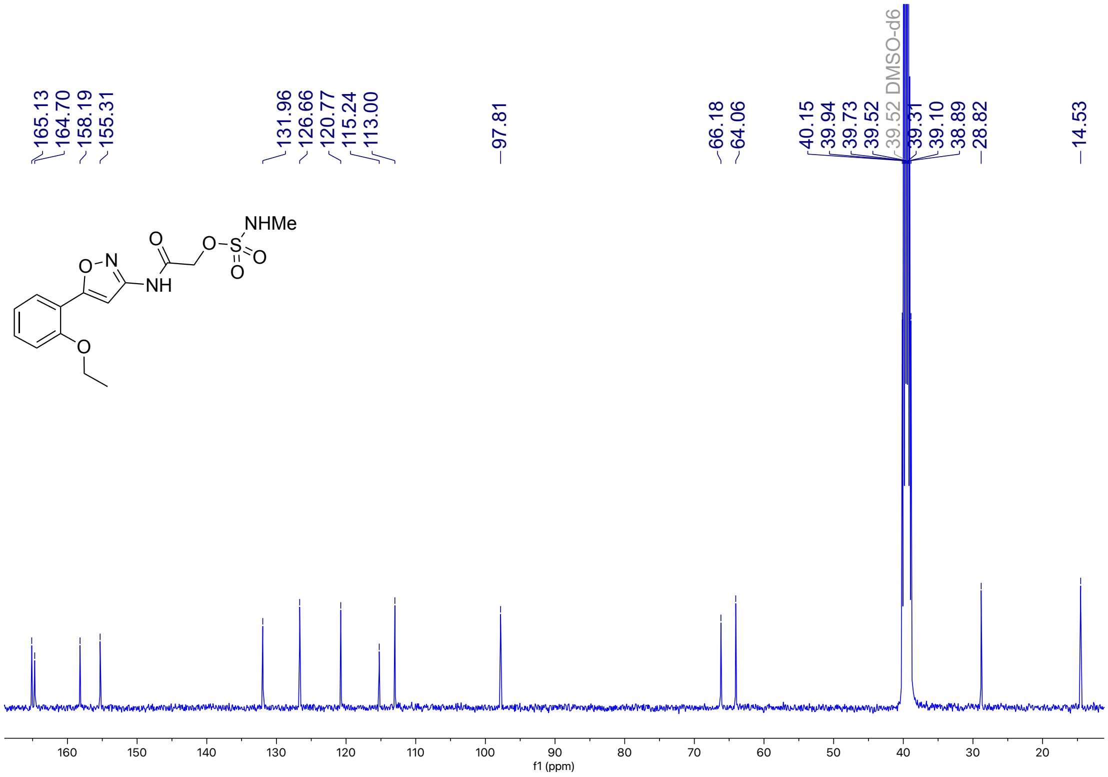

**Figure S28**: $^{1}$H NMR (401 MHz, DMSO-$d_6$) for **7b**

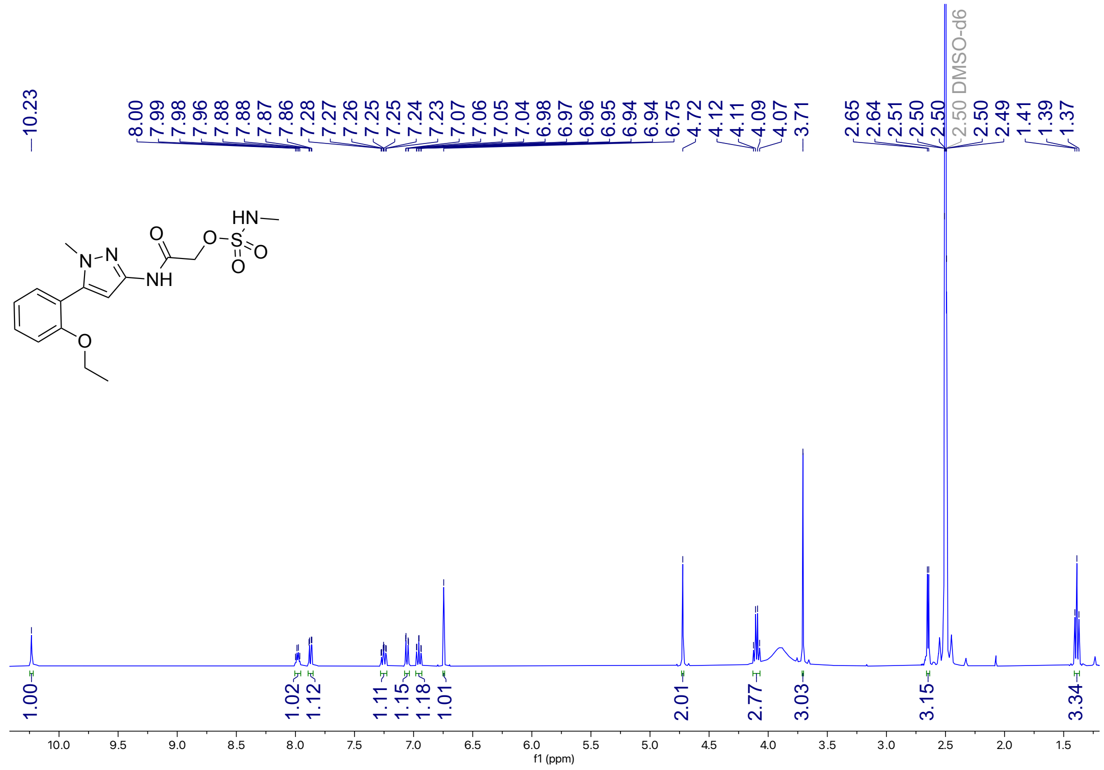


**Figure S29**: $^{13}$C NMR (101 MHz, DMSO-$d_6$) for **7b**

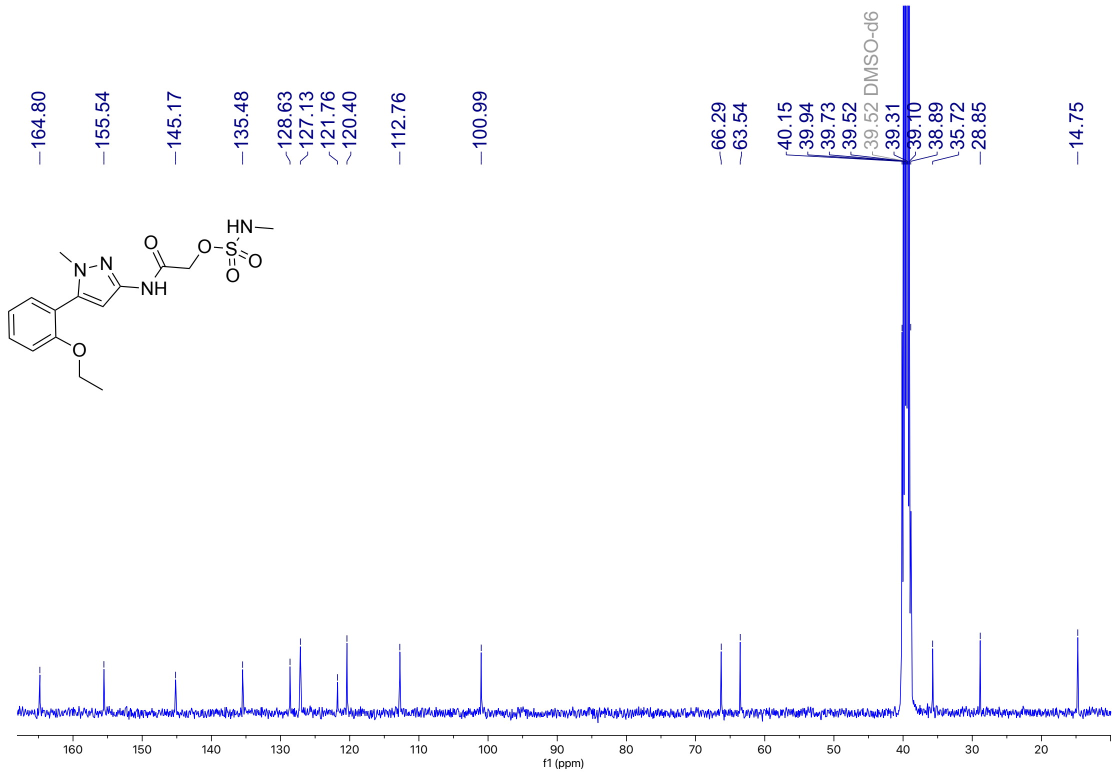

**Figure S30**: $^{1}$H NMR (401 MHz, DMSO-$d_6$) for **7c**

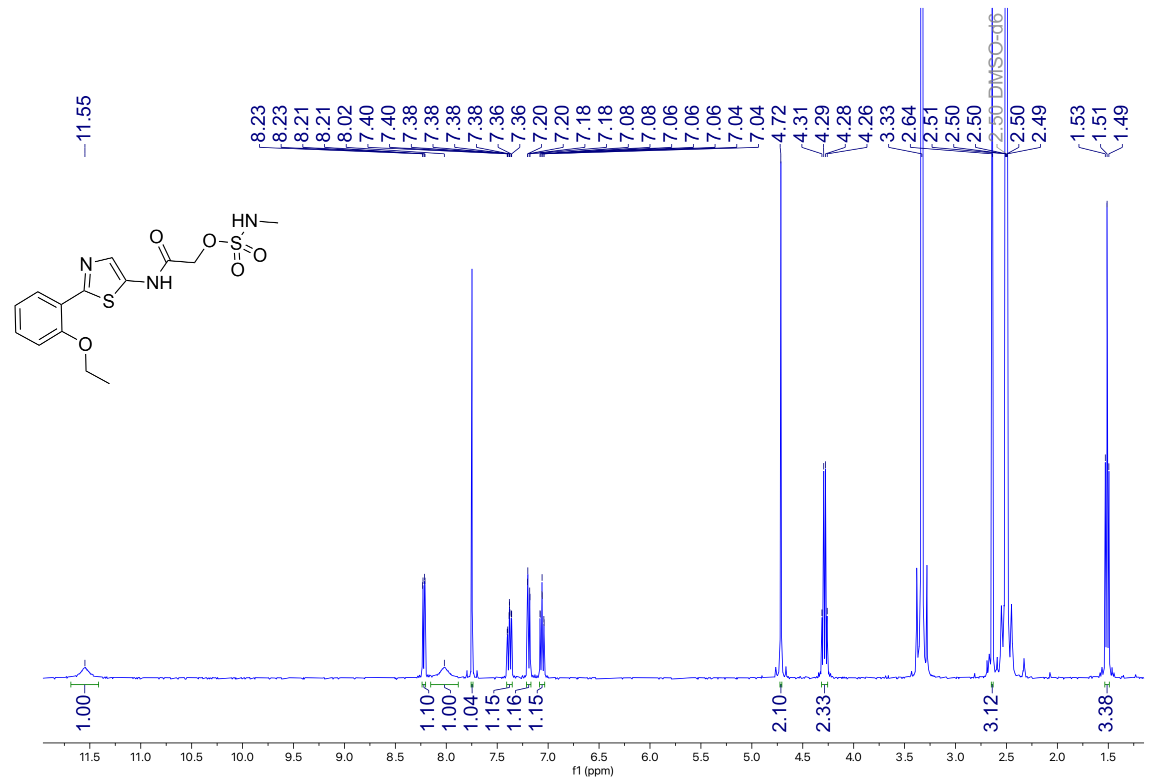

**Figure S31**: $^{13}$C NMR (101 MHz, DMSO-$d_6$) for **7c**

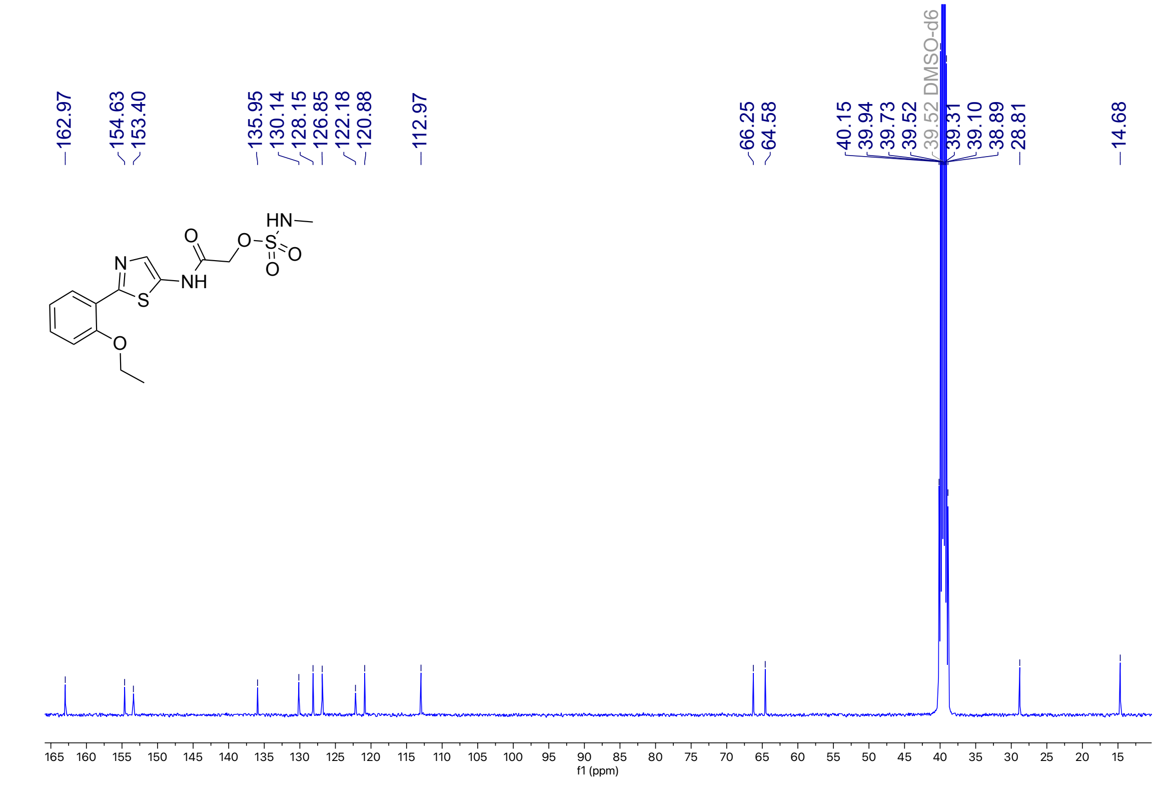

**Figure S32**: $^{1}$H NMR (401 MHz, DMSO-$d_6$) for **7d**

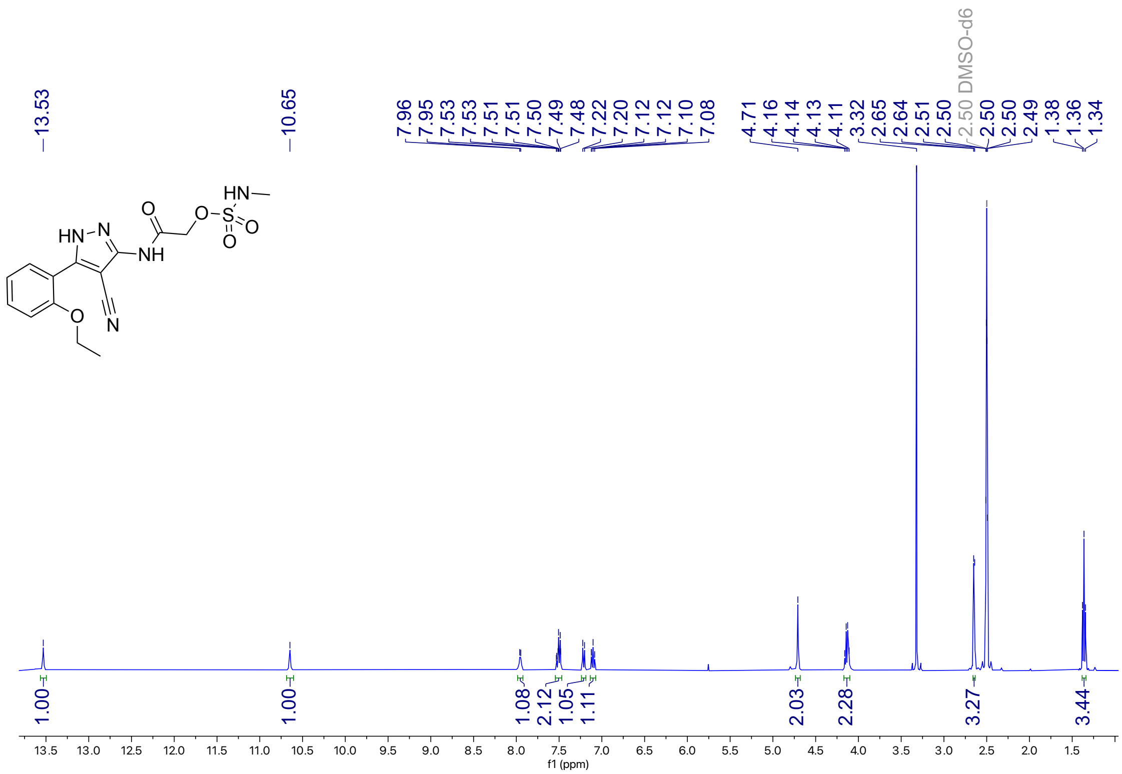


**Figure S33**: $^{13}$C NMR (101 MHz, DMSO-$d_6$) for **7d**

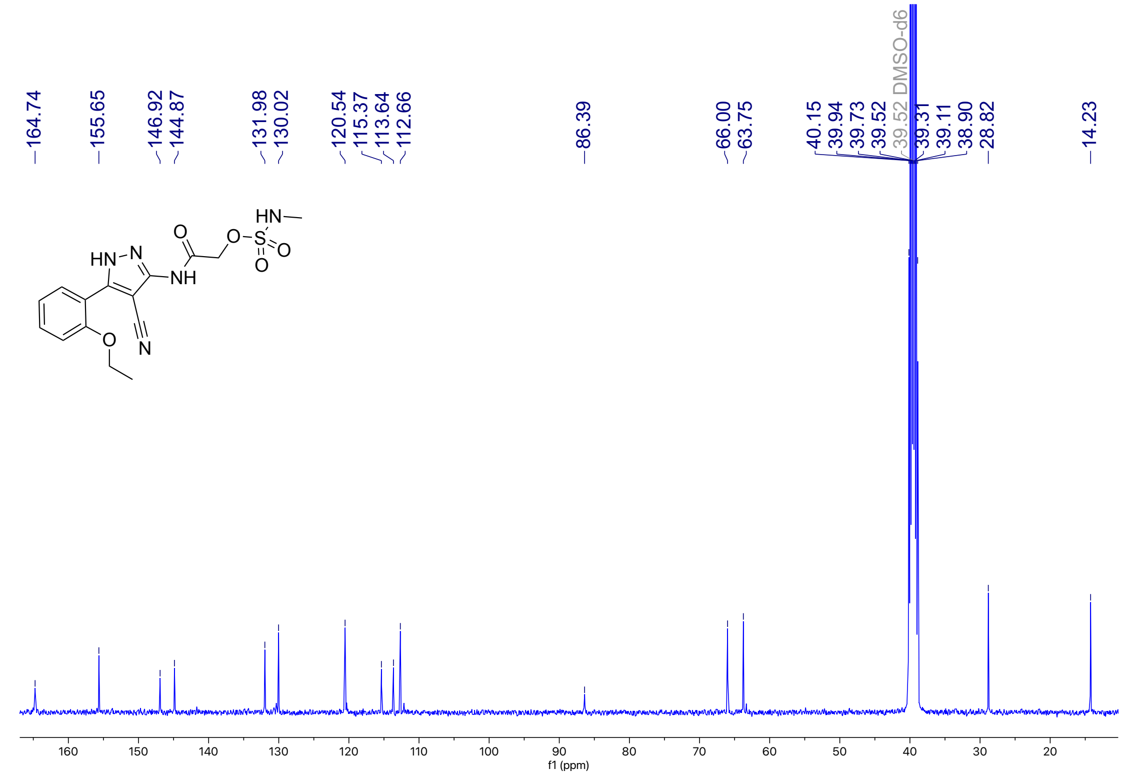

**Figure S34**: $^{1}$H NMR (401 MHz, DMSO-$d_6$) for **8a**

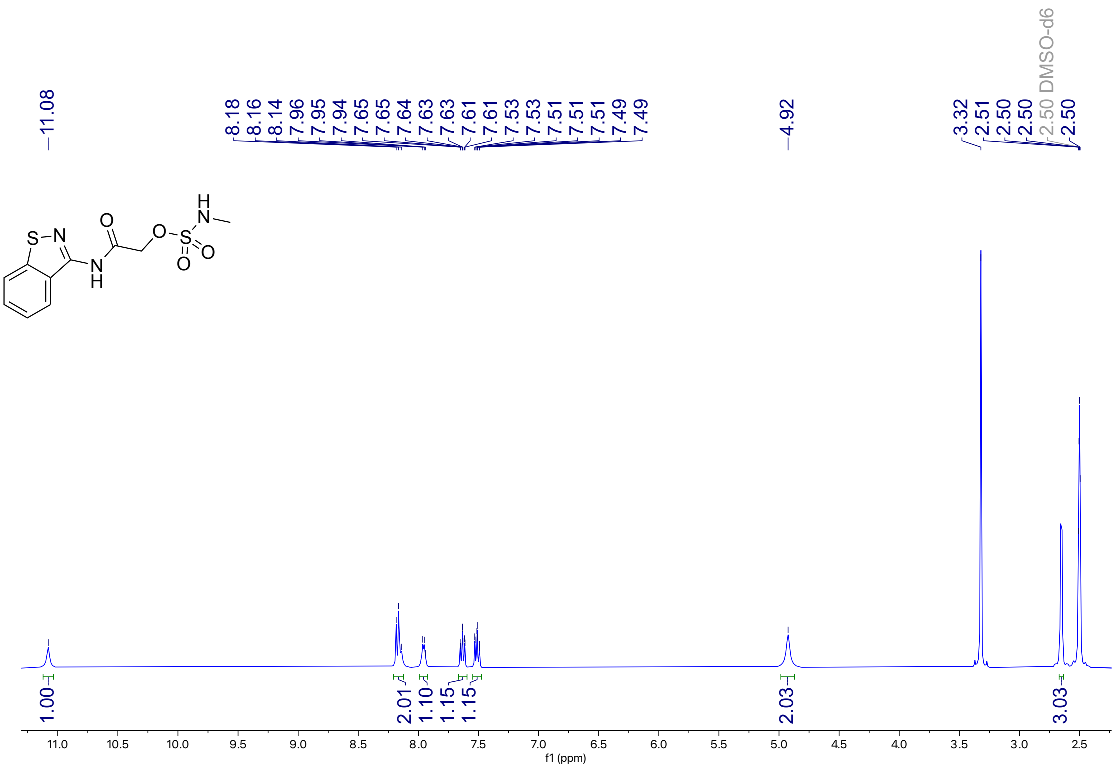

**Figure S35**: $^{13}$C NMR (101 MHz, DMSO-$d_6$) for **8a**

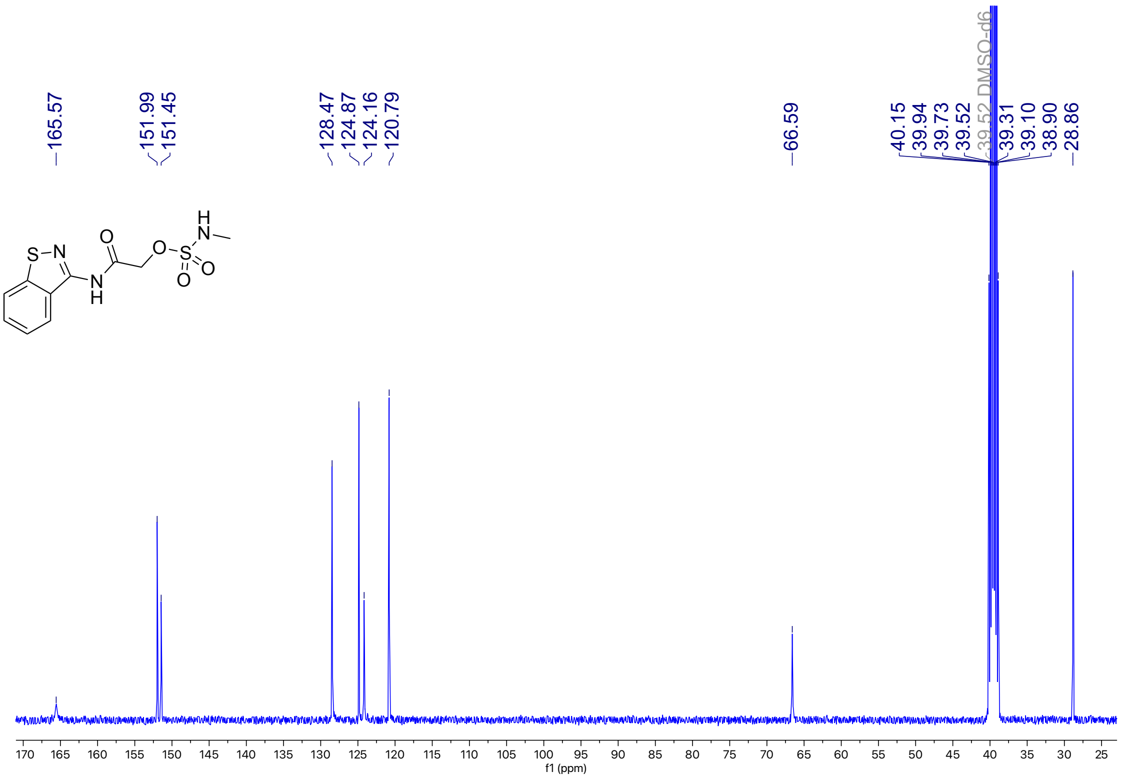

**Figure S36**: $^{1}$H NMR (401 MHz, DMSO-$d_6$) for **8b**

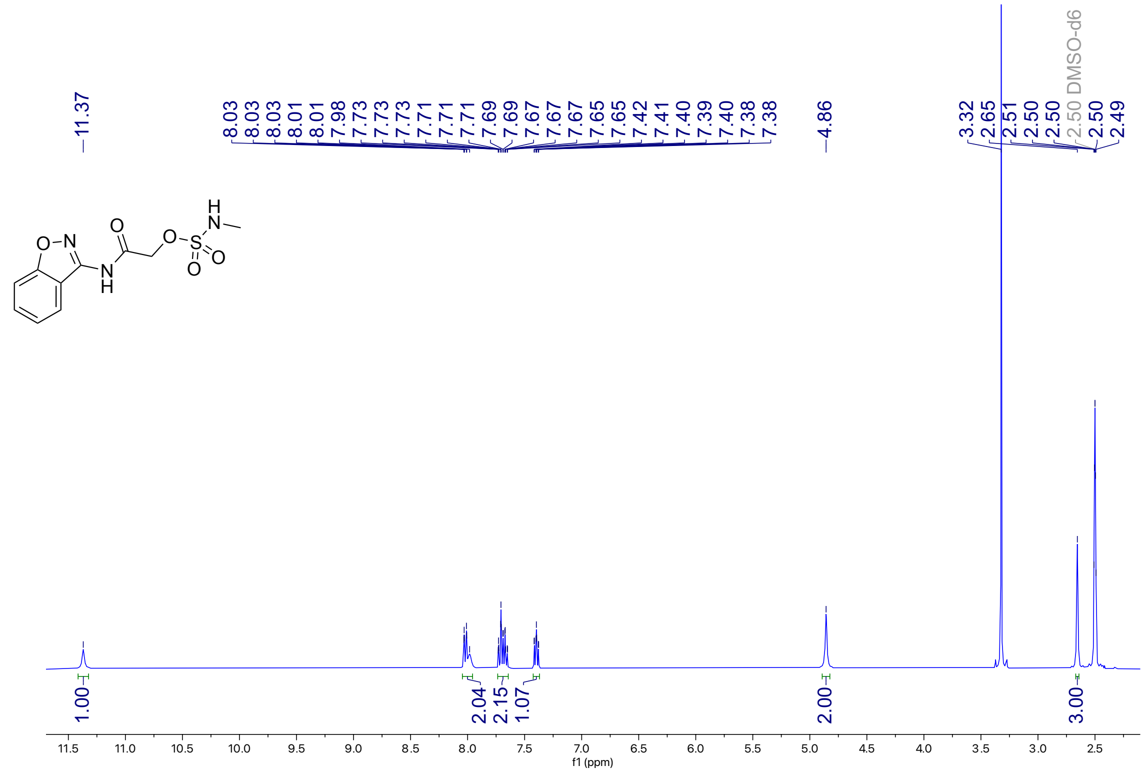

**Figure S37**: $^{13}$C NMR (101 MHz, DMSO-$d_6$) for **8b**

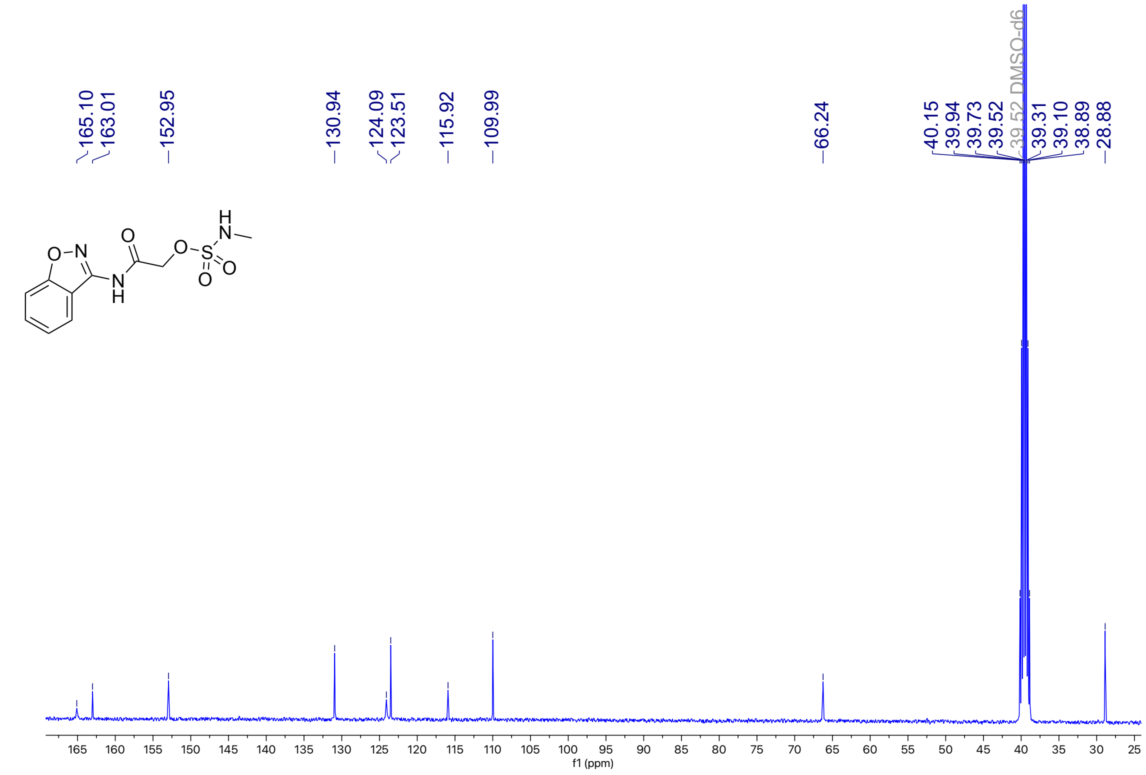

**Figure S38**: $^1$H NMR (401 MHz, DMSO-$d_6$) for **8c**

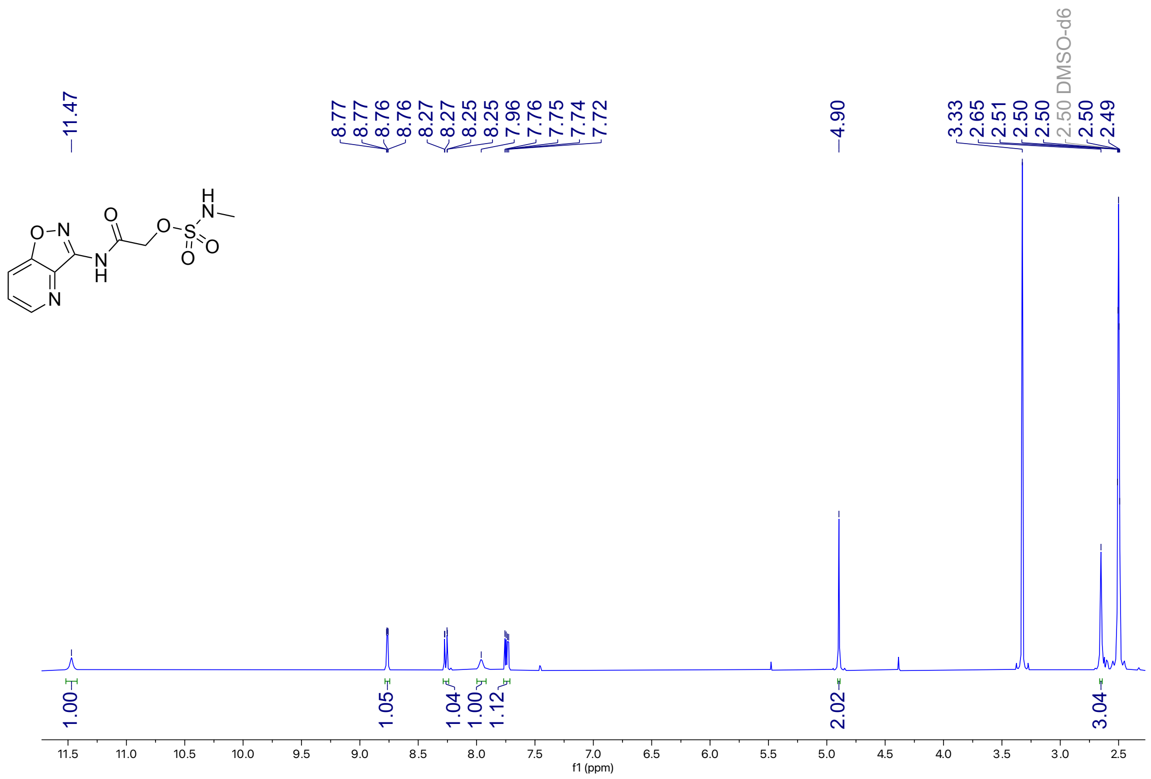

**Figure S39**: $^{13}$C NMR (101 MHz, DMSO-$d_6$) for **8c**

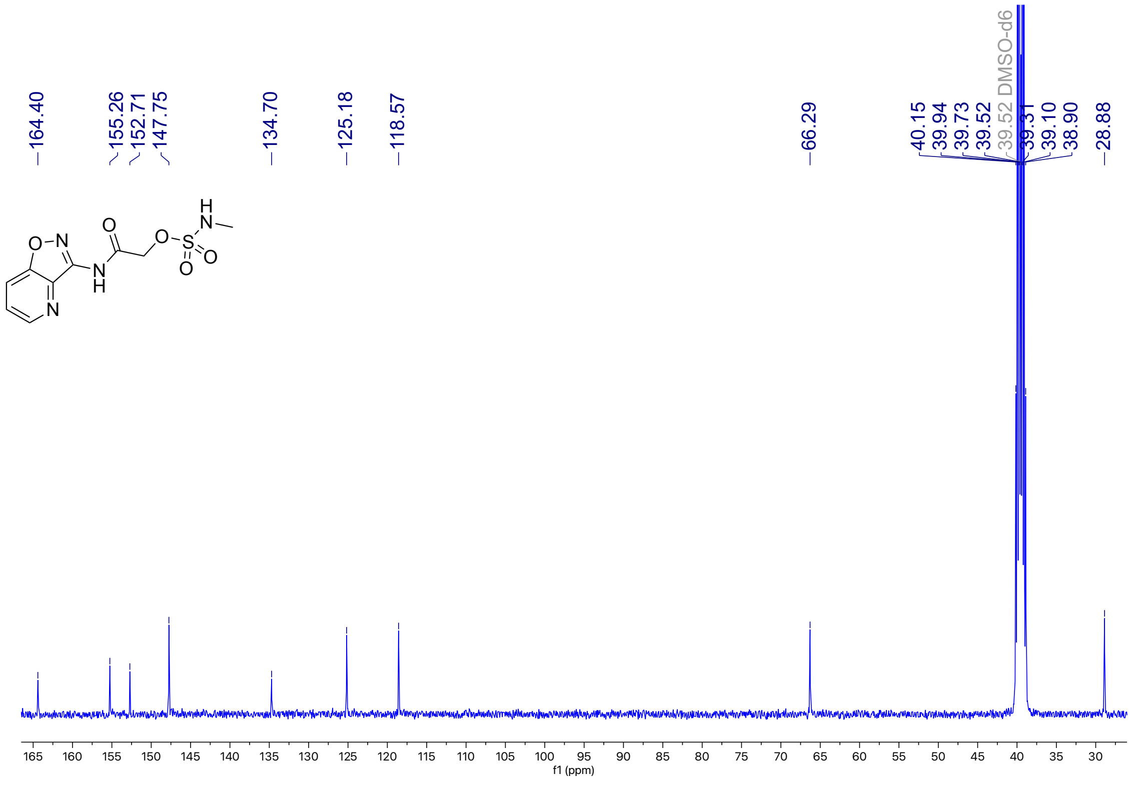

**Figure S40**: $^{1}$H NMR (401 MHz, DMSO-$d_6$) for **9a**

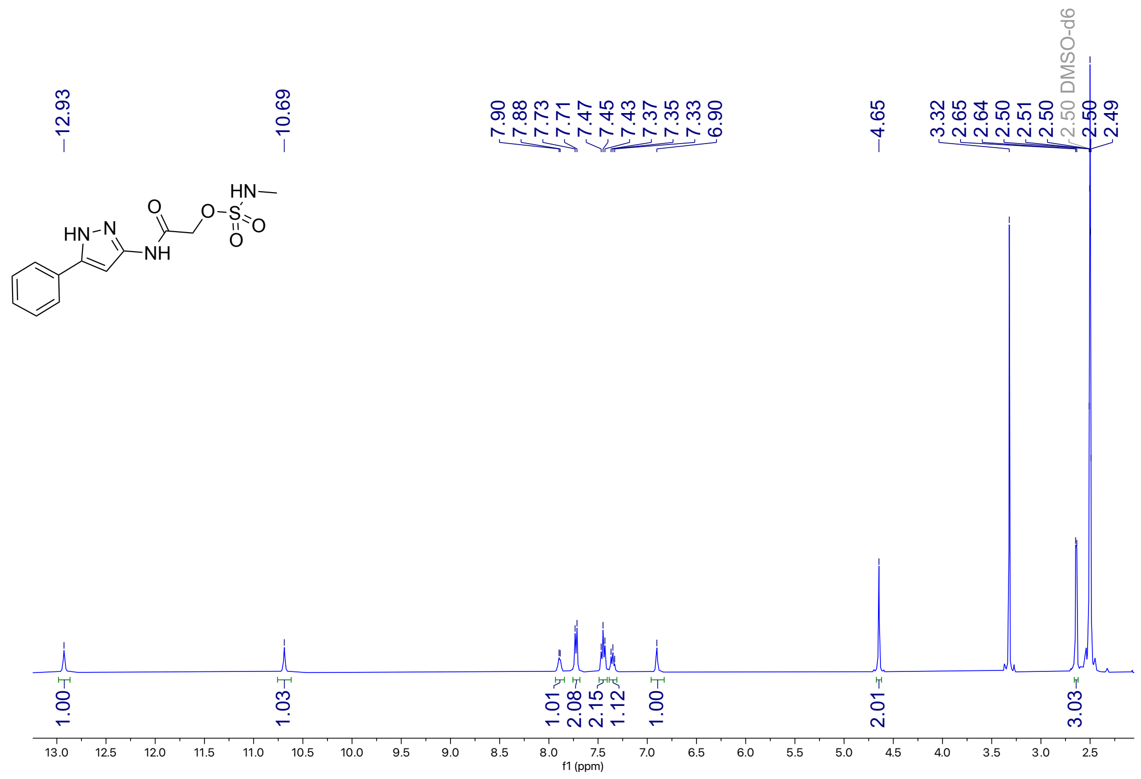

**Figure S41**: $^{13}$C NMR (101 MHz, DMSO-$d_6$) for **9a**

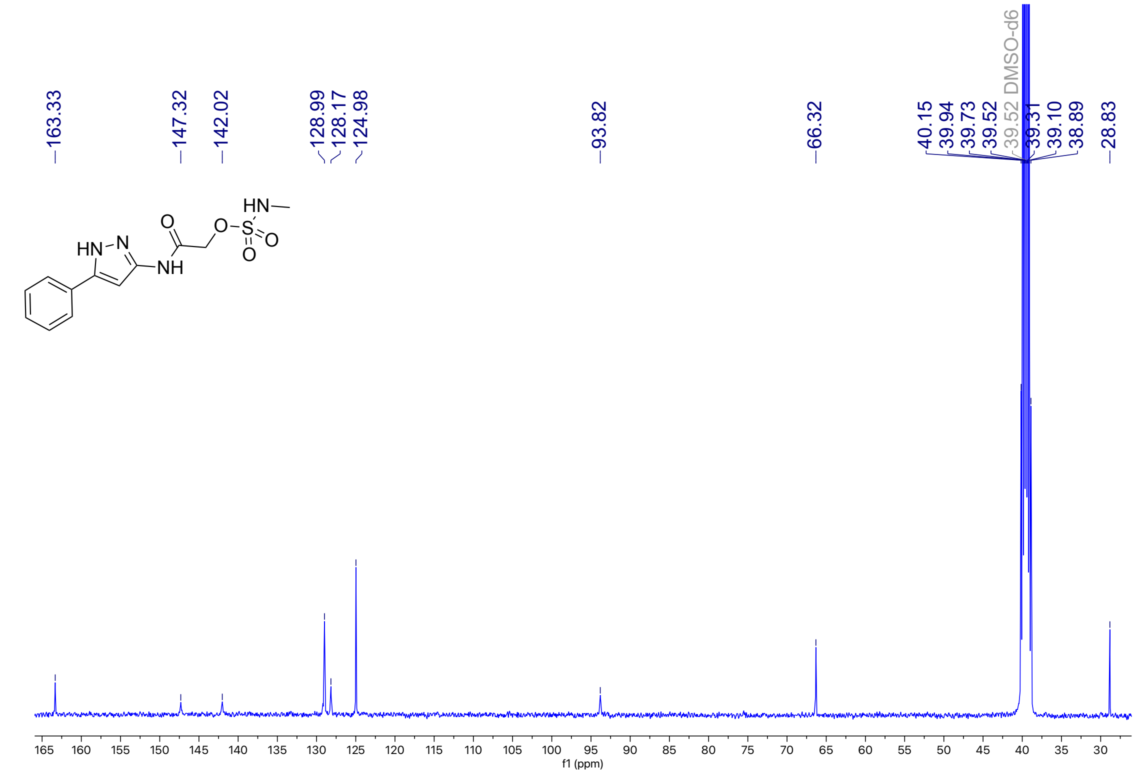

**Figure S42**: $^{1}$H NMR (400 MHz, DMSO-$d_6$) for **9b**

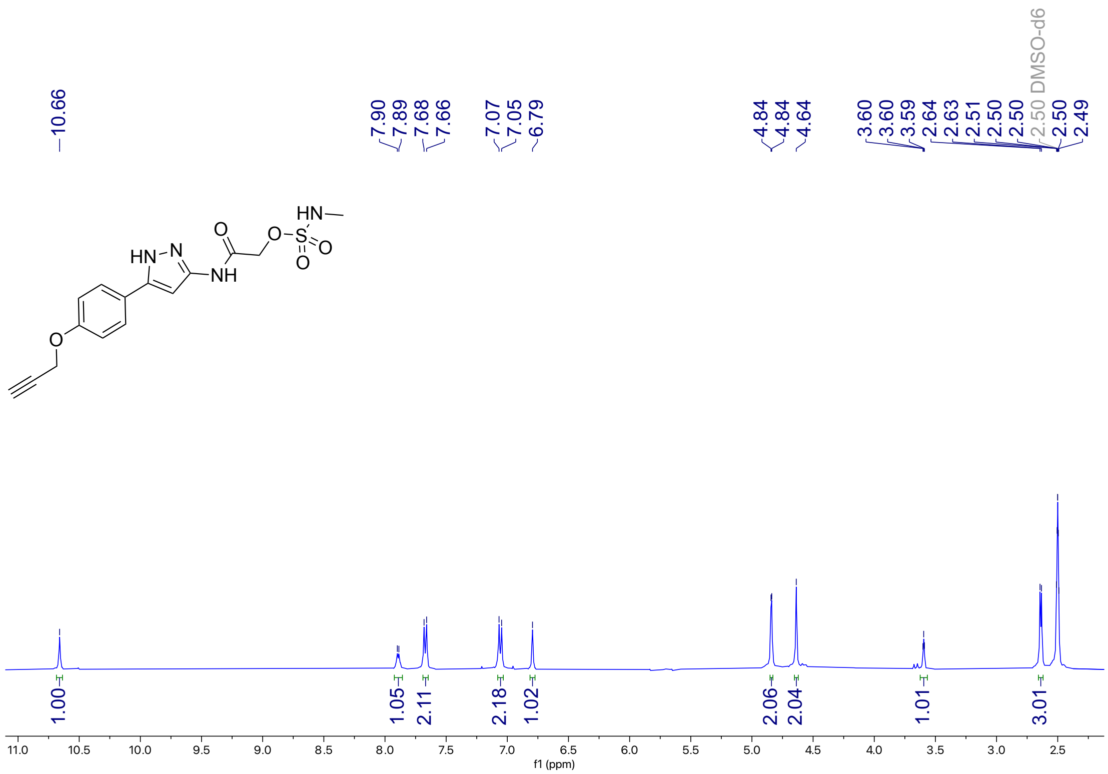

**Figure S43**: $^{13}$C NMR (214 MHz, DMSO-$d_6$) for **9b**

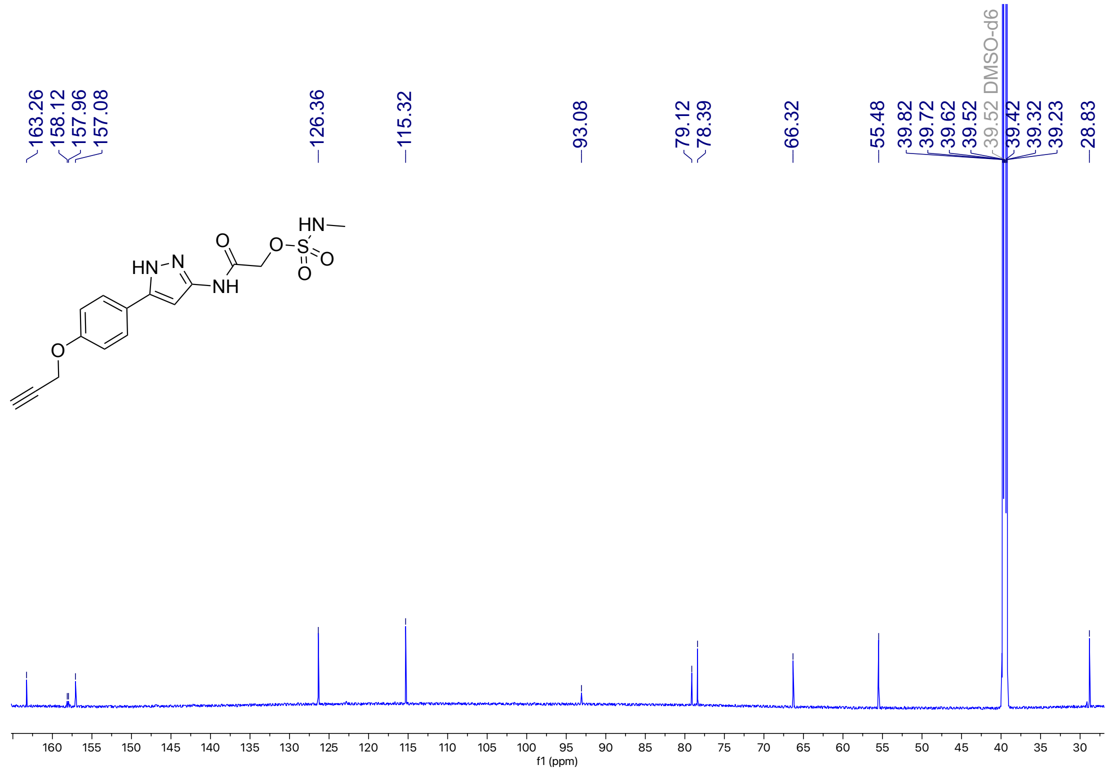

**Figure S44**: $^{1}$H NMR (400 MHz, DMSO-$d_6$) for **9c**

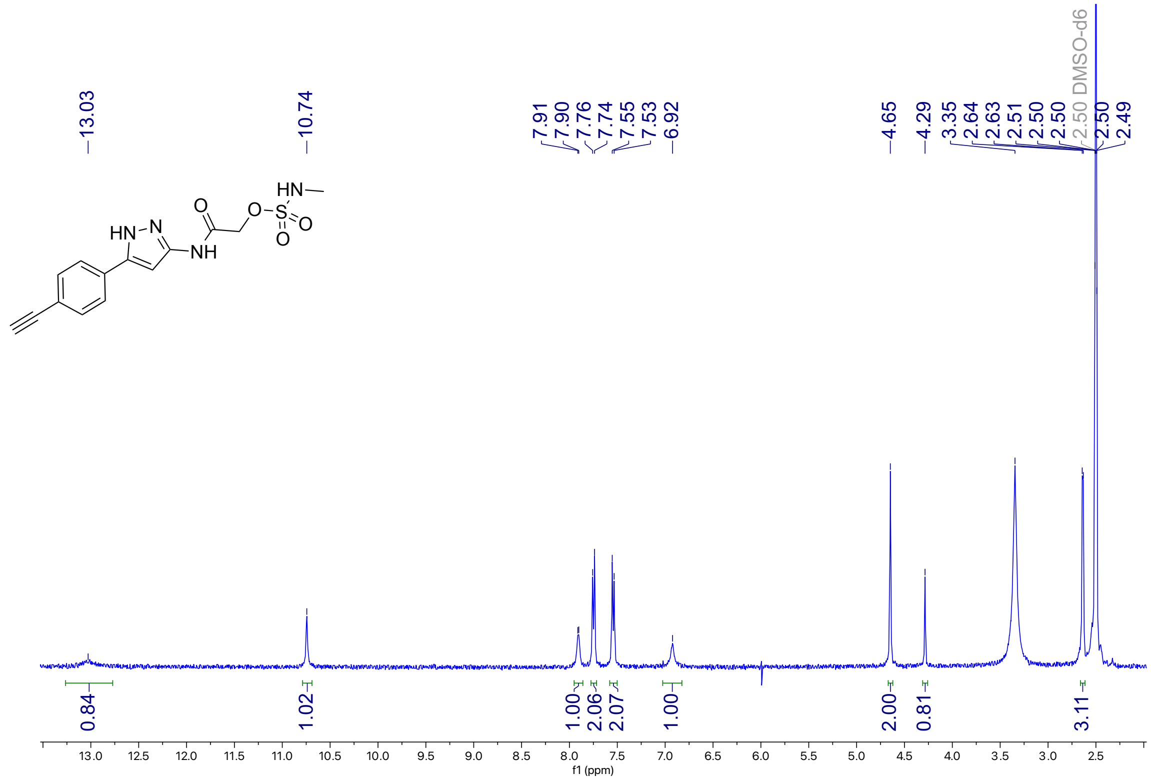

**Figure S45**: $^{13}$C NMR (214 MHz, DMSO-$d_6$) for **9c**

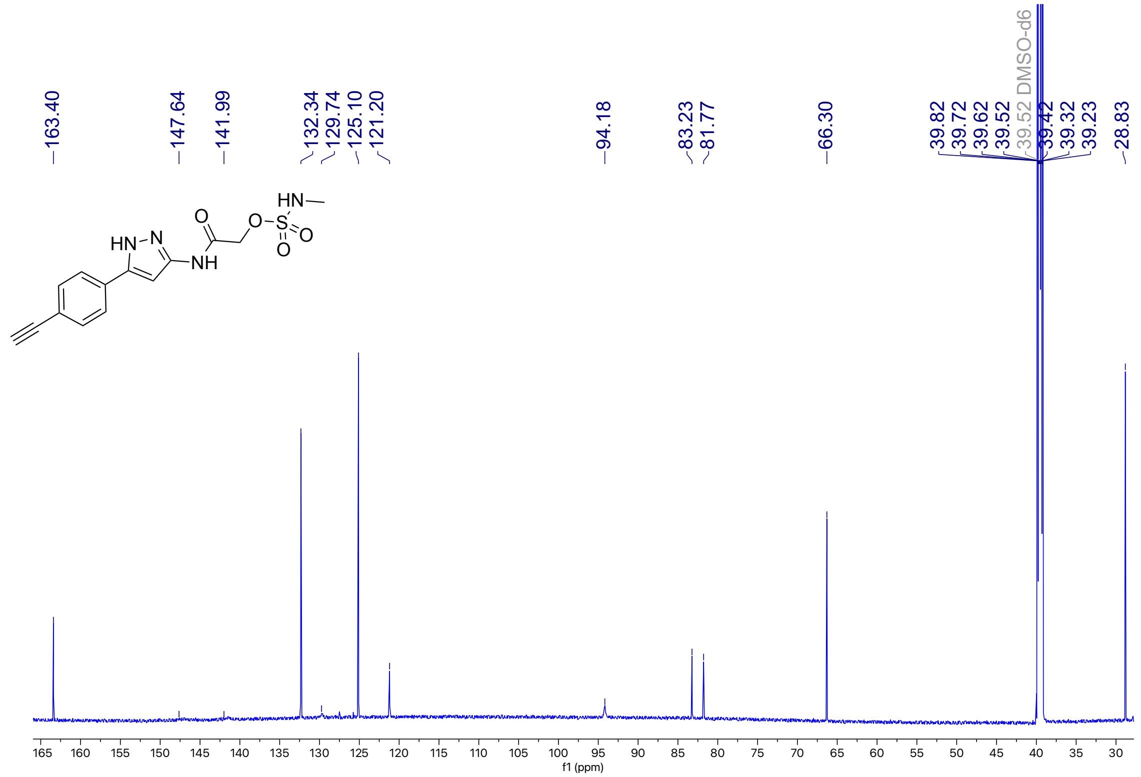

### HPLC analyses

**Figure S46**. HPLC trace of **1**

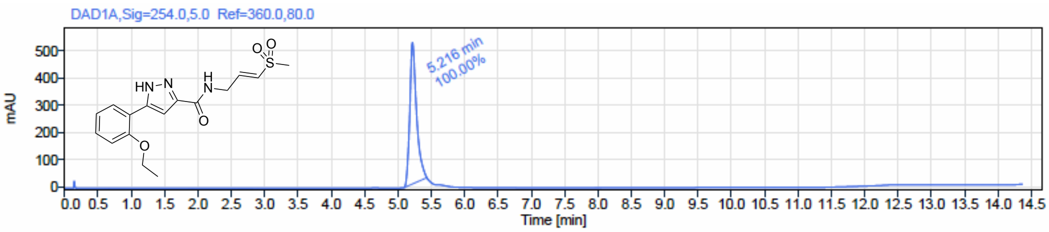


Signal Name DAD1A

| RT (min) | Signal description | m/z | Purity UV(‰) / MS(%) | Area | Area% |
|---|---|---|---|---|---|
| 5.216 | DAD1A,Sig=254.0,5.0 Ref=360.0,80.0 | | | 3652.8 | 100.00 |

**Figure S47**. HPLC trace of **2**

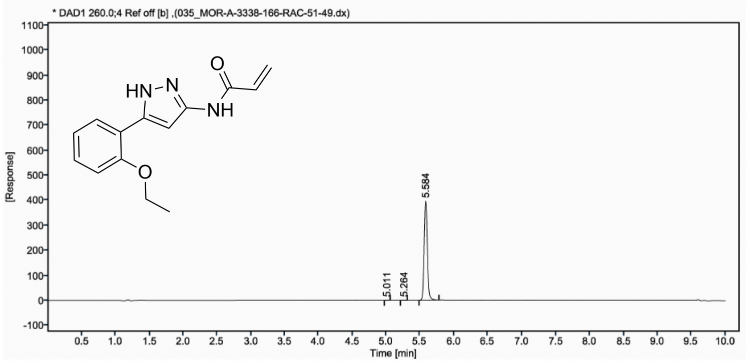


| Sr.No. | RT | Area | Height | % Area |
|---|---|---|---|---|
| 1 | 5.01 | 2 | 1 | 0.16 |
| 2 | 5.26 | 2 | 1 | 0.15 |
| 3 | 5.58 | 1286 | 395 | 99.69 |

**Figure S48**. HPLC trace of **3**

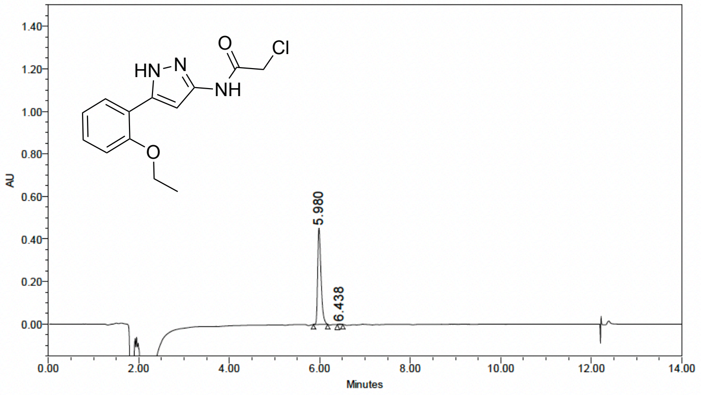


Peak Results

| | Name | RT | Height (µV) | Area (µV*sec) | % Area |
|---|---|---|---|---|---|
| 1 | | 5.980 | 452570 | 2320222 | 99.49 |
| 2 | | 6.438 | 2795 | 11856 | 0.51 |

**Figure S49**. HPLC trace of **4**

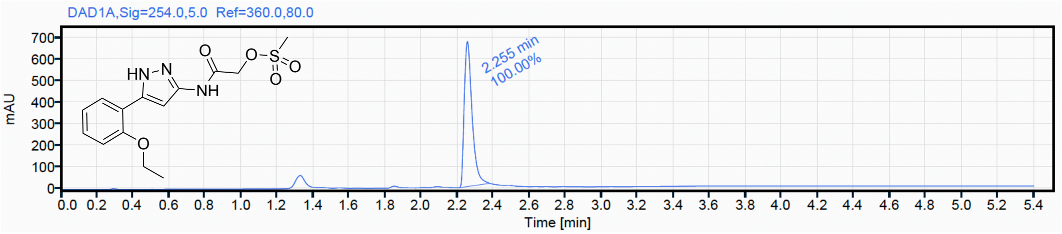


| Signal Name | DAD1A | | |
|---|---|---|---|
| **RT (min)** | **Signal description** | **Area** | **Area%** |
| 2.255 | DAD1A,Sig=254.0,5.0 Ref=360.0,80.0 | 2044.2 | 100.00 |

**Figure S50**. HPLC trace of **5**

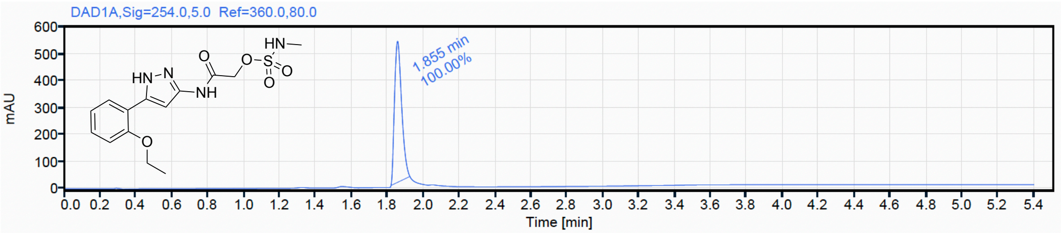


| Signal Name | DAD1A | | |
|---|---|---|---|
| **RT (min)** | **Signal description** | **Area** | **Area%** |
| 1.855 | DAD1A,Sig=254.0,5.0 Ref=360.0,80.0 | 1446.9 | 100.00 |

**Figure S51**. HPLC trace of **6a**

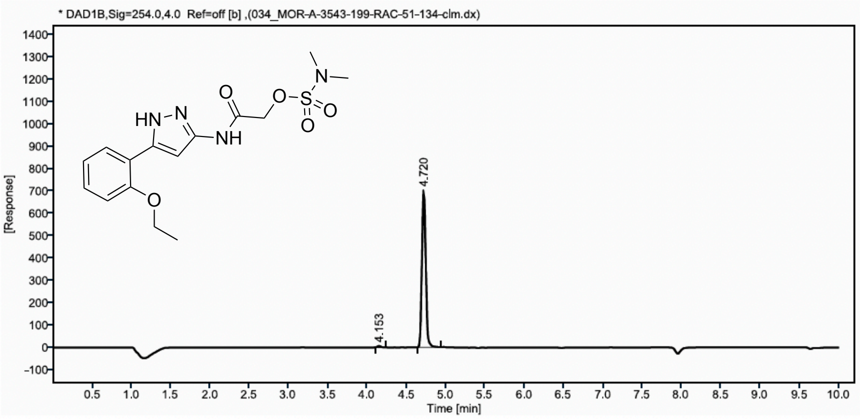


| Sr,No. | RT | Area | Height | % Area |
|---|---|---|---|---|
| 1 | 4,15 | 18 | 6 | 0,76 |
| 2 | 4,72 | 2404 | 704 | 99,24 |

**Figure S52**. HPLC trace of **6b**

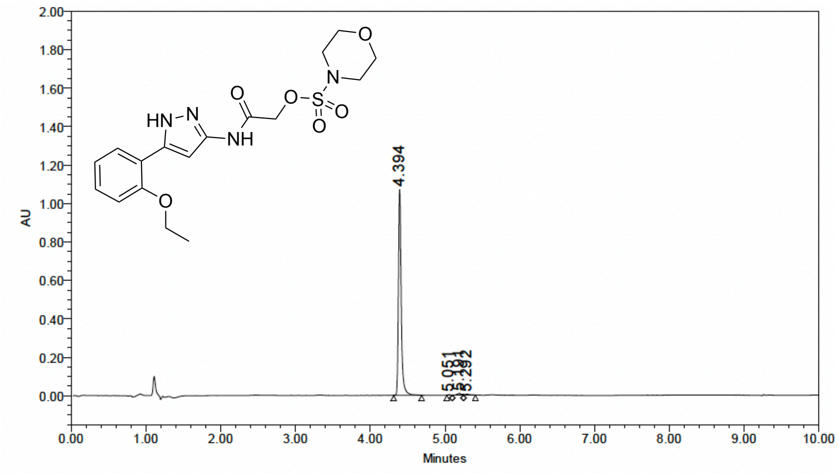


**Peak Results**

| | Name | RT | Height (μV) | Area (μV*sec) | % Area |
|---|---|---|---|---|---|
| 1 | | 4.394 | 1072091 | 2779288 | 97.84 |
| 2 | | 5.051 | 2777 | 5985 | 0.21 |
| 3 | | 5.191 | 7935 | 33688 | 1.19 |
| 4 | | 5.292 | 4552 | 21689 | 0.76 |

**Figure S53**. HPLC trace of **6c**

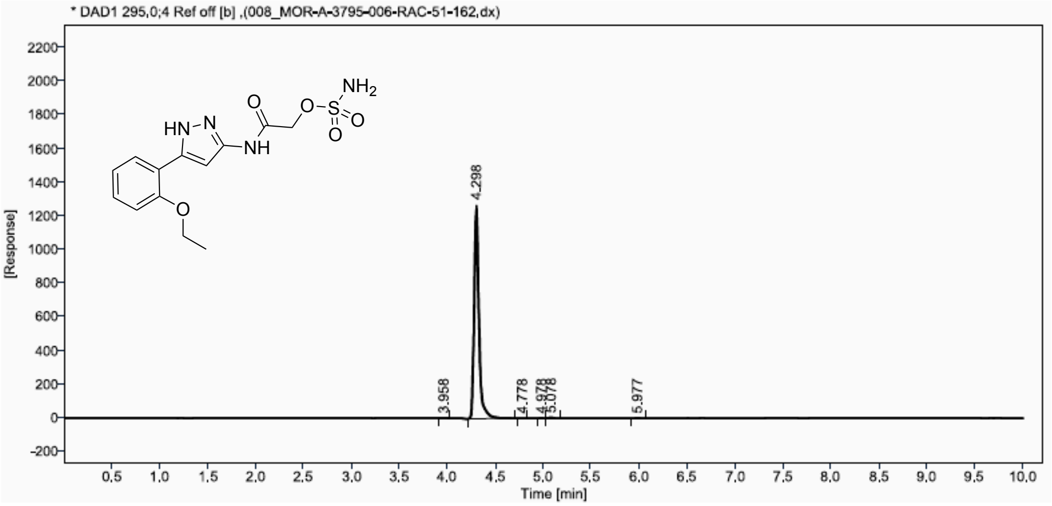


| Sr.No. | RT | Area | Height | % Area |
|---|---|---|---|---|
| 1 | 3,96 | 3 | 1 | 0,07 |
| 2 | 4,30 | 4445 | 1275 | 99,38 |
| 3 | 4,78 | 1 | 0 | 0.03 |
| 4 | 4,98 | 4 | 1 | 0,09 |
| 5 | 5,08 | 15 | 4 | 0.33 |
| 6 | 5,98 | 4 | 1 | 0,10 |

**Figure S54**. HPLC trace of **6d**

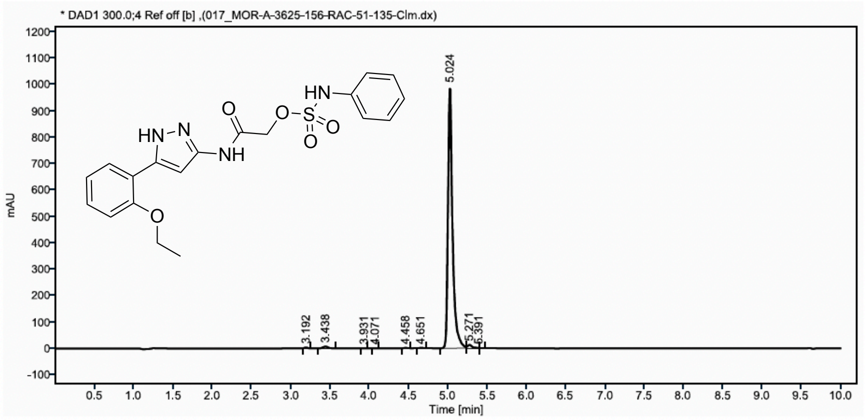


| Sr.No. | RT | Area | Height | % Area |
|---|---|---|---|---|
| 1 | 3.19 | 9 | 3 | 0.23 |
| 2 | 3.44 | 29 | 7 | 0.71 |
| 3 | 3.93 | 1 | 0 | 0.03 |
| 4 | 4.07 | 1 | 0 | 0.02 |
| 5 | 4.46 | 4 | 1 | 0.10 |
| 6 | 4.65 | 3 | 1 | 0.08 |
| 7 | 5.02 | 4004 | 993 | 97.56 |
| 8 | 5.27 | 51 | 13 | 1.25 |
| 9 | 5.39 | 1 | 0 | 0.02 |

**Figure S55**. HPLC trace of **6e**

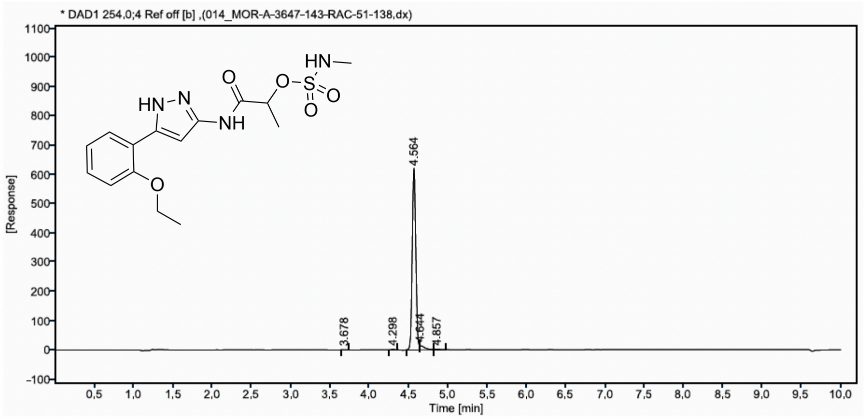


| Sr.No. | RT | Area | Height | % Area |
|---|---|---|---|---|
| 1 | 3.68 | 1 | 0 | 0.06 |
| 2 | 4.30 | 6 | 2 | 0.28 |
| 3 | 4.56 | 1966 | 617 | 96.23 |
| 4 | 4.64 | 61 | 14 | 2.97 |
| 5 | 4.86 | 9 | 2 | 0.46 |

**Figure S56**. HPLC trace of **6f**

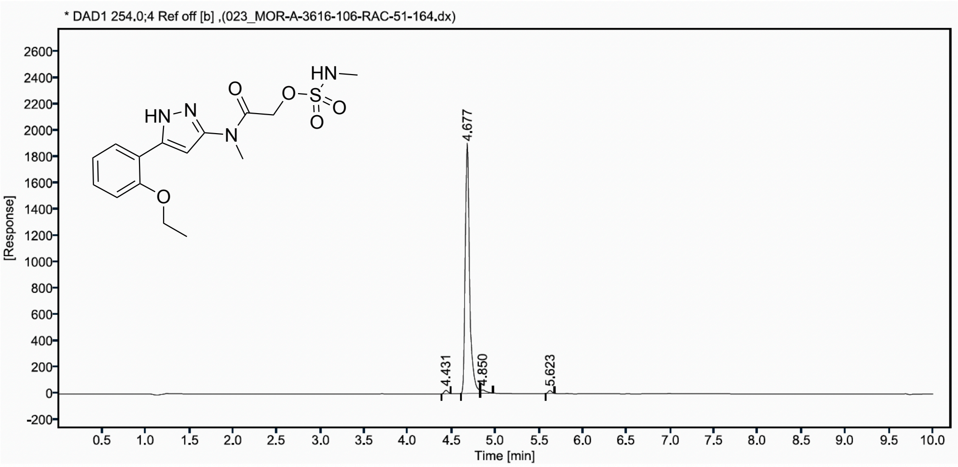


| Sr.No. | RT | Area | Height | % Area |
|---|---|---|---|---|
| 1 | 4.43 | 75 | 26 | 1.08 |
| 2 | 4.68 | 6649 | 1883 | 96.41 |
| 3 | 4.85 | 107 | 25 | 1.55 |
| 4 | 5.62 | 66 | 22 | 0.95 |

**Figure S57**. HPLC trace of **7a**

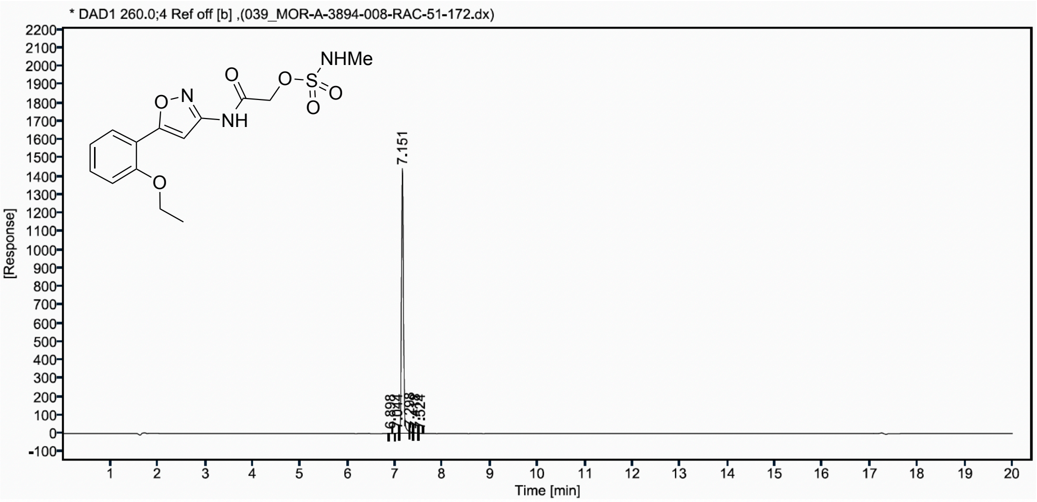


| Sr.No. | RT | Area | Height | % Area |
|---|---|---|---|---|
| 1 | 6.90 | 3 | 1 | 0.07 |
| 2 | 7.04 | 10 | 4 | 0.23 |
| 3 | 7.15 | 4320 | 1439 | 98.26 |
| 4 | 7.30 | 35 | 11 | 0.79 |
| 5 | 7.42 | 21 | 4 | 0.49 |
| 6 | 7.52 | 7 | 2 | 0.16 |

**Figure S58**. HPLC trace of **7b**

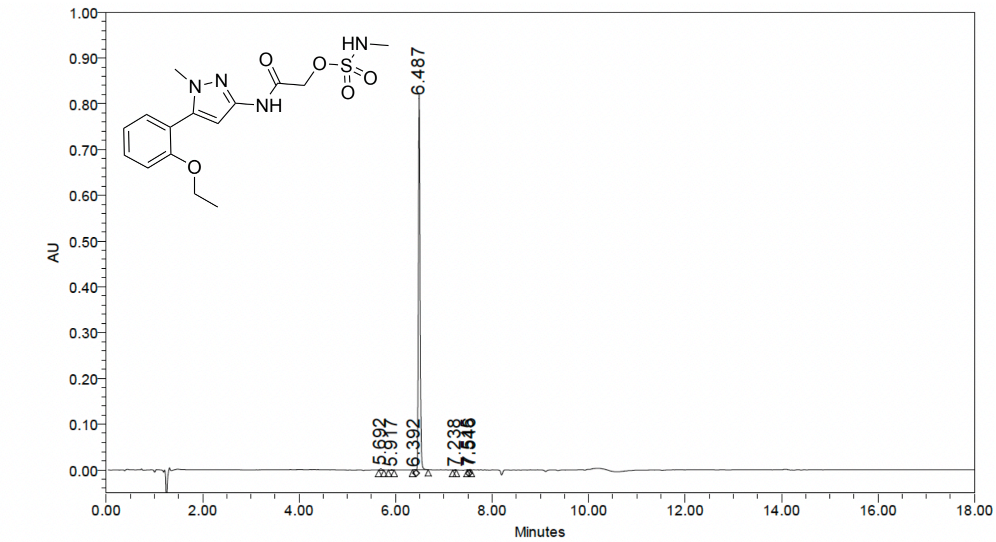


**Peak Results**

| | RT | Area (µV*sec) | Height (µV) | % Area |
|---|---|---|---|---|
| 1 | 5.692 | 5958 | 2519 | 0.29 |
| 2 | 5.917 | 1347 | 579 | 0.07 |
| 3 | 6.392 | 1426 | 658 | 0.07 |
| 4 | 6.487 | 2018110 | 820961 | 99.35 |
| 5 | 7.238 | 1630 | 1009 | 0.08 |
| 6 | 7.515 | 772 | 524 | 0.04 |
| 7 | 7.546 | 2007 | 1096 | 0.10 |

**Figure S59**. HPLC trace of **7c**

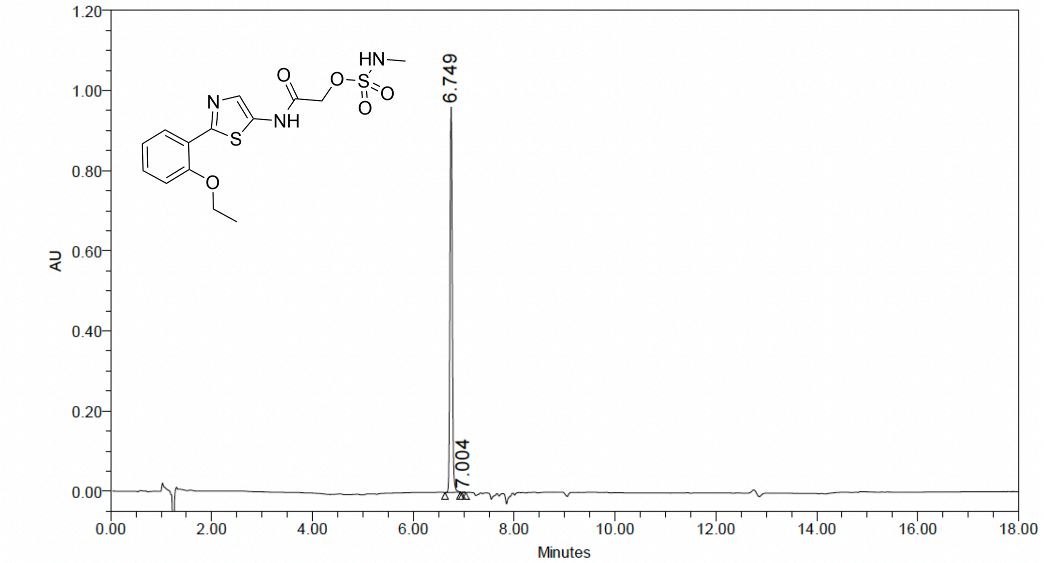


**Peak Results**

| | RT | Area (µV*sec) | Height (µV) | % Area |
|---|---|---|---|---|
| 1 | 6.749 | 3132122 | 962202 | 99.94 |
| 2 | 7.004 | 2011 | 751 | 0.06 |

**Figure S60**. HPLC trace of **7d**

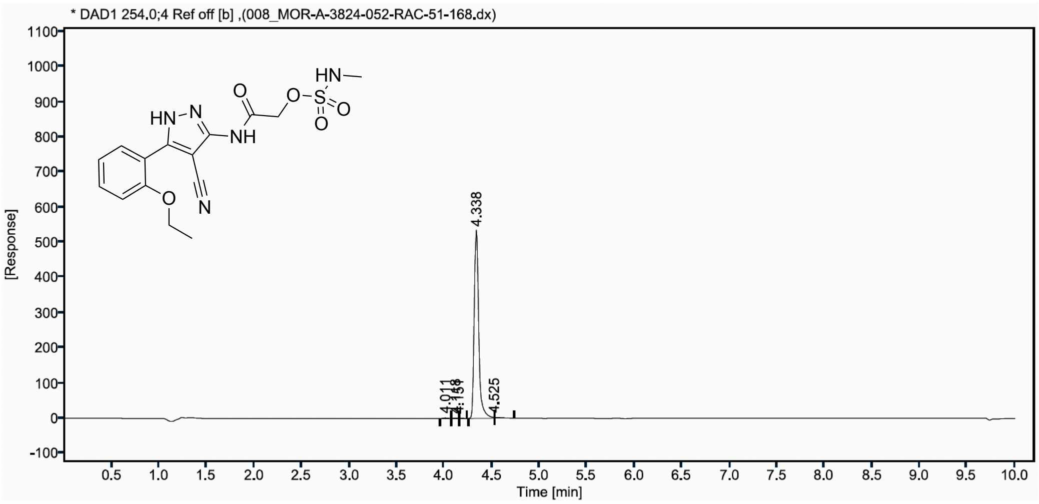


| Sr.No. | RT | Area | Height | % Area |
|---|---|---|---|---|
| 1 | 4.01 | 5 | 2 | 0.30 |
| 2 | 4.12 | 3 | 1 | 0.18 |
| 3 | 4.15 | 2 | 1 | 0.09 |
| 4 | 4.34 | 1775 | 533 | 98.60 |
| 5 | 4.52 | 15 | 3 | 0.83 |

**Figure S61**. HPLC trace of **8a**

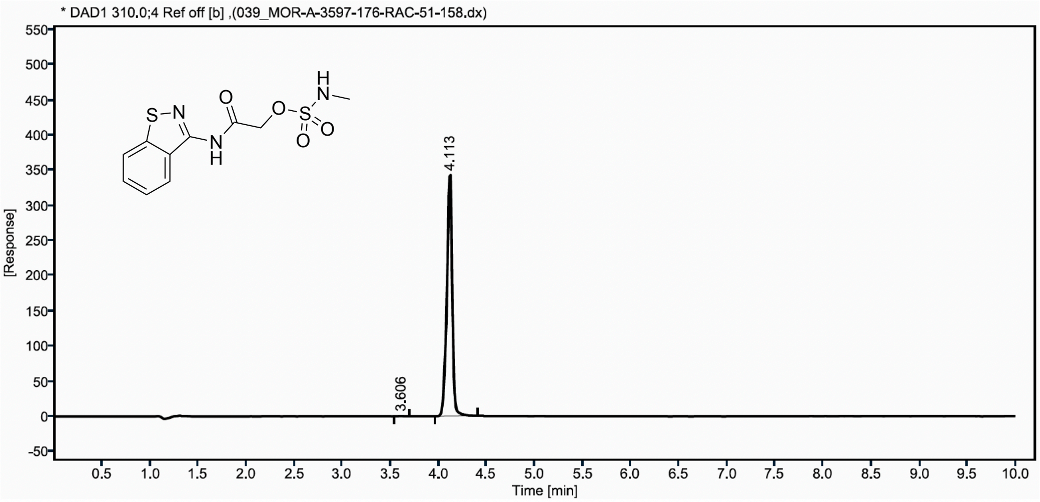


| Sr.No. | RT | Area | Height | % Area |
|---|---|---|---|---|
| 1 | 3.61 | 2 | 0 | 0.14 |
| 2 | 4.11 | 1337 | 342 | 99.86 |

**Figure S62**. HPLC trace of **8b**

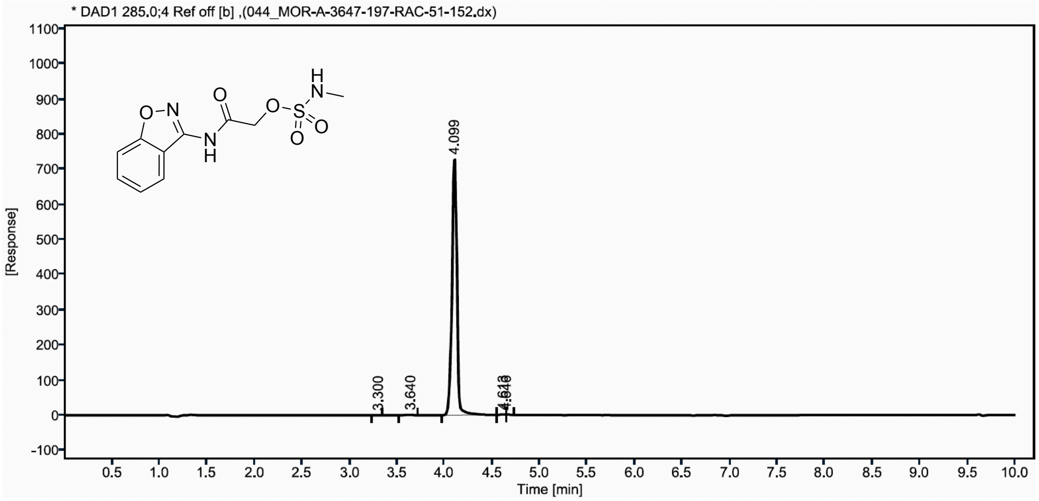


| Sr.No. | RT | Area | Height | % Area |
|---|---|---|---|---|
| 1 | 3.30 | 1 | 0 | 0.05 |
| 2 | 3.64 | 9 | 2 | 0.34 |
| 3 | 4.10 | 2663 | 729 | 99.35 |
| 4 | 4.61 | 5 | 1 | 0.17 |
| 5 | 4.65 | 3 | 1 | 0.09 |

**Figure S63**. HPLC trace of **8c**

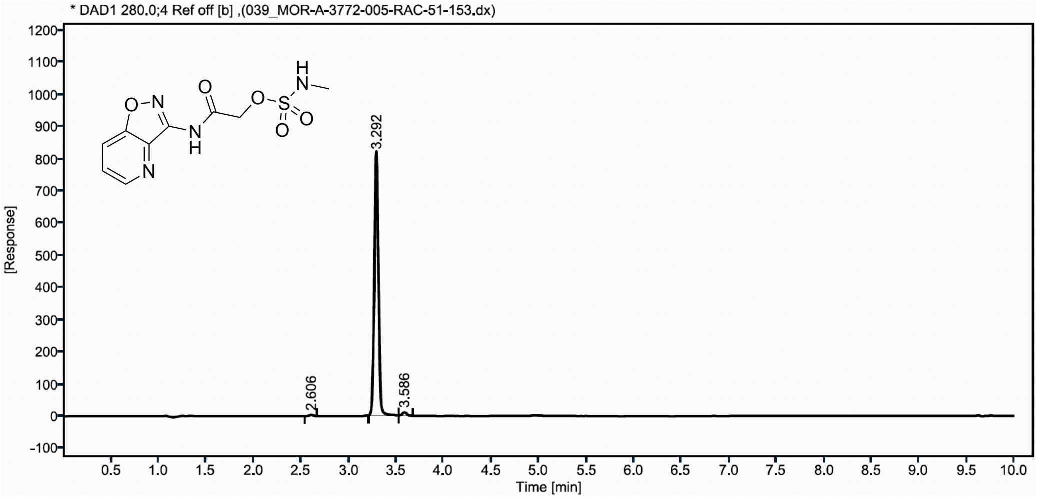


| Sr.No. | RT | Area | Height | % Area |
|---|---|---|---|---|
| 1 | 2.61 | 9 | 3 | 0.35 |
| 2 | 3.29 | 2503 | 813 | 98.36 |
| 3 | 3.59 | 33 | 11 | 1.28 |

**Figure S64**. HPLC trace of **9a**

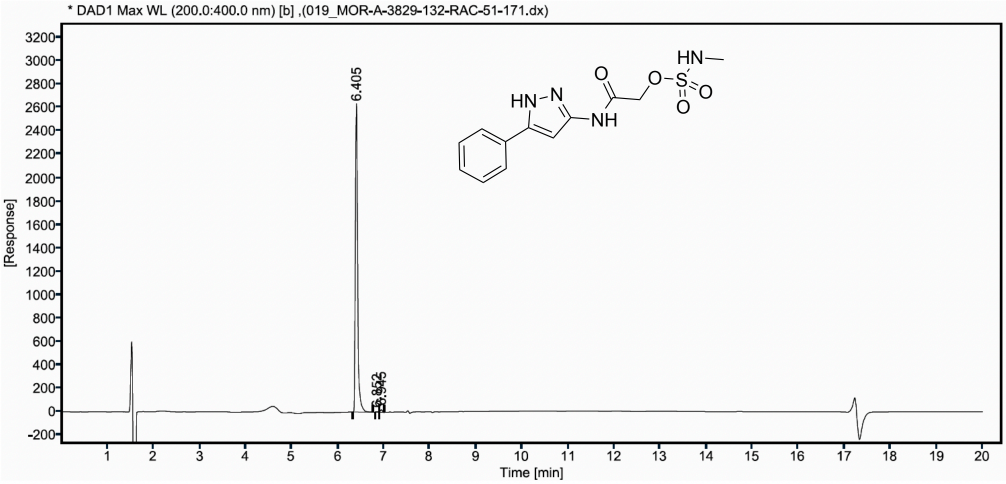


| Sr.No. | RT | Area | Height | % Area |
|---|---|---|---|---|
| 1 | 6.41 | 8877 | 2622 | 99.17 |
| 2 | 6.85 | 9 | 3 | 0.10 |
| 3 | 6.94 | 65 | 22 | 0.72 |

**Figure S65**. HPLC trace of **9b**

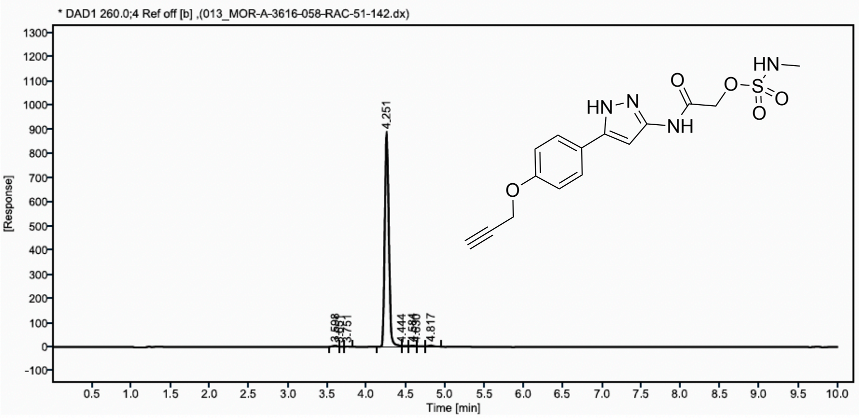


| Sr.No. | RT | Area | Height | % Area |
|---|---|---|---|---|
| 1 | 3.60 | 19 | 5 | 0.58 |
| 2 | 3.65 | 1 | 1 | 0.04 |
| 3 | 3.75 | 4 | 1 | 0.11 |
| 4 | 4.25 | 3194 | 886 | 97.20 |
| 5 | 4.44 | 13 | 3 | 0.39 |
| 6 | 4.58 | 21 | 5 | 0.63 |
| 7 | 4.63 | 11 | 2 | 0.35 |
| 8 | 4.82 | 23 | 5 | 0.71 |

**Figure S66**. HPLC trace of **9c**

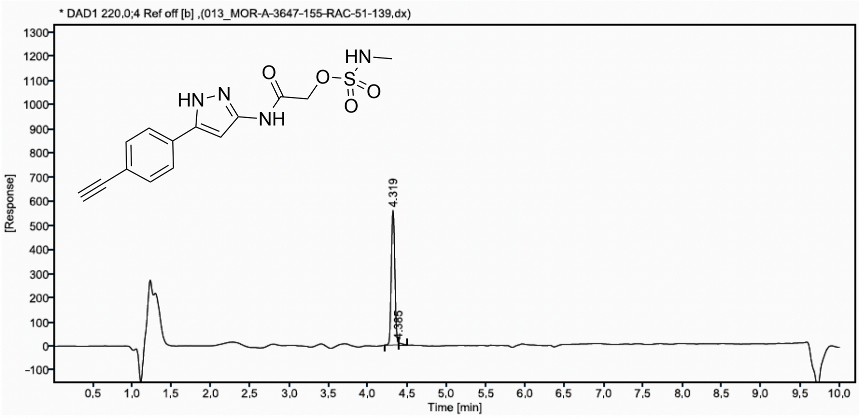


| Sr.No. | RT | Area | Height | % Area |
|---|---|---|---|---|
| 1 | 4.32 | 1723 | 556 | 98.52 |
| 2 | 4.39 | 26 | 8 | 1.48 |
